# Microenvironmental Determinants of Reaction Kinetics in Biomolecular Condensates Probed with Protein Ligation

**DOI:** 10.64898/2026.03.26.714449

**Authors:** Jangwon Bae, Kibeom Hong, Donghyuk Lee, Jaebum Jun, Yongwon Jung

## Abstract

Cells utilize liquid–liquid phase separation to organize biochemical reactions within biomolecular condensates, which function as membraneless organelles. Although these assemblies are known to enhance reaction rates by concentrating reactants, the mechanisms beyond simple mass-action effects remain poorly understood. Here, we examined how the physicochemical microenvironment within condensates modulates reaction kinetics using spontaneous protein ligation as a model reaction, conducting a systematic analysis across various condensates, ranging from structured scaffolds (PRM–SH3 systems) to intrinsically disordered protein (IDP)-based scaffolds such as LAF, TAF, and FUS. We designed a FRET-based proximity-sensitive client probe to quantify increases in effective local concentration arising from excluded-volume effects. In parallel, we measured internal hydrophilicity and water activity, revealing them as additional key determinants of reaction acceleration. Together, the findings presented here elucidate how phase-separated compartments regulate biochemical reactions through the interplay of physical (effective concentration) and chemical (hydrophilicity and water activity) microenvironments and provide mechanistic insights for engineering condensates with tunable reactivity.

## Introduction

Biomolecular condensates, formed through liquid–liquid phase separation (LLPS), have emerged as central organizational structures in cells, enabling the compartmentalization of biochemical reactions without membrane boundaries.^1–3^ These dynamic assemblies can accelerate reaction rates by concentrating reactants, a phenomenon attributed to the law of mass action. However, accumulating evidence suggests that biochemical processes within condensates are governed by mechanisms that extend beyond simple mass-action concentration effects. Proposed modulating factors include altered diffusion, extended dwell times, macromolecular confinement, specific molecular organization, and shifts in physicochemical environments.^3–6^ Despite growing interest, the mechanistic basis of these factors remains poorly understood and experimentally underexplored, mainly due to the lack of quantitative and controllable model systems. In addition, disentangling the individual contributions of these physicochemical factors to reaction kinetics proves challenging, given the complexity and interdependence of properties inherent to native cellular condensates. Gaining mechanistic insights into how condensates modulate biochemical reactions is essential not only for elucidating their physiological roles but also for leveraging these principles in synthetic biology and biocatalysis.^7–12^

Recent investigations have demonstrated that biochemical reactions within condensates can be either accelerated or suppressed relative to bulk solutions with identical enzyme and substrate concentrations. For instance, condensates can modulate enzymatic activity by altering molecular proximity and spatial organization, thereby either facilitating or impeding reaction kinetics.^10^ Additionally, ribozymes highly concentrated within peptide-based coacervates have exhibited reduced catalytic rates due to diffusion limitations arising from molecular crowding.^11^ Similarly, reaction rates often decline during condensate aging, a process accompanied by material maturation and restricted mass transport.^12^ The characterization of macroscopic condensate properties, particularly translational diffusion probed by fluorescence recovery after photobleaching (FRAP), is well established. In contrast, a significant knowledge gap exists regarding how microscopic characteristics govern reaction kinetics within these dense phases.

In this study, we aimed to identify the microenvironmental determinants of reaction kinetics within condensates, distinguishing factors that extend beyond simple mass-action effects. We established a model system utilizing the well-characterized SpyTag–SpyCatcher (ST–SC) protein ligation^13^ as an example of specific biomolecular binding analogous to natural biological regulatory functions. This platform enables precise quantification of reaction kinetics in diverse protein-based condensates. To systematically investigate how the physicochemical properties of condensates influence reaction kinetics, we examined structured protein scaffolds (PRM–SH3 system) and intrinsically disordered protein (IDP)-based scaffolds. Among the IDPs, LAF, TAF, and FUS, which differ in charge density, were analyzed to assess how sequence-dependent features modulate reactivity. Furthermore, our model permits flexible adjustment of the relative proportions between structured and IDP-based scaffolds, reflecting the multicomponent nature of biological condensates. Condensate microenvironments can also be tuned within the same scaffold by varying external conditions, such as ionic strength.

To achieve a comprehensive interpretation of the internal condensate environment, we expanded our analytical scope beyond compositional descriptors alone. First, we designed a fluorescence resonance energy transfer (FRET)-based client pair to measure the effective local client concentration within condensates. In parallel, the polarity-sensitive dye PRODAN was employed to assess relative hydrophilicity. Complementing these analyses, conventional characterizations, such as FRAP and scaffold density measurements, were performed to validate internal molecular dynamics and packing states. This multifaceted approach enables a precise and granular dissection of how the condensate microenvironment functions as a tunable bioreactor. Specifically, we demonstrate that the interplay between excluded-volume effect and hydrophilicity is a dominant determinant of ST–SC reaction kinetics. Ultimately, this framework enabled us to deconvolute the specific contributions to the overall reaction rates within each condensate system.

## Results

### Design of scaffold and client systems for quantifying reaction kinetics inside condensates

To elucidate the determinants of reaction kinetics within condensates, we developed a defined *in vitro* model system comprising diverse protein scaffolds and a reaction-competent client pair (Fig. 1). We employed a metal-ion-induced phase-separation method that leverages multivalency via Ni²⁺–His-tag coordination.^14^ This approach efficiently induces LLPS of His-tagged scaffold proteins without requiring additional domains and minimizes perturbation to the system. To generate condensate environments with distinct physicochemical landscapes, we designed multiple types of phase-separating scaffolds. As a representative structured scaffold, we utilized a previously reported modular construct comprising a proline-rich motif (PRM), a Src homology 3 (SH3) domain, and linkers.^14^ The specific, reversible interaction between PRM and SH3 can drive LLPS by forming multivalent intermolecular networks.^15^ We also selected three IDP constructs: LAF (residues 1–168), TAF (residues 1–208), and FUS (residues 2–214), each exhibiting unique residue compositions and physicochemical properties. The LAF segment corresponds to the RGG domain, characterized by a relatively high content of charged residues, with arginine serving as the electrostatic key that drives LLPS.^16^ The prion-like domains of TAF^17^ and FUS^18^ used in this study are rich in serine, tyrosine, glutamine, and glycine (SYQG domain), where multiple tyrosine residues mediate weak π–π stacking interactions. While FUS is nearly devoid of charged residues and forms a hydrophobic core, TAF contains a higher proportion of ionizable residues (see Supplementary Materials Section 1.1). To ensure solubility and stability, a small ubiquitin-like modifier (SUMO) tag was fused to these IDP constructs.

**Fig. 1.**
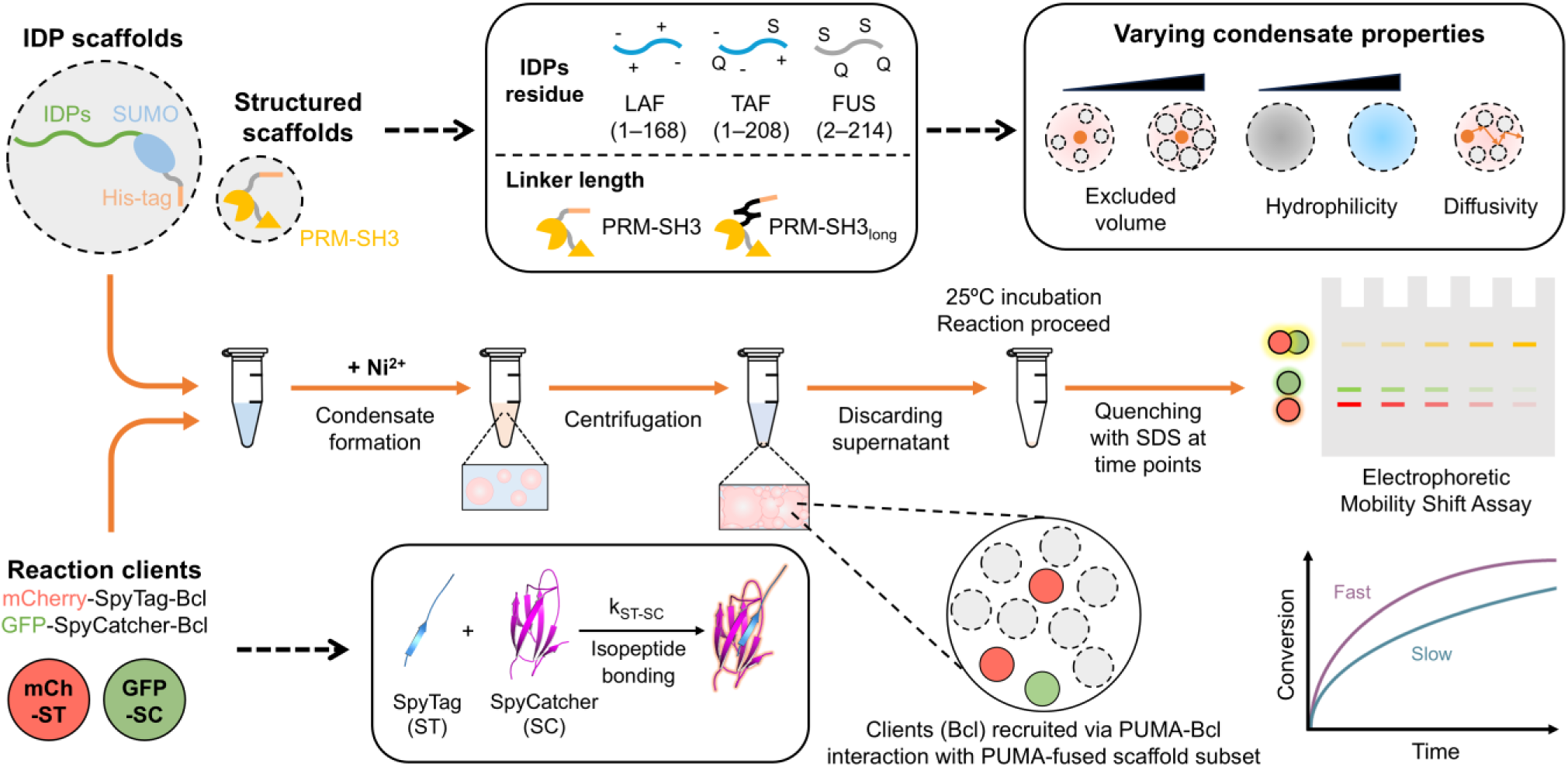
Schematic of the experimental workflow to measure reaction kinetics within biomolecular condensates. To create condensates with diverse physicochemical properties, multiple types of phase-separating scaffolds (His-tagged IDPs or PRM–SH3 modules) are employed. Phase separation is driven by the coordination between Ni^2+^ ions and His-tags on the scaffold proteins. The reaction client pair consists of mCh–ST and GFP–SC, which form a covalent bond to yield a mCh–GFP ligated product. Reaction progress is quantified using gel electrophoresis and band-shift analysis. As the reaction proceeds, the amount of the ligated product (yellow band) increases, corresponding to a decrease in the unreacted mCh–ST (red band) and GFP–SC (green band).

Fluorescent protein-fused clients, mCherry–SpyTag (mCh–ST) and GFP–SpyCatcher (GFP–SC), were constructed to quantify reactants and ligated products. These clients spontaneously form an irreversible isopeptide bond upon mixing. The reaction was quenched at defined time points by the addition of sodium dodecyl sulfate (SDS) buffer as previously described.^19^ This approach allows for the quantitative assessment of reaction progress via band-shift analysis using polyacrylamide gel electrophoresis (PAGE; Fig. 2A). The ST/SC client pair was recruited into condensates via high-affinity interaction between the PUMA peptide and the Bcl-xL protein (K_d_ ≈ 3 nM),^20–22^ which were fused to the scaffolds and clients, respectively. This recruitment mechanism is orthogonal to the metal-ion-induced LLPS process. A small fraction (1–10%, with 5% as the standard condition) of the PUMA-fused scaffold was incorporated into the total scaffold pool to avoid perturbing the intrinsic material properties of the condensates. His-tags were removed from client proteins to prevent nonspecific recruitment or self-association mediated by metal ions. After inducing phase separation, the samples were centrifuged to separate the condensate and dilute phases, and the supernatant was then discarded. Reaction kinetics within condensates were quantified by resuspending pelleted condensates in SDS-containing buffer for immediate quenching, followed by the band-shift assay (Fig. 2B).

**Fig. 2.**
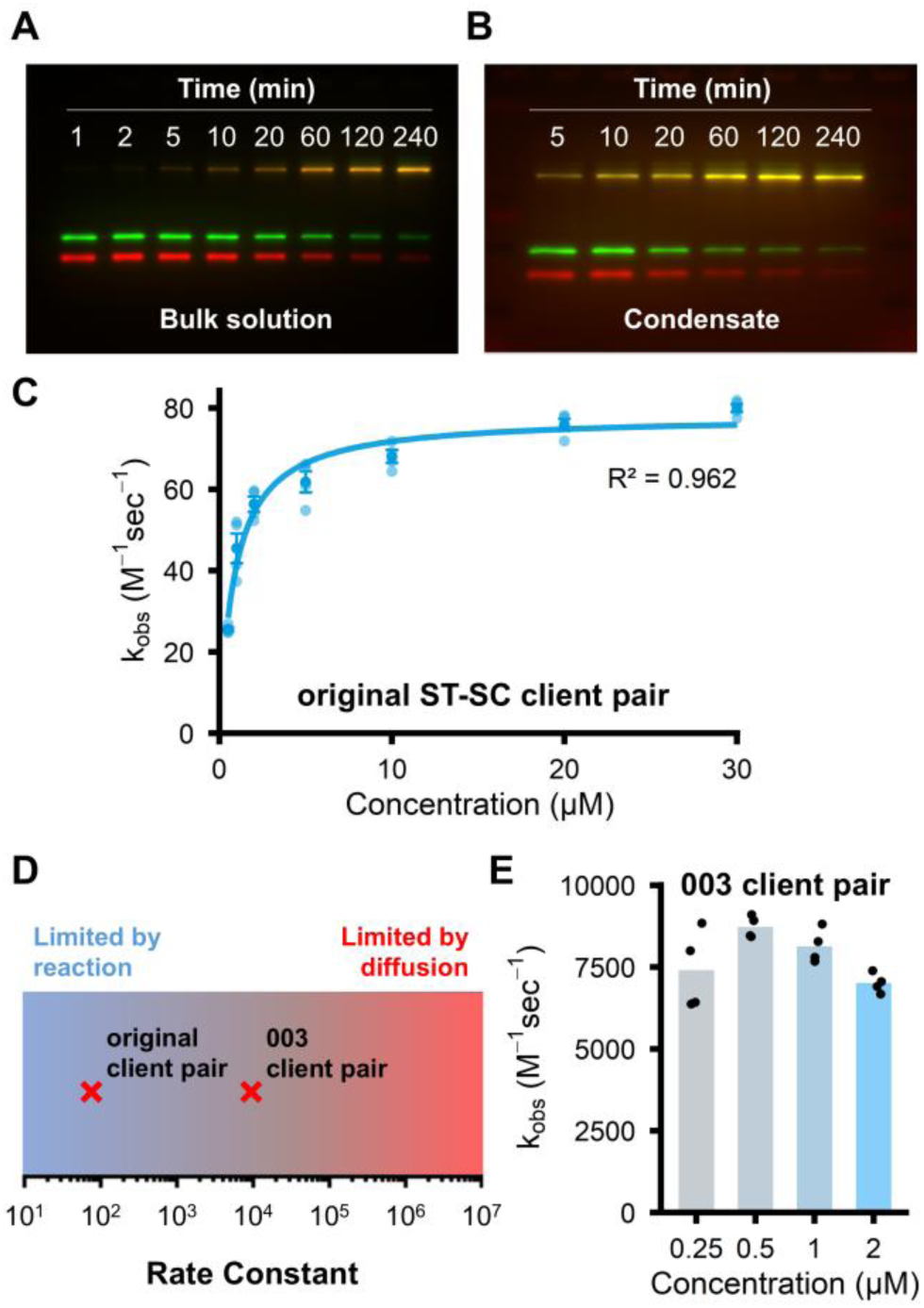
Quantification and characterization of ST–SC reaction kinetics. (**A**) A representative PAGE gel fluorescence image showing the reaction between mCh–ST (red) and GFP–SC (green) over time in a 5 µM bulk solution. (**B**) A corresponding reaction kinetics gel for the PRM–SH3 condensate system, demonstrating capture of the reaction within the condensed phase. (**C**) A plot of the apparent second-order rate constant (k_obs_) versus initial concentration for the original client pair in this study. Bold data points denote the average rate constant at each concentration, with raw data depicted as semi-transparent dots. The solid line indicates the best fit to hyperbolic kinetics. Error bars correspond to ±1 standard error (SE). (**D**) A schematic illustrating that the original ST–SC client pair (k ≈ 10¹–10² M⁻¹ s⁻¹) operates in the limited by reaction regime, while the 003 client pair (k ≈ 10⁴ M⁻¹ s⁻¹) approaches the diffusion-influenced regime. (**E**) A plot of the apparent rate constant of the 003 client pair. In all bar graphs, the bars represent the mean values, and the overlaid black dots indicate the raw data of each measurement.

### Non-diffusion-limited characteristics of ST–SC ligation

Diffusion-limited reactions are strongly influenced by diffusivity, which is often substantially lower inside condensates than in the bulk solution. The influence of internal diffusion on reaction kinetics within biomolecular condensates has been characterized using various experimental and theoretical frameworks.^11,12,23–26^ Moreover, many natural processes operate below the diffusion ceiling, with kinetics governed by intrinsic binding and activation steps rather than encounter frequency alone.^27^ Therefore, we sought to examine a non-diffusion-limited model reaction within condensates to investigate other determinants. The ST–SC reaction is reported to proceed with a rate constant of approximately 10³ M⁻¹ s⁻¹, well below the theoretical diffusion-limited regime, which typically exceeds 10⁵–10⁶ M⁻¹ s⁻¹.^13,19^ We measured the reaction rate constant of the ST–SC client pair used in our system under bulk solution conditions. The rate constant was further reduced to 10¹–10² M⁻¹ s⁻¹, indicating an additional kinetic slowdown caused by the fused protein domains (Fig. 2C). This slow reaction rate places the system entirely within the non-diffusion-limited regime (Fig. 2D), which is particularly advantageous for deconvoluting microenvironmental determinants from macroscopic diffusion. Additionally, this regime provides high temporal resolution owing to reaction timescales on the order of minutes. As shown in Fig. 2C, the apparent second-order rate constant (k_obs_) exhibited concentration dependence and followed hyperbolic kinetics, a characteristic feature of reaction-limited processes.^28,29^ In contrast, the engineered SpyTag003–SpyCatcher003 pair, which is known to approach the diffusion-limited regime,^19^ showed a nearly constant rate of about 10⁴ M⁻¹ s⁻¹ across the 0.25–2 μM range, and this reaction rate was notably decreased under viscous conditions (Fig. 2E; see Supplementary Materials Sections 1.4 and 1.5).

### Unreacted fraction and client heterogeneity under certain crowded conditions

When characterizing the ST–SC reaction under various conditions, we observed that in the presence of high concentrations of polyethylene glycol (PEG), the reaction proceeded only up to a specific extent and then ceased even after prolonged incubation (Fig. 3A). While the ST–SC reaction exhibits reduced reactivity at lower reactant concentrations, leading to deviations from simple second-order kinetics, the observed plateau cannot be explained solely by these kinetic limitations. Confocal laser scanning microscopy revealed that, at high PEG concentrations, the clients in the reaction mixture became compartmentalized (Fig. 3B). Although these client-rich phases did not exhibit fully liquid-like properties, they disappeared completely upon dilution, indicating that these phases represent a reversible state rather than irreversible aggregation. Interestingly, diluting PEG restored ST–SC reactivity, allowing the reaction to resume and proceed to completion (see Supplementary Materials Section 1.6).

**Fig. 3.**
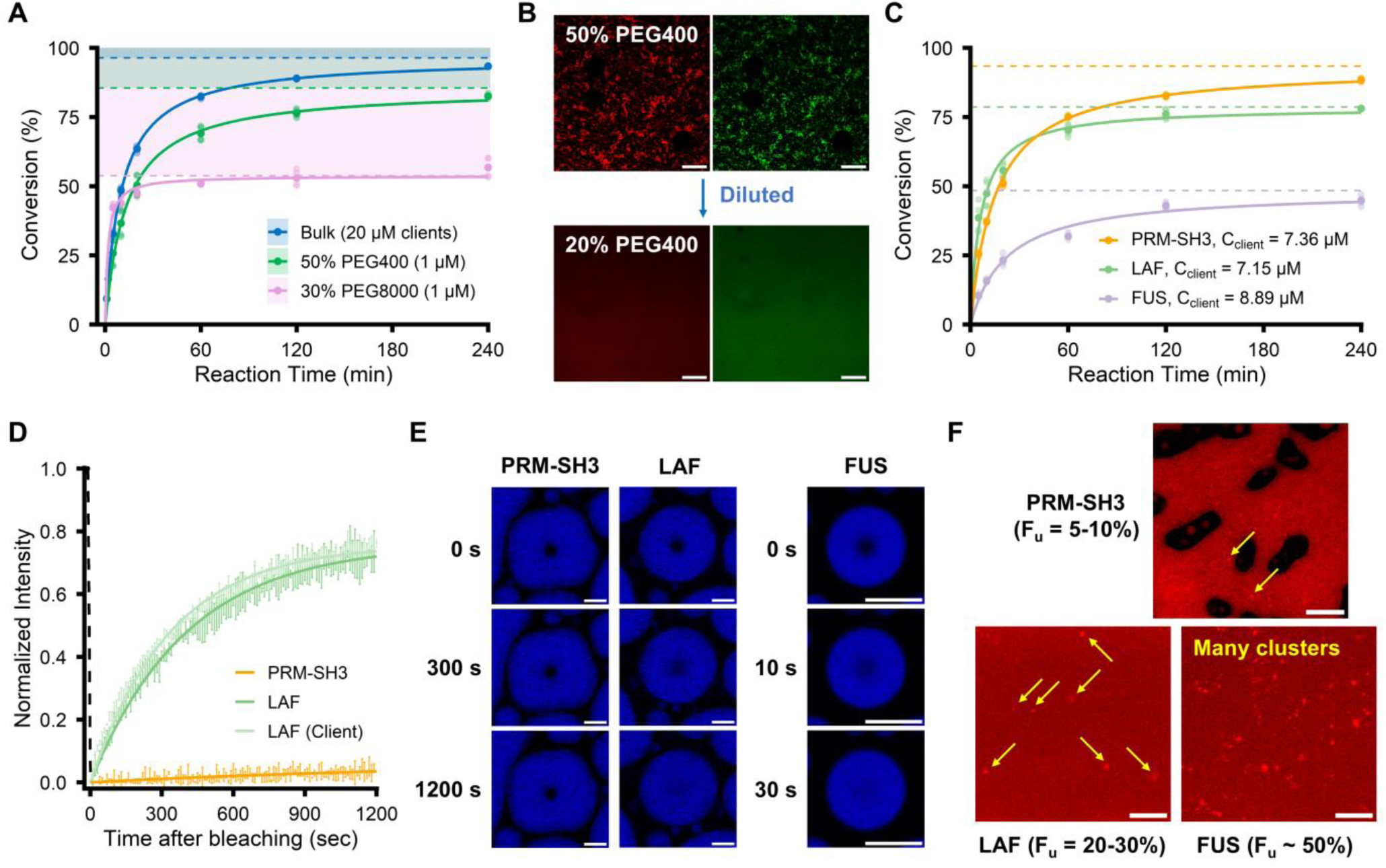
The presence of unreacted fractions and the heterogeneous clustering of client molecules under certain crowded conditions. (**A**) Reaction conversion over time in a bulk solution of 20 µM clients versus solutions with high concentrations of PEG. Bold data points in all conversion plots denote the average conversion at each time point, with raw data depicted as semi-transparent dots. The solid line indicates the best fit to modified second-order kinetics, and dashed lines represent the fitted unreacted fraction. (**B**) Confocal microscopy images of the reaction clients (mCh in red, GFP in green). In 50% PEG400, the clients associate and compartmentalize, and this phase separation is reversible upon dilution. Scale bars, 100 µm. (**C**) Reaction kinetics and unreacted fractions within three types of condensates (PRM–SH3, LAF, and FUS), where C_client_ represents the internal client concentration within the condensate phase. (**D**) Condensate FRAP curves for PRM–SH3 and LAF scaffolds, alongside the recovery of mCh client partitioned within LAF condensates. Error bars represent ±1 standard deviation (SD; N=3). (**E**) Representative FRAP images for PRM–SH3, LAF, and FUS condensates. Scale bars, 10 µm. (**F**) Confocal microscopy images showing that the heterogeneous clustering of mCh clients becomes more pronounced as the unreacted fraction increases within the condensates. Scale bars, 10 µm.

This phenomenon was similarly observed in condensates, particularly in those formed by IDP-based scaffolds. For instance, substantial unreacted fractions were observed in both the LAF (21.2%) and FUS (51.5%) condensates (Fig. 3C). These values represent the populations remaining unreacted even at long reaction times, as estimated from kinetic fitting. In contrast, the structured PRM–SH3 scaffold showed an unreacted fraction of only 6.57%. Notably, IDP condensates displayed significantly higher diffusivity and mobile fractions in FRAP analysis compared to the PRM–SH3 condensate (Fig. 3D,E). This result implies that the incomplete reaction is not caused by restricted macroscopic molecular mobility. Furthermore, the heterogeneous clustering of client molecules became more pronounced as the unreacted fraction increased in LAF and FUS condensates (Fig. 3F). Such internal heterogeneity aligns with recent studies suggesting that condensates are not simple, uniform liquid droplets.^30,31^ While client clustering under crowded conditions (high PEG and within IDP condensates) may contribute to the unreacted fraction, the mechanistic basis of this heterogeneous reaction population remains unclear.

In our analysis, we treated the unreacted fraction (F_u_) as the population unable to participate in the reaction. Therefore, it was excluded from the client concentration to model the reaction progress. Accordingly, the data were fitted to a modified second-order rate equation v=k_adj_([client]_t_-F_u_[client]_0_)^2^. Here, [client]_t_ and [client]_0_ denote the concentrations of non-ligated clients at time t and at t=0, respectively. Although this model assumes that F_u_ remains inert and constant over time, it provides a robust metric for comparing relative reaction rates across samples. Unless otherwise specified, all rate constants reported in this study refer to these adjusted values (k_adj_). The ST and SC clients were recruited into condensates at a 1:1 stoichiometric ratio, and the final rate constant was determined by averaging the values derived from the depletion kinetics of each client. The kinetics across different condensates were evaluated at internal client concentrations (C_client_) within the optimal reaction range (1–10 µM), and the rate constant measured in 5 μM bulk solution (72.98 M^-1^ s^-1^) was selected as the reference (see Supplementary Materials Section 1.4).

### Examination of excluded-volume effect by modulating condensate scaffold density

We first investigated the impact of internal density within a consistent scaffold framework by modulating the linker length positioned between the His-tag and the PRM–SH3 module as previously established (Fig. 1).^14^ Specifically, we compared the original PRM–SH3 scaffold with a long-linker counterpart (PRM–SH3_long_) featuring a 48-residue rigid linker composed of an α-helical (EAAAK)_8_ repeat. We observed that the PRM–SH3 scaffold formed significantly denser condensates and exhibited a higher reaction rate constant than the long-linker variant (Fig. 4A,B). This correlation suggests that the denser environment of the original scaffold imposes more considerable excluded-volume effects and accelerates reaction kinetics.

**Fig. 4.**
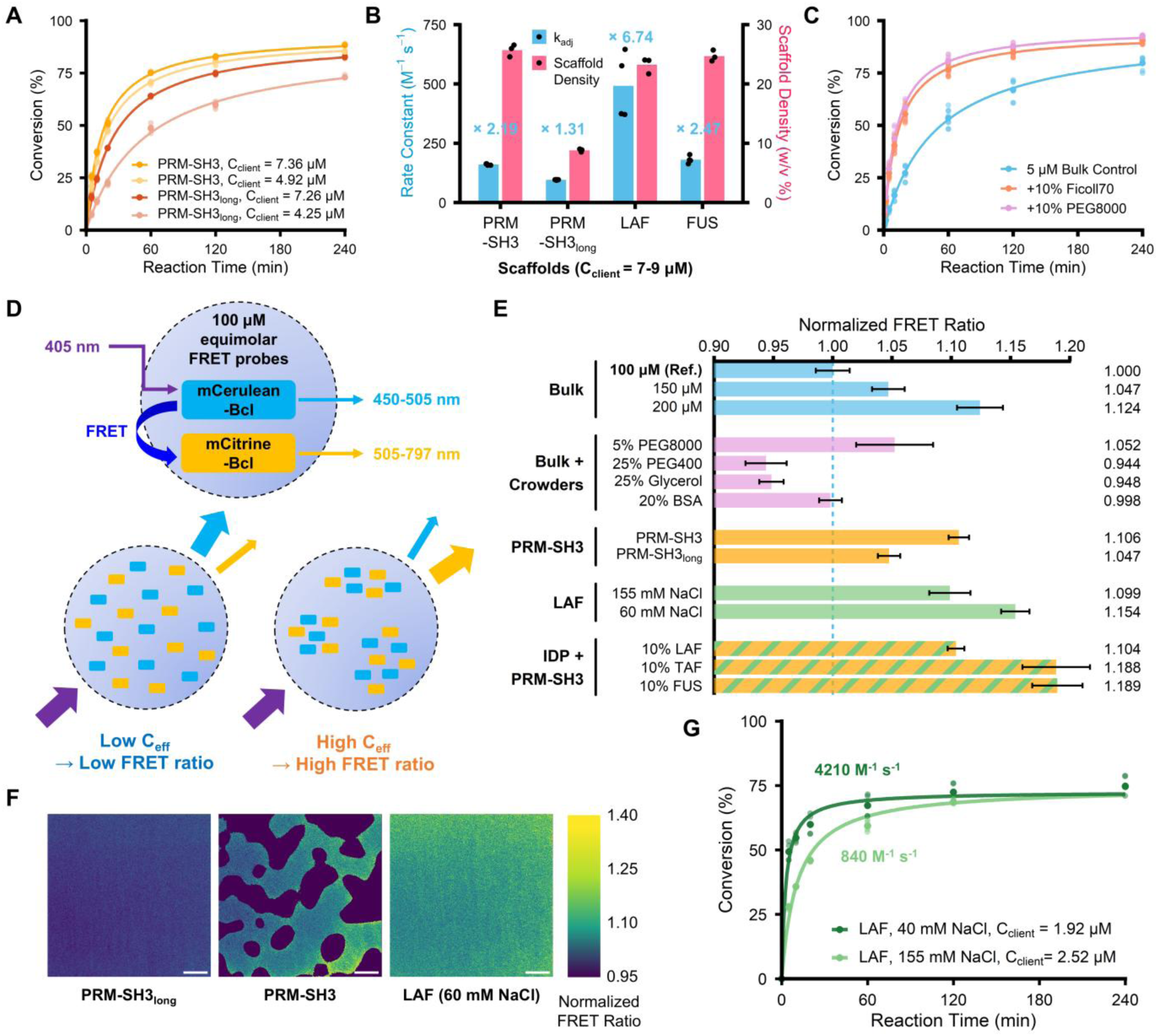
A designed FRET probe system demonstrating that increased effective concentration enhances reaction rates in condensates. (**A**) Reaction conversion over time for various PRM–SH3 scaffold conditions. (**B**) Analysis of rate constants and internal scaffold density for different condensates. The fold-increase values (e.g., ×2.19) indicated above the bars denote the rate enhancement relative to the 5 µM bulk solution. All internal client concentrations within condensates in this plot fall within the range of 7–9 µM. (**C**) Comparison of reaction conversion over time in bulk solution versus crowded environments containing 10% (w/v) Ficoll70 or PEG8000. (**D**) Schematic representation of the FRET-based probe system used to directly measure relative effective client concentration (C_eff_). (**E**) Comparison of normalized FRET ratios across various bulk, crowder, and condensate conditions (referenced to 100 µM bulk solution of FRET probes = 1.00). Error bars represent ±1 SD (N=5). (**F**) Normalized FRET ratio heatmaps illustrating effective client concentration within PRM–SH3_long_, PRM–SH3, and LAF condensates. Scale bars, 20 µm. (**G**) Reaction conversion kinetics of the LAF condensate measured under the standard condition (155 mM NaCl) and a lower-salt condition (40 mM NaCl).

Macromolecularly crowded environments are known to exert excluded-volume effects, slow molecular diffusion, and induce conformational changes in macromolecules.^32–35^ Focusing specifically on the excluded-volume effect, macromolecules occupy a substantial fraction of the total volume, which confines reactants to a reduced free volume. This confinement effectively increases the local reactant concentration and enhances the probability of molecular encounters. To investigate the role of effective concentration driven by these excluded-volume effects, we examined the ST–SC reaction in a bulk solution containing the macromolecular crowding agents PEG8000 or Ficoll70. As expected, the addition of these agents accelerated the reaction rate (Fig. 4C). This result supports the general notion that increased molecular encounter frequency, driven by excluded-volume effects, enhances reactivity. However, while these bulk assays qualitatively support the proposed mechanism, quantitative determination of the effective client concentration within condensates is required to assess its contribution to reaction kinetics.

### Quantification of effective concentration using a designed FRET client pair

To quantitatively assess whether clients experience a closer proximity and, consequently, a higher effective concentration within condensates than in the bulk solution, we implemented a FRET-based readout tailored for direct comparison across environments. While prior FRET sensors for macromolecular crowding have been reported,^36^ they typically employ a single-chain design in which the donor and acceptor are covalently linked. These sensors respond to crowding-induced conformational changes, but because crowding is inherently a phenomenological measure, it often aggregates multiple physical contributions into a single response. In contrast, our objective was to measure effective concentration, which provides a direct, quantifiable physical measure of the molecular density within condensates. To achieve this, we employed a discretized FRET pair strategy where the mCerulean donor and mCitrine acceptor were expressed as separate Bcl protein fusions rather than being linked (Fig. 4D). Because these probes are physically separated, a FRET signal occurs when the high effective concentration of the clients brings them into close proximity, approaching the Förster radius. These probes were introduced at equimolar ratios at the highest practical concentration of 100 μM for each component. By exciting the donor, we measured the FRET ratio, defined as the ratio of acceptor emission (505–797 nm) to donor emission (450–505 nm), which serves as a proxy for the local effective concentration.

We first validated the probe response in a bulk solution to establish a reference baseline. When the FRET ratio in the 100 μM bulk solution of the probes was normalized to 1.00, it increased by 4.7% at 150 μM and by 12.4% at 200 μM (Fig. 4E). This result confirms that increasing client concentration reduces average intermolecular distances, which enhances FRET efficiency. Notably, adding 5% (w/v) PEG8000 to a 100 μM bulk solution increased the FRET ratio by 5.2%, comparable to that at 150 μM. Assuming spherical geometry for PEG8000, the excluded volume fraction at 5% (w/v) was calculated to be 38.7%, resulting in a 1.63-fold increase in effective concentration (see Supplementary Materials Section 2). In contrast, small cosolutes such as PEG400 or glycerol impart negligible excluded volume and primarily increase viscosity while reducing diffusivity, thereby suppressing the FRET signal. Furthermore, the FRET ratio remained almost unchanged even in the presence of 20% (w/v) BSA. This finding aligns with reports suggesting that the cytoplasm, though containing 200–300 mg/mL of macromolecules, exhibits a low level of crowding comparable to approximately 4% PEG1000, corresponding to about 8% excluded volume.^33,37^

Next, we applied this sensor system to the PRM–SH3 condensates to determine if the scaffold density-dependent kinetic differences arose from variations in effective concentration (Fig. 4E,F). The PRM–SH3_long_ scaffold exhibited a FRET ratio comparable to that observed under the 150 μM bulk condition. In contrast, the original PRM–SH3 scaffold showed a higher FRET ratio, closer to the 200 μM benchmark. Based our bulk observation that high viscosity suppresses FRET and the lower diffusivity of the original PRM–SH3 condensate (Fig. 3D), the observed FRET ratios might be conservative underestimates of the actual values. Considering these findings in combination, the observed increases in effective concentration for these two types of PRM–SH3 condensates largely explain the corresponding reaction rate enhancements of ×1.31 (long-linker) and ×2.19 (original).

### Rate acceleration beyond effective concentration factor in LAF condensates

To examine whether the effective concentration factor observed in PRM–SH3 condensates also applies to condensates formed by intrinsically disordered proteins (IDPs), we measured the ST–SC reaction rates within LAF condensates. Interestingly, the LAF condensates exhibited significantly faster reaction rate (×6.74) than the PRM–SH3 system (×2.19), despite the two condensates having comparable scaffold densities and FRET-based effective concentrations (Fig. 4B,E). To further probe the driving forces behind this phenomenon, we reduced ionic strength to enhance electrostatic interactions that govern LAF LLPS. Lowering the NaCl concentration from 155 mM to 40 mM substantially accelerated the reaction, with the rate constant increasing to 4210 M^-1^ s^-1^ (Fig. 4G). This represents a 57.7-fold enhancement relative to the bulk phase. While lowering the salt concentration increases effective client concentration (Fig. 4E,F), the magnitude of the observed acceleration was disproportionately large and cannot be explained by the effective concentration alone. Together, these results indicate that the LAF microenvironment provides additional physicochemical advantages for the ST–SC reaction.

### Acceleration of ST–SC reaction in a densely packed hydrophilic molecular environment

We investigated how the residue composition of the IDP scaffold influences the ST–SC reaction kinetics to uncover the specific factors within the LAF condensates that drive additional rate acceleration. To this end, we generated mixed condensates by replacing a portion of the PRM–SH3 scaffold with various IDP scaffolds. Notably, mixed condensates containing the charged-residue-rich LAF or TAF scaffolds exhibited significantly faster reaction kinetics than those observed in the original PRM–SH3 system or the hydrophobic FUS-mixed variant (Fig. 5A). This result was particularly striking because the FUS-mixed system exhibited the highest FRET ratios among all tested conditions, implying a higher effective concentration of the ST–SC components (Fig. 4E). Given that the primary distinction of FUS from LAF and TAF is its significantly lower charge density, we hypothesized that the high charged residue content generates a relatively hydrophilic microenvironment that facilitates the ST–SC reaction. Supporting this interpretation, the dramatic acceleration of reaction rates in the LAF condensate under low-salt conditions (Fig. 4G) suggests that reducing ionic shielding and strengthening electrostatic interactions promote a microenvironment more favorable for the reaction.

**Fig. 5.**
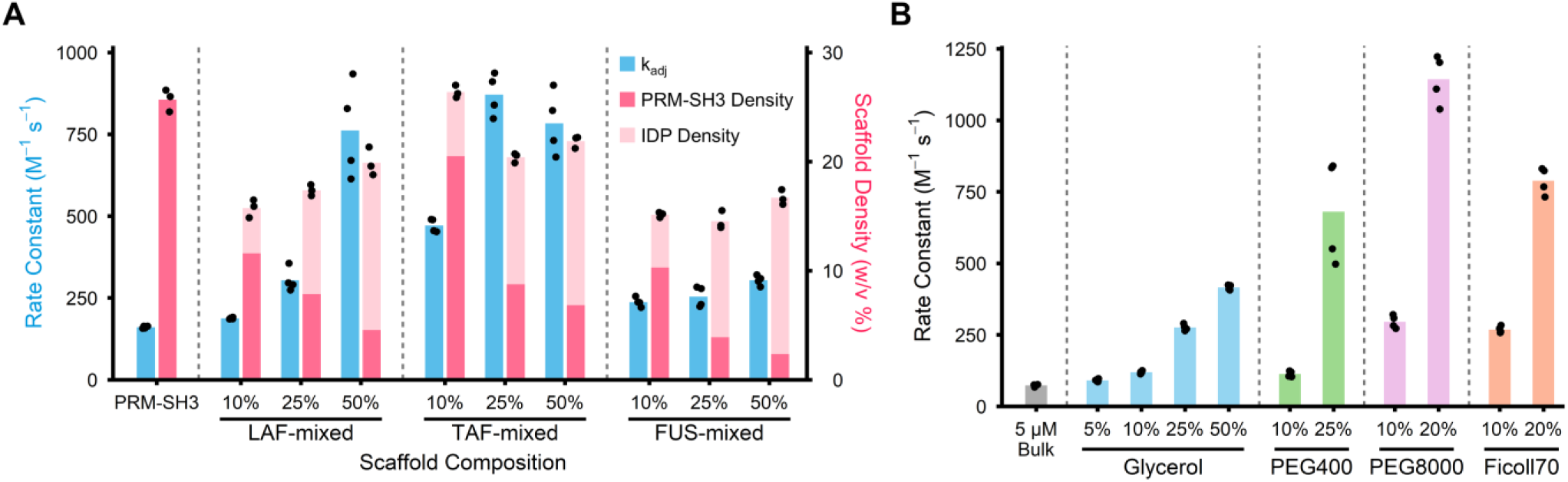
Acceleration of ST–SC reaction in a densely packed hydrophilic molecular environment. (**A**) Comparison of the reaction rate constant (blue) and scaffold w/v density (pink) within mixed condensates of PRM–SH3 and IDP (LAF, TAF, and FUS). (**B**) Rate constants for the ST–SC reaction in bulk solution containing various cosolutes.

To verify the role of hydrophilicity in accelerating the ST–SC reaction, we introduced hydrophilic additives to the bulk solution. The addition of glycerol, a small hydrophilic molecule with negligible excluded-volume effects, markedly enhanced the reaction rate (Fig. 5B). A similar trend was observed with PEG400. Although PEG400 contributes minimally to excluded-volume effects, it produced a rate increase that exceeded expectations based on its calculated excluded-volume contribution. This enhancement can be attributed to PEG400’s greater hydrophilicity relative to PEG8000. Similarly, Ficoll70, which can be considered a nearly spherical crowder, exhibits a relatively low excluded-volume effect for its molecular weight, approximately one-third of that of PEG8000 (see Supplementary Materials Section 1.10). Nevertheless, it substantially increases the reaction rate, likely due to the hydrophilic nature of the Ficoll polymer, which is enriched in hydroxyl groups.

### Internal hydrophilicity governs condensate-specific reactivity

We next characterized the internal hydrophilicity of various condensates using PRODAN, a polarity-sensitive fluorescent dye. PRODAN exhibits a characteristic red shift in its emission maximum as the surrounding environment becomes more hydrophilic.^6,38,39^ By calculating the emission intensity ratio between the long-wavelength (500–550 nm) and short-wavelength (430–475 nm) bands, we quantified the relative hydrophilicity of each condensate. As expected, LAF condensates with high client reactivity displayed significantly higher hydrophilicity ratios than the PRM–SH3 (Fig. 6A). Among all condensates examined, the LAF condensate at low salt (40 mM NaCl) exhibited the most significant red shift in PRODAN emission, revealing the highest internal hydrophilicity and suggesting that this property critically contributes to the pronounced reactivity. Similarly, TAF-mixed PRM–SH3 condensates, again with high client reactivity (Fig. 5A), exhibited overall higher hydrophilicity than original PRM–SH3 condensates, with a modest decrease at 50% TAF that mirrored the slight reduction in reaction rate observed under the same condition (Fig. 6B). In contrast, the FUS scaffold exhibited strong hydrophobicity, which led to PRODAN association with FUS and prevented accurate quantification (Fig. 6C). Nevertheless, the overall emission profile of the FUS-mixed condensate was substantially blue-shifted relative to PRM–SH3 condensates, consistent with the formation of a hydrophobic interior. Consequently, the observed rate enhancement in the FUS-mixed system appears to be driven by the increase in effective concentration, as in PRM–SH3, without the synergistic advantages provided by high hydrophilicity.

**Fig. 6.**
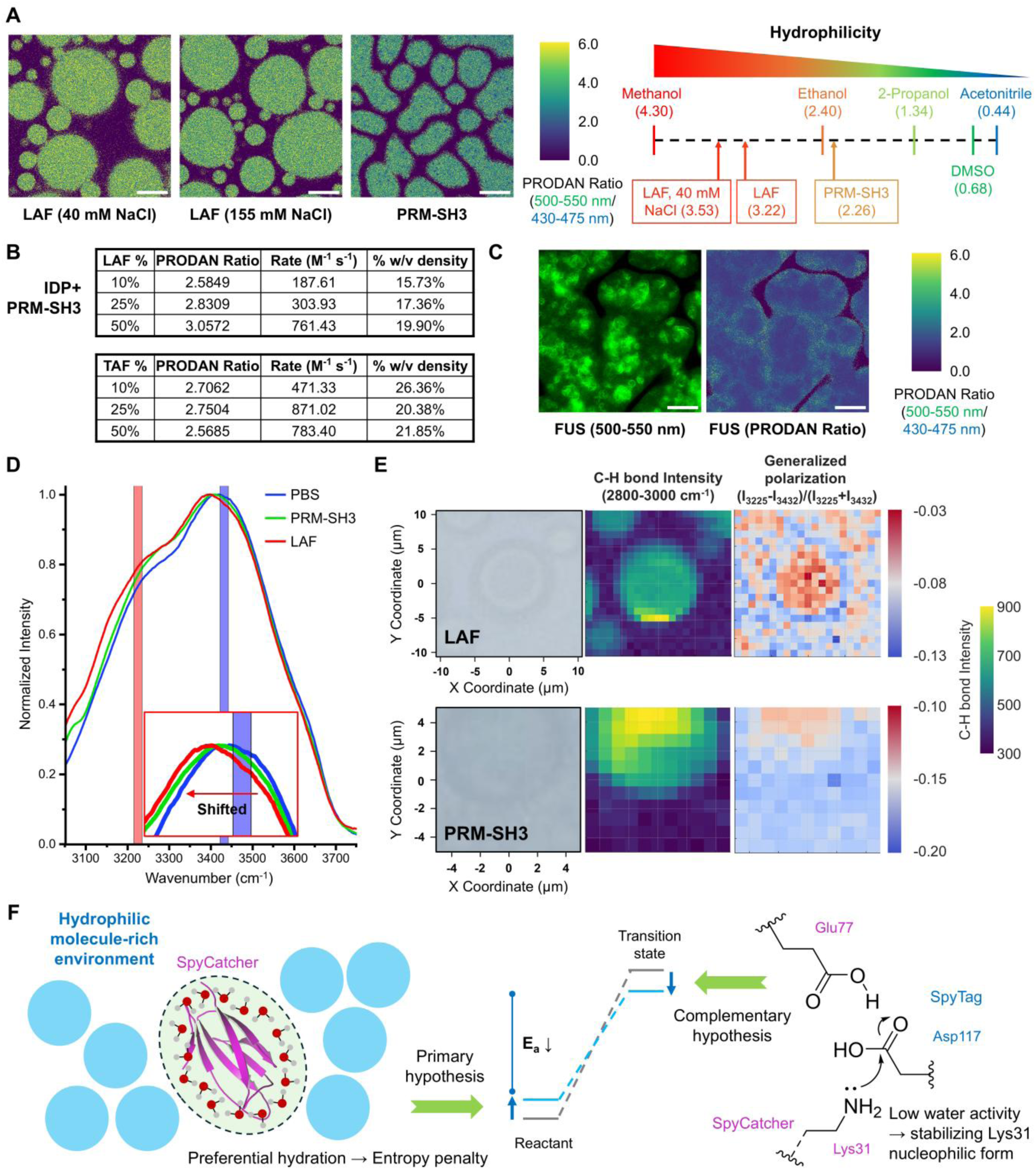
Internal hydrophilicity governing condensate-specific ST–SC reactivity. (**A**) Hydrophilicity mapping of condensates using the polarity-sensitive dye PRODAN. The ratiometric images represent local hydrophilicity, where a higher ratio (yellow) indicates a more hydrophilic environment. Scale bars, 20 µm. Emission ratios of organic solvents are provided for comparison to assess the relative hydrophilicity of LAF and PRM–SH3 condensates. (**B**) Tables summarizing the PRODAN ratio, reaction rate constant, and total scaffold density (% w/v) for PRM–SH3 condensates mixed with either LAF or TAF at varying compositions. (**C**) Representative PRODAN fluorescence and ratiometric images showing the internal environment of FUS droplets. Scale bars, 20 µm. (**D**) Normalized Raman spectra of the O–H stretching band comparing PBS buffer, PRM–SH3, and LAF condensates. A progressive blue shift in the peak position (inset) indicates an alteration in water activity. (**E**) 2D maps of LAF (top) and PRM–SH3 (bottom) condensates showing the bright-field image, C–H bond intensity, and the generalized polarization of water, derived from the intensity ratio of different O–H stretching modes. (**F**) Proposed mechanisms for accelerated reactivity via a hydrophilic microenvironment. The primary hypothesis involves preferential hydration and an entropy penalty, while the complementary hypothesis suggests that low water activity stabilizes the nucleophilic form of Lys31, both of which contribute to reducing activation energy.

We also investigated water activity within LAF and PRM–SH3 condensates by using confocal Raman microscopy. The O–H stretching band of water varies with hydrogen-bonding states, with tetrahedral and distorted (di-coordinated) water molecules giving rise to peaks near 3225 and 3432 cm⁻¹, respectively.^39^ Stronger hydrogen bonding corresponds to a reduced fraction of free water and, consequently, lower water activity. These differences were quantified using the generalized polarization function, defined as GP(tetra/di)=(I_3225_−I_3432_)/(I_3225_+I_3432_). When comparing PBS buffer, PRM–SH3 condensates, and LAF condensates, we observed a progressive increase in GP values, accompanied by a corresponding spectral shift (Fig. 6D,E), indicating that the hydrophilic LAF scaffold binds water more effectively and reduces the fraction of free water.

Recent studies have increasingly sought to explain how enzymatic activity within condensates is modulated by the charge content or hydrophobicity of scaffold biomolecules.^40,41^ Mechanistic analyses of hydrophobicity influence have primarily focused on factors such as macroscopic diffusivity assessed by FRAP, reactant recruitment quantified by partition coefficients, and restricted substrate accessibility. However, these factors fail to explain our observations, such as the acceleration of the reaction by glycerol in bulk solution despite reduced diffusivity and fundamentally unchanged reactant concentration.

Our primary hypothesis for enhanced reactivity by internal hydrophilicity is that the presence of hydrophilic molecules around proteins induces preferential hydration.^42^ This preferential hydration promotes partial pre-rigidification of the protein surface, reduces conformational entropy, and elevates the ground-state free energy of the reactants, effectively lowering the activation barrier (Fig. 6F, left). This mechanism is consistent with the design principle underlying the enhanced reactivity of the SpyTag003–SpyCatcher003 pair.^19^ Theoretical studies of how preferential hydration influences protein structure and enzymatic reactions have long been investigated in biophysics.^43^ Unlike reactions involving small organic molecules, biochemical reactions often exhibit pronounced enthalpy-entropy compensation due to the large number of associated water molecules and conformational complexity of macromolecules. As a result, enthalpic stabilization imposes substantial entropic penalties on the overall free-energy landscape. In this context, recent studies showing that the presence of Ficoll70 elevates the free-energy level of enzyme-substrate complexes provide strong support for this framework, as they indicate that preferential hydration can reduce the activation energy.^44^ A complementary hypothetical mechanism supported by Raman microscopy data is that hydrophilic crowders lower the local water activity (a_w_) and reduce the dielectric constant within the hydrophobic ST–SC binding pocket.^45^ This reduction shifts the pK_a_ of Lys31 downward, stabilizing its neutral nucleophilic form, while maintaining the optimal charge distribution of the Asp117/Glu77 proton-shuttle network (Fig. 6F, right). These changes collectively facilitate the nucleophilic attack by Lys31, thereby accelerating the covalent ligation reaction.

### Dissecting the contributions of effective concentration and hydrophilicity to reaction kinetics

Across the diverse scaffolds examined, excluding the relatively hydrophobic PRM–SH3 and FUS condensates, changes in reaction rate arose from the combined influence of effective concentration and a hydrophilic microenvironment. Because the designed FRET-based client pair and PRODAN analysis enable quantitative decoupling of these two distinct contributions, we additionally modified condensate microenvironments to further explore the underlying kinetic principles within condensates. We first investigated how the addition of a bulky domain to the scaffold architecture modulates the physical environment within the condensate. To this end, we generated LAF scaffolds recombinantly fused with a bulky, folded maltose-binding protein (MBP) designated as MBP–LAF–SUMO–His. Interestingly, when half of the LAF scaffold was substituted with this MBP-fused counterpart (half-substituted MBP/LAF condensate), the total scaffold density of the condensate decreased to less than half of the original value. However, the rate constant remained 6.82-fold higher than that of the bulk solution, comparable to that of the original LAF condensate (Fig. 7A). Even at 100% substitution, where the condensate density dropped to a mere 1.63% (w/v), representing approximately 7% of the original LAF density, the rate constant was still 2.73-fold higher than the bulk phase.

**Fig. 7.**
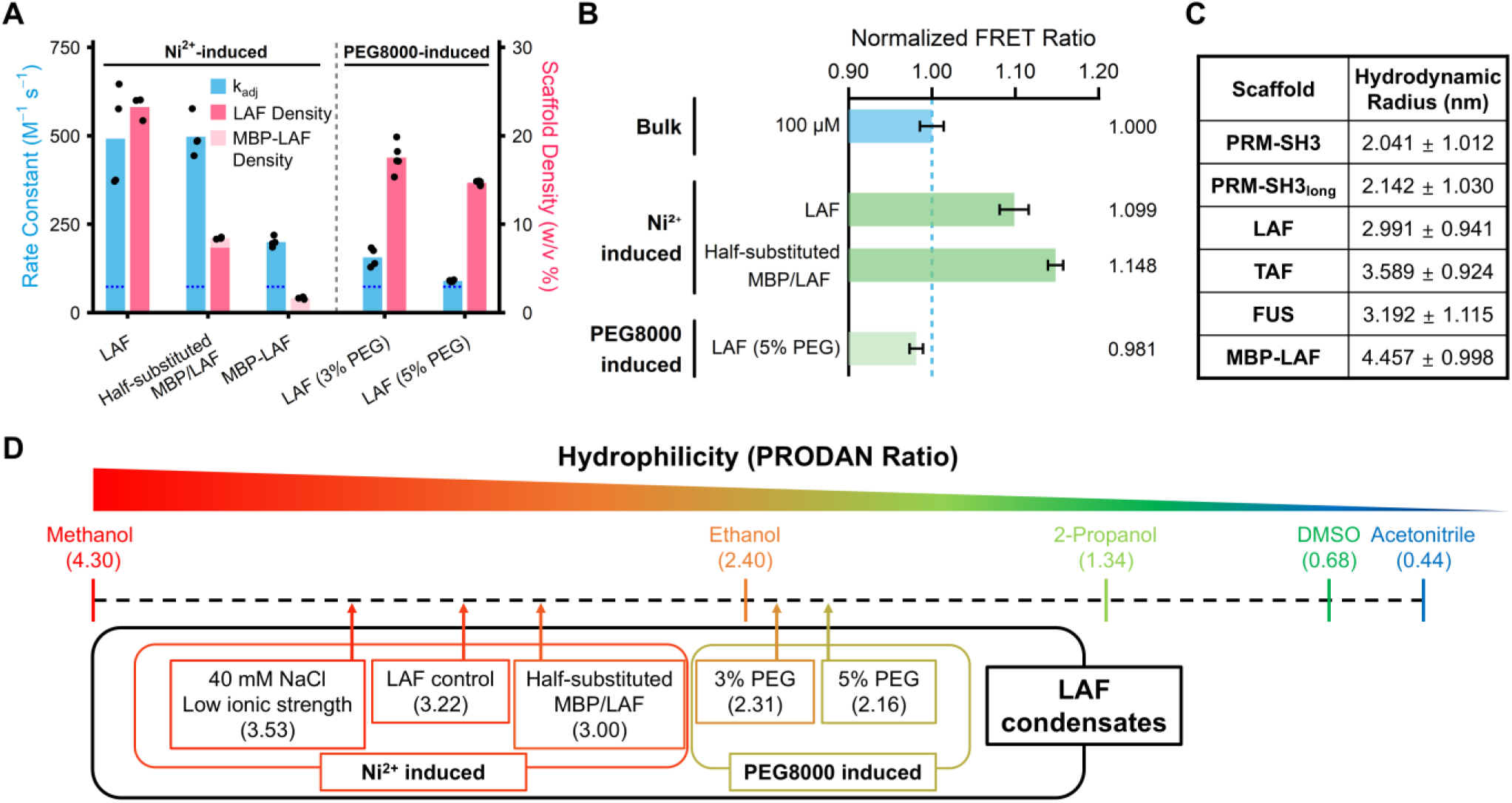
Decoupling the contributions of effective concentration and microenvironmental hydrophilicity on ST–SC reaction kinetics. (**A**) Rate constants and corresponding scaffold densities are shown for Ni^2+^-induced and PEG8000-induced LAF condensates. Blue dotted lines represent the rate constant of 5 μM bulk solution. (**B**) Normalized FRET ratios of the probes indicate the degree of probe compaction, reflecting the effective concentration effect within the bulk solution and various LAF condensates. Error bars represent ±1 SD (N=5). (**C**) Table summarizing the hydrodynamic radius values (mean size ± SD of the number-weighted distribution) measured by dynamic light scattering for various scaffold proteins in their dispersed state, providing a baseline for their physical dimensions. (**D**) Comprehensive hydrophilicity scale measured by PRODAN for various LAF condensate conditions.

This sustained acceleration despite the drastic reduction in scaffold density can be explained by the effective concentration factor. In the half-substituted MBP/LAF condensate, the FRET ratio was notably higher than that of the original LAF condensate, suggesting that the effective concentration of clients remains high or is even amplified by the presence of bulky folded domains (Fig. 7B). This interpretation is supported by dynamic light scattering (DLS) measurements, which revealed a significantly larger hydrodynamic radius for the MBP-fused scaffold (4.457 nm) compared to the original LAF scaffold (2.991 nm) (Fig. 7C). While IDPs generally possess larger hydrodynamic radii than folded proteins of equivalent molecular weight,^46–48^ our DLS and FRET data imply that the combination of an IDR with a bulky folded domain offers a high excluded volume despite the reduction in scaffold mass density (w/v %). The architectural synergy arising from the combination of the IDP’s inherently expanded conformational volume and the discrete steric footprint of the fused folded domain should therefore be carefully integrated into evaluations of the interplay between molecular crowding and internal reactivity.

However, increased effective concentration does not fully account for the observed similarity in reaction rates. PRODAN analysis of the half-substituted MBP/LAF condensate revealed a decrease in internal hydrophilicity compared to that of the original LAF system, as shown in Fig. 7D. This indicates that the comparable reaction rate is a balanced outcome. While the bulky MBP domain maximizes effective concentration through the excluded-volume effect, it simultaneously creates a slightly less hydrophilic environment, thereby inhibiting the even higher accelerations that might be expected from the FRET increase alone.

The dissection of both factors was additionally investigated by examining PEG8000-induced LAF condensates. Unlike their metal-ion-induced counterparts, these condensates exhibited lower effective concentration (Fig. 7B) and reduced internal hydrophilicity (Fig. 7D) relative to the original LAF, resulting in only modest (3% PEG) or negligible (5% PEG) increases in rate (Fig. 7A). Collectively, these findings demonstrate that effective concentration and hydrophilicity act as independent yet synergistic modulators of ST–SC reaction kinetics. By integrating solvatochromic PRODAN analysis with discretized FRET sensors, we provide a robust framework for dissecting the distinct microscopic physical and chemical drivers of reactivity within biomolecular condensates.

## Discussion

In this study, we established a quantitative framework to dissect how microenvironmental factors within biomolecular condensates govern reaction kinetics beyond the classical law of mass action. By employing the well-characterized SpyTag–SpyCatcher protein ligation, a reaction with kinetics that are not diffusion-limited, we were able to isolate the chemical step and effectively exclude translational diffusion effects. We systematically analyzed diverse condensate systems differing in effective concentration (excluded-volume fraction), internal hydrophilicity, and macromolecular architecture.

Our findings offer a new perspective on molecular enrichment within condensates, suggesting a fundamental shift in how we conceptualize internal concentration. We demonstrate that the reaction kinetics are governed not by the simple global reactant concentration partitioned into the dense phase, but by the local effective concentration precisely constrained by the scaffold’s specific architecture and excluded-volume characteristics. Additionally, these excluded-volume effects do not scale directly with condensate density but instead depend strongly on scaffold geometry and hydration characteristics. For instance, condensates formed with scaffolds containing both bulky folded and disordered domains illustrate that enlarged molecular dimensions can preserve the effective concentration of clients despite decreased overall density, underscoring the nonlinear nature of these effects.

Furthermore, internal hydrophilicity can exert a potent impact, driving rate enhancements of up to several dozen-fold. Recent studies have reported that the solvent properties of biomolecular condensates can resemble those of organic solvents.^49^ Such solvent-like microenvironments not only modulate the recruitment and partitioning of small molecules and macromolecules but also suggest that LLPS can serve as a natural mechanism for tuning intracellular reaction conditions. Although the SpyTag–SpyCatcher ligation represents a single model system, many biochemical reactions are intrinsically water-dependent, and a wide range of enzymes, particularly hydrolases and dehydratases, exhibit marked sensitivity to changes in hydrophilicity or to low-water-activity conditions.^50–52^ These precedents support the notion that the hydration state within condensates can directly modulate biochemical reactivity. Collectively, these indicate that the biomolecular condensates do not merely act as a passive host for partitioning; rather, they create a discretized, LLPS-mediated local environment, ultimately serving as a tunable driver of internal reactivity (Fig. 8).

**Fig. 8.**
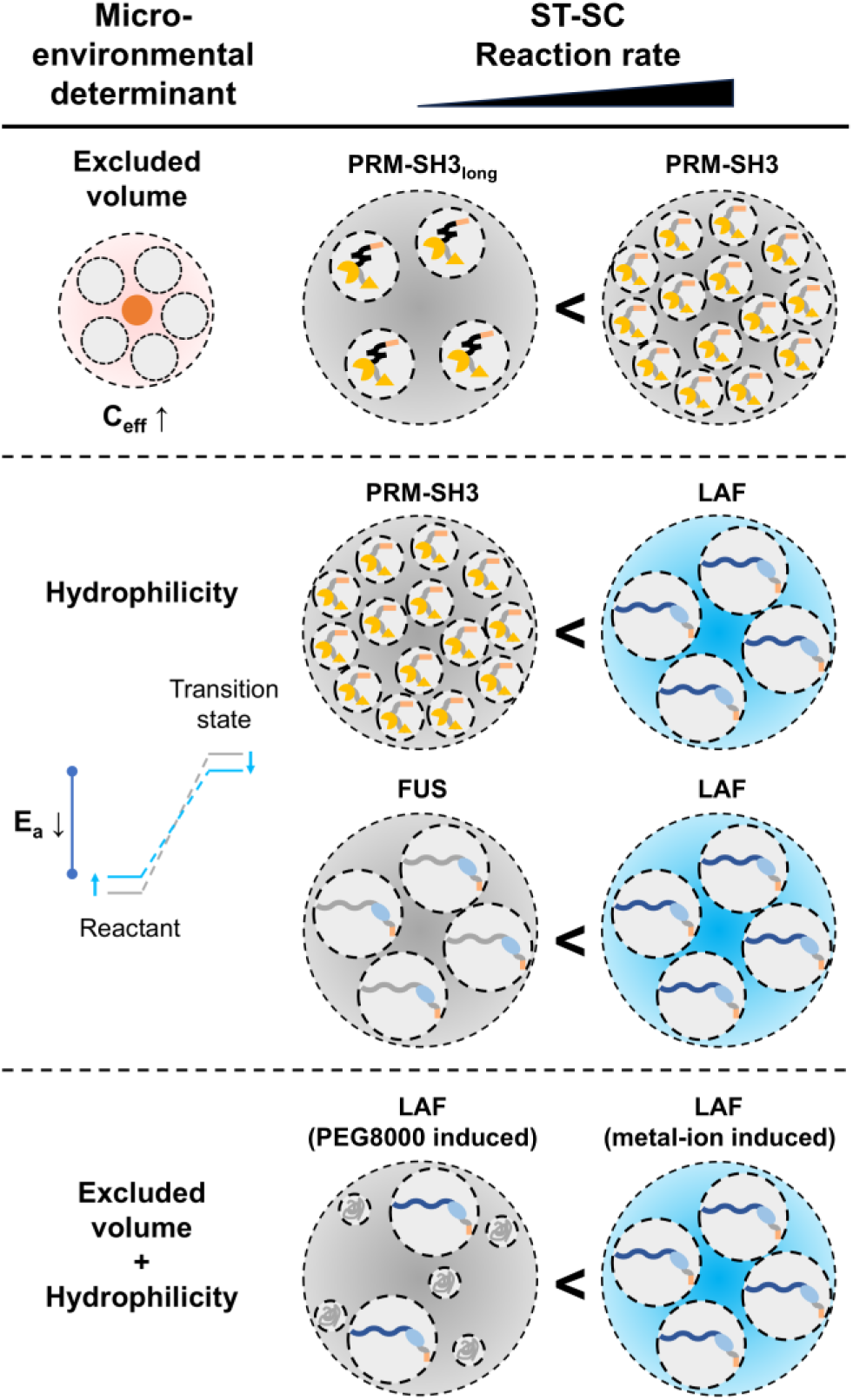
Proposed model for the microenvironmental regulation of reaction kinetics. This schematic summarizes how physical and chemical determinants within condensates synergistically govern ST–SC reactivity. The reaction rate is enhanced by excluded-volume effects, which increase effective concentration, and is further accelerated in hydrophilic microenvironments, which lower the activation energy. The interplay between these factors dictates the distinct kinetic profiles observed across different condensates.

This framework bridges microscopic physicochemical parameters with macroscopic reaction outcomes and provides generalizable design principles for engineering synthetic condensates as tunable microreactors. It gives a conceptual roadmap for the rational design of tailor-made condensates, where microenvironments can be precisely optimized for specific catalytic pathways. Future studies integrating single-molecule spectroscopy and computational modeling may further elucidate how local hydration dynamics and dielectric fluctuations couple to both ground-state energetics and transition-state stabilization, deepening our understanding of biochemical regulation within membraneless organelles.

## Materials and Methods

### Protein expression and purification

All proteins used in this study were cloned into the pET-21a(+) vector. Complete protein sequences are provided in Table S1 and Table S2. The recombinant plasmids were transformed into *Escherichia coli* BL21(DE3) cells and cultured in LB medium supplemented with 100 µg/mL ampicillin at 37 °C. When the OD_600_ reached 0.5–0.7, protein overexpression was induced by adding 1 mM isopropyl β-D-1-thiogalactopyranoside (IPTG), followed by overnight incubation at 20 °C. Cells were harvested by centrifugation at 6,000 rpm (∼6,400 × g) and washed once with 1× PBS (160 mM NaCl, pH 7.3). The cell pellets were resuspended in Ni-IDA binding buffer (500 mM NaCl, 50 mM Tris-HCl, pH 8.0) and lysed by sonication. Lysates were clarified by centrifugation at 12,000 rpm (∼16,000 × g) for 15 min, and the supernatants were collected. The supernatant was loaded onto a Ni-IDA Excellose (Takara) resin column, washed with wash buffer (50 mM Tris-HCl, 500 mM NaCl, 50 mM imidazole, pH 8.0), and eluted with elution buffer (50 mM Tris-HCl, 500 mM NaCl, 500 mM imidazole, pH 8.0). For IDP scaffolds, the binding, washing, and elution buffers were modified by the addition of 10% glycerol and optimized NaCl concentration (500 mM for LAF, 300 mM for TAF, 200 mM for FUS, 0.3 mM EDTA can be added to binding buffer to prevent nonspecific binding) to enhance protein stability. Protein solutions were dialyzed against 1× PBS at 4 °C. For IDP scaffolds, dialysis was performed in 1× PBS containing 10% glycerol at 28 °C. Final protein solutions were aliquoted and flash-frozen in liquid nitrogen and stored at −80 °C for long-term use. Prior to use, IDP scaffolds were incubated at 37 °C, and other proteins at room temperature for at least 30 min. To eliminate potential effects of Ni²⁺–His₆ interactions from the client proteins, His-tags were removed by TEV protease cleavage. The client protein solutions were incubated with TEV protease at a protease-to-protein molar ratio of 1:25 at 4 °C for 24 hours. The resulting mixtures were further purified by Ni-IDA chromatography to remove uncleaved protein and residual TEV protease.

### Measurement of reaction progress via electrophoretic mobility shift assay (EMSA)

#### Bulk solutions

The PBS-based solutions containing designated additives (e.g., crowding agent, glycerol) were prepared. SpyTag–SpyCatcher or SpyTag003–SpyCatcher003 client pairs were then added to the solution at final concentrations. The reaction mixture was aliquoted into separate tubes corresponding to the number of time points and incubated at 25 °C. At each time point, the reaction was quenched by adding SDS–PAGE loading dye. Samples were subsequently analyzed by SDS–PAGE on 10% acrylamide gels (120 V, 80 min). The gel bands corresponding to the reacted clients (upper band) and unreacted clients (lower band) were visualized with ChemiDoc (Bio-Rad). Their intensities were quantified by extracting the line profiles of both bands using ImageJ (Fig. S10). The conversion (reaction progress) at each time point was expressed as the ratio of the intensity of the reacted client band to the total client band intensity. This final conversion value represents the average of the individual reaction extents calculated for both the mCherry–SpyTag and GFP–SpyCatcher clients.

#### Condensates

The reaction mixture was prepared sequentially by adding PBS, scaffold protein, client protein, PUMA-fused scaffold, and either NiCl₂ (equimolar concentration to scaffold unless otherwise specified) or PEG8000 in that order. When IDP scaffolds were mixed with PRM–SH3, the PUMA–PRM–SH3 was used. The reaction mixture was aliquoted into separate tubes, briefly centrifuged at 12,000 rpm (∼13,500 × g) for 1 min at 25 °C, and the supernatant was completely removed. The resulting condensate samples were incubated at 25 °C for the designated reaction period (typically 5, 10, 20, 60, 120, and 240 min). To terminate the reaction, 1× SDS–PAGE loading dye was added to the centrifuged condensate, followed by brief vortexing and 90 sec of sonication to ensure thorough dissolution before electrophoretic analysis. EMSA was performed using the same method as for the bulk solution. The initial molar ratio of the client proteins was optimized to ensure equal internal concentrations within the condensates. Through iterative trials in which EMSA was used, ensuring that the percentage of reaction progress for each client at the same time point was almost identical. Final reaction kinetics experiments were performed using these optimized settings.

### Fitting method

The rate equation described in the results section, *v*=k_adj_([client]_t_ – F_u_[client]_0_)^2^, was employed for kinetic analysis. The parameters k_adj_ and F_u_ were treated as fitting variables. Using the integrated second-order rate equation, theoretical reaction conversions were calculated for each time point. The experimentally measured reaction progress was then compared with theoretical values, and the optimal k_adj_ and F_u_ were determined by the least-squares fitting method. Each experiment was performed in duplicate, and SDS–PAGE analysis was also conducted twice for the same quenched samples. Consequently, four gel fluorescence images were obtained in total, and the rate constant was determined by averaging the values obtained for each client across all images. In most cases, each sample was prepared and analyzed independently by different researchers, confirming a high degree of experimental reproducibility. All source data—including reaction progress at each time point, initial input ratios of mCh and GFP client proteins, internal client concentrations within the condensates, calculated rate constants, and unreacted fractions—are provided in the “Data collection of reaction progress” section of Supplementary Materials.

### Measurement of client concentration and scaffold density in condensates

To determine the client concentration (i.e., partitioning) within condensates, the mCh–ST–Bcl client was recruited using the same initial total concentrations as in the reaction progress assays. To measure scaffold density, condensates were formed by mixing the unlabeled scaffold with a small fraction (< 5%) of Cy5-labeled scaffold. 50 μL of protein solutions were loaded onto 96-well black/clear bottom plates, which were previously passivated with 2 mg/mL bovine serum albumin (BSA; Fraction V, Thermo Fisher, dissolved in PBS) for 2 h at 25 °C. The samples were then centrifuged at 4000 rpm for 10 min (to sediment the condensates) and imaged by confocal laser microscopy (A1R HD25 with Eclipse Ti2, Nikon) using a 60×/1.40 numerical aperture (NA) apochromatic oil-immersion objective lens. We used the 561 nm laser for mCherry and the 640 nm laser for Cy5-labeled proteins. Subsequently, the fluorescence intensity was fitted to a standard curve of known concentrations to calculate the final concentration, averaging the values obtained from three images.

### Fluorescence recovery after photobleaching (FRAP)

FRAP experiments were performed on the A1R HD25 confocal microscope using the 60×/1.40 NA oil-immersion objective lens. Condensates containing either Cy5-labeled scaffold (mixed at < 5%) or the mCh–ST–Bcl client were prepared in BSA-passivated 96-well black/clear bottom plates as described previously. A circular region of interest (ROI) with a 5 μm diameter, positioned near the center of the condensate, was photobleached using a single, high-intensity pulse of the corresponding laser. Fluorescence recovery within the ROI was immediately monitored by time-lapse imaging at low laser power (captured every 10 s). For data analysis, the average fluorescence intensity of the bleached ROI was measured over time. The recovery curves were normalized using the intensities measured immediately after bleaching (t=0): the average intensity of the bleached ROI was set as the 0% reference, and the average intensity of the entire bleached condensate was set as the 100% reference (N=3 condensates measured). Both the scaffold and client components exhibited similar recovery dynamics, as exemplified by the LAF condensate in Fig 3d.

### Fluorescence resonance energy transfer (FRET)

Samples were prepared in BSA-passivated 96-well black/clear-bottom plates as previously described. To compare effective concentrations, the FRET pair mCerulean–Bcl (donor) and mCitrine–Bcl (acceptor) was added to the bulk solution or recruited into condensate samples at a final equimolar concentration of 100 μM each. The recruited client concentration was adjusted through multiple iterations, which was confirmed by A1R HD25 confocal laser microscopy. FRET imaging was performed on an LSM 880 laser scanning confocal microscope (ZEISS) using a 63×/1.40 NA plan-apochromatic oil-immersion objective lens. The mCerulean donor was excited using the 405 nm laser, and two emission channels were collected simultaneously: the donor channel (450–505 nm) and the acceptor FRET channel (505–797 nm). For analysis, a ratiometric image was generated by dividing the acceptor-channel (FRET) intensity image by the donor-channel intensity image pixel by pixel. The FRET ratio (I_acceptor_ / I_donor_) was calculated by averaging the intensity values within condensates or in the bulk solution. To obtain the “Normalized FRET Ratio” (as shown in Fig. 4e), all measured FRET ratios were normalized to the FRET ratio obtained from the 100 μM bulk solution reference condition, which was set to a value of 1.00. A higher normalized ratio indicates a higher local effective client concentration. Five measurements were averaged under identical image acquisition settings.

### PRODAN hydrophilicity assay with fluorescence imaging

A 10 mM PRODAN stock solution was prepared in DMSO due to its limited aqueous solubility. Immediately before use, 10–20 µL of the stock solution was added to 1 mL of distilled water and vortexed to achieve rapid dilution, followed by filtration through a 0.22 µm syringe filter. PRODAN was introduced at a final concentration of 10 µM prior to LLPS induction. Although higher PRODAN concentrations cause a slight red shift in the emission spectrum, even a 100-fold increase (from 5 µM to 500 µM in ethanol) yielded only a ∼20% intensity enhancement. Samples were loaded into BSA-passivated 96-well plates, and condensates gradually settled under gravity. Imaging was performed using an A1R HD25 confocal microscope under identical acquisition settings (laser power, digital gain, and pinhole size) as used for concentration measurements. Excitation was conducted at 405 nm, and emission was collected through two channels: 430–475 nm and 500–550 nm. For analysis, a ratiometric image was generated by dividing the long-wavelength channel (500–550 nm) intensity by the short-wavelength channel (430–475 nm) intensity on a pixel-by-pixel basis. The PRODAN ratio was determined by averaging intensity values inside the condensates or organic solvent phases. Each measurement was performed five times, and the results were averaged. Notably, fluorescence was detectable within condensates and organic solvents but was quenched in the aqueous phase, making measurement in the aqueous phase difficult.

### Confocal Raman microscopy

Confocal Raman microscopy was performed using an inVia Qontor (Renishaw) system equipped with a 532 nm excitation laser and a 50x objective lens. Samples (PBS buffer, PRM–SH3 condensates, and LAF condensates) were prepared on a non-coated cover glass, which was then covered with a BSA-coated slide glass. Area maps (Fig. 6e) were acquired with a spatial resolution of 1 μm x 1 μm per pixel. 2300–4300 cm⁻¹ region was analyzed and a linear baseline correction was applied to each spectrum, defined by the trend between the 2500–2700 cm⁻¹ and 3800–4000 cm⁻¹ regions. The protein-rich condensate phase was identified by integrating the signal from the C-H stretching band (2800–3000 cm⁻¹), which serves as a proxy for protein concentration. The O-H stretching band (3000–3800 cm⁻¹) was analyzed to determine the hydrogen-bonding state of water. The Generalized Polarization (GP) value was calculated on a pixel-by-pixel basis using the formula: GP(tetra/di)=(I_3225_-I_3432_)/(I_3225_+I_3432_), where I_3225_ and I_3432_ represent the spectral intensities at the average band intensities over a ±10 cm⁻¹ range centered at 3225 cm⁻¹ and 3432 cm⁻¹, respectively, corresponding to tetrahedral (strong hydrogen bonding) and distorted (weak hydrogen bonding) water structures. The representative spectra shown in Fig. 6d are baseline-corrected averages. Condensate spectra represent the average of five measurements taken at the centers of different condensates (N=5), and the PBS spectrum is the average of measurements (N=5). These average spectra were then smoothed using a moving average filter (15 data points, corresponding to approx. 30 cm⁻¹).

### Dynamic light scattering (DLS)

Protein scaffold solutions were filtered and diluted to 20 μM. 100 μL of the solutions in disposable solvent-resistant micro cuvettes (ZEN0040, Malvern Instruments) were analyzed (173° backscattered) with a Zetasizer Nano ZS DLS system (Malvern Instruments) at 25 ℃. Each measurement averaged ten runs with a 10 s running time per run. Data processing was performed with the Zetasizer software.

## Acknowledgements

This work was supported by the National Research Foundation of Korea (NRF) grants (2023R1A2C2005183) and the Global Science Research Center Program (RS-2024-00411134) funded by the Korean government (MSIT).

## Competing interests

The authors declare that they have no competing interests.

## Author contributions

J.B.: Conceptualization, methodology, validation, formal analysis, investigation, writing–original draft, writing–review and editing, and visualization. K.H.: Conceptualization, methodology, validation, formal analysis, writing–review and editing, and visualization. D.L.: Validation, formal analysis, investigation, writing–review and editing, and visualization. J.J.: Conceptualization. Y.J.: Conceptualization, resources, writing–review and editing, supervision, project administration, and funding acquisition.

## Notes

### Competing Interest Statement

The authors have declared no competing interest.

