## Supplementary Materials for "Microenvironmental Determinants of Reaction Kinetics in Biomolecular Condensates Probed with Protein Ligation"

### 1. Supplementary Figures and Supplementary Notes

#### 1.1. Sequence analysis and physicochemical properties of IDP constructs

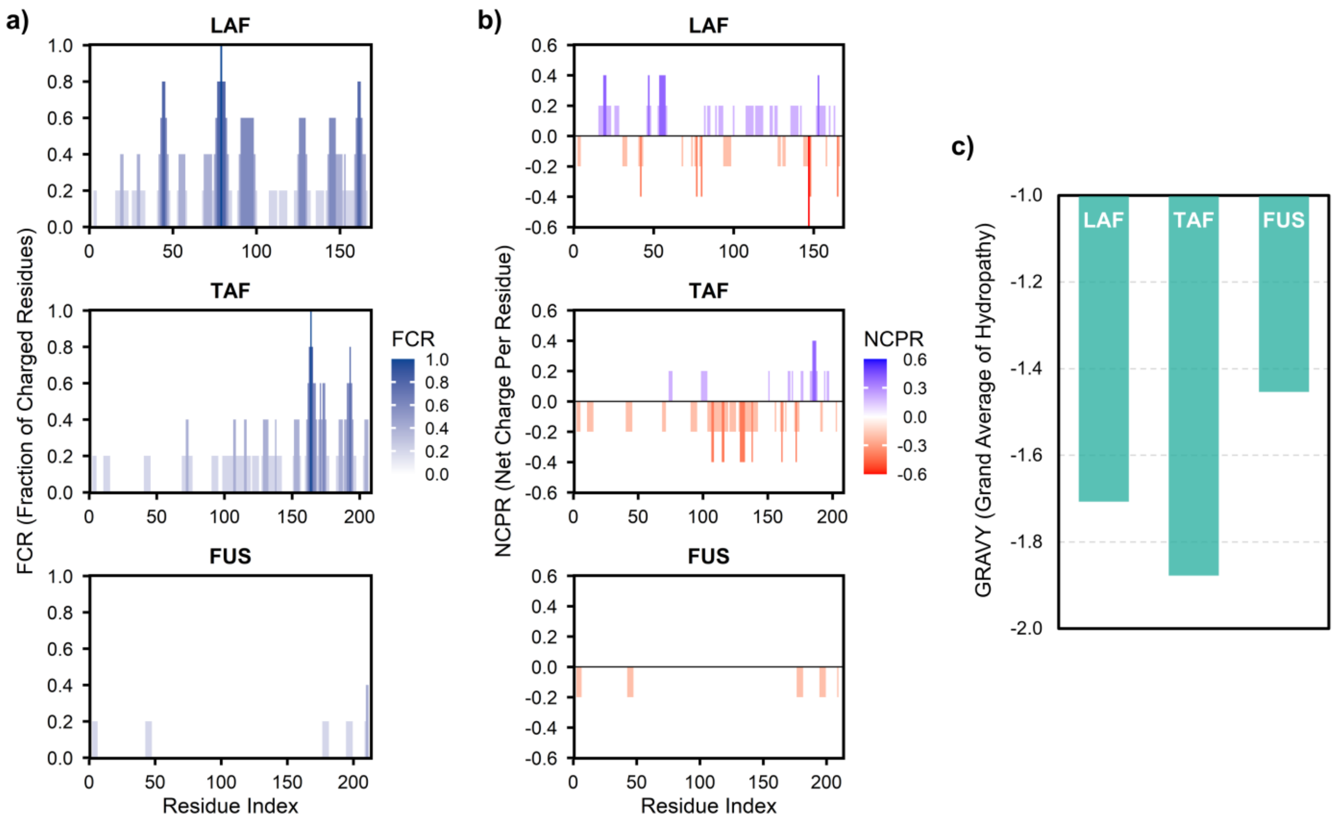

**Fig. S1.** Sequence-based physicochemical landscapes of LAF, TAF, and FUS. **a)** Fraction of charged residues (FCR) plotted against the residue index for each protein, highlighting the high charge density of the LAF RGG domain. **b)** Net charge per residue (NCPR) for each construct, where blue bars indicate positive charges and red bars indicate negative charges. **c)** Comparison of the grand average of hydropathy (GRAVY) scores for the three constructs. All scores are negative, reflecting the intrinsically disordered and hydrophilic nature of these proteins.

The sequence-based physicochemical properties of intrinsically disordered protein (IDP) scaffold constructs used in this study are shown in **Fig. S1**. To systematically investigate how different microenvironmental determinants influence protein ligation rates, we chose LAF, TAF, and FUS as representative model systems. These proteins provide distinct chemical landscapes that allow us to dissect the contributions of electrostatic and hydrophilic interactions to the observed reaction kinetics.

LAF (residues 1–168) was selected to represent a highly charged microenvironment. As shown in **a** and **b**, the LAF RGG domain possesses a high fraction of charged residues (FCR) and a distinct net charge per residue (NCPR) profile.

TAF (residues 1–208) and FUS (residues 2–214) were chosen to represent prion-like domains (SYQG-rich sequences). Although both are SYQG-rich, TAF exhibits the lowest GRAVY score among all constructs, indicating it is the most hydrophilic system, as shown in **c**. Furthermore, TAF contains a significantly higher proportion of ionizable residues than nearly neutral FUS. This selection allows us to evaluate how a highly hydrophilic, charged microenvironment within a prion-like domain modulates reaction kinetics, providing a clear contrast to the more neutral, relatively less hydrophilic environment of FUS.

#### 1.2. Condensate centrifugation for internal kinetic and concentration measurements

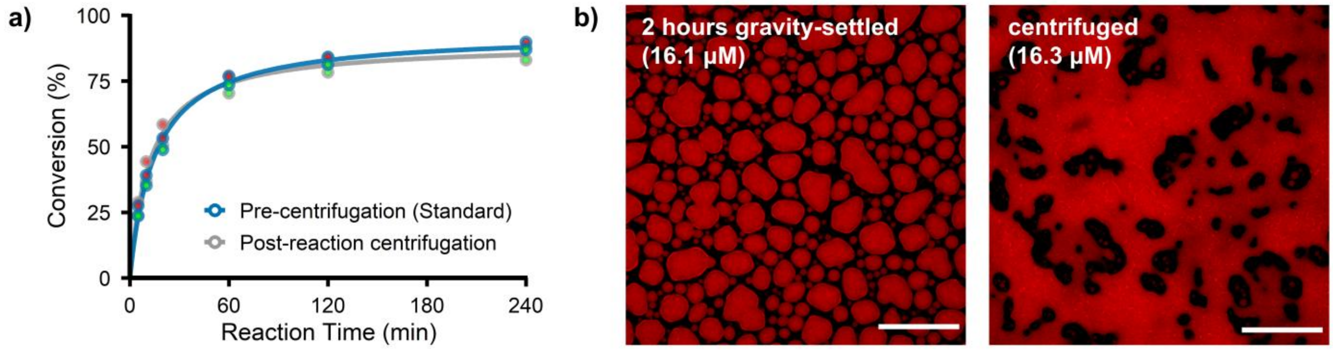

**Fig. S2.** Comparative characterization of centrifuged condensates. **a)** Comparison of reaction conversion profiles between the standard pre-centrifugation protocol (reaction proceeds inside centrifuged condensates with supernatant discarded) and post-reaction centrifugation (reaction proceeds inside condensate with supernatant) for PRM–SH3 condensates. The slight divergence observed at 240 min highlights the impact of continuous client partitioning from the dilute phase during the 4-h reaction period. mCh-ST conversion is represented by red dots, and GFP-SC conversion by green dots. **b)** Confocal microscopy images of PRM–SH3 condensates recruited with 0.2 μM mCh–ST clients. Images compare gravity-settled (2-h, left) versus immediately centrifuged (right) condensates. The parenthetical values represent the internal mCh client concentrations, which remain stable across both methods when using controlled sample volumes. Scale bars, 50 μm.

The centrifugation protocol was used to isolate the condensed phase for both kinetic and internal concentration measurements (**Fig. S2**). As shown in **a**, the timing of centrifugation does not cause a critical shift in the overall reaction rate constants. However, we observed that in the post-reaction centrifugation group, the reaction conversion decreased slightly over time compared with the pre-centrifugation protocol. This minor deviation likely results from the continuous entry of free clients in the dilute phase into the condensates. Because our protein ligation assay requires a relatively long 4-h incubation, the cumulative effect of such partitioning can vary across different condensate systems and interfere with precise kinetic analysis. To minimize these inconsistencies, we consistently applied centrifugation immediately after phase separation to remove the dilute phase and isolate the reaction environment.

Therefore, the internal scaffold densities and client concentrations were also measured within centrifuged condensates immediately after phase separation. Analyses across multiple systems revealed that the internal concentrations measured after centrifugation were consistent with those obtained using gravity-settling methods, as representatively shown in **b**. However, when the centrifuged sample volume was large, we observed an artificial increase in the internal protein concentration, likely due to centrifugal condensate stacking. To prevent this physical condensate compaction, an optimal sample volume of 50 μL in a BSA-coated 96-well plate was used throughout the analysis. These results validate the use of immediate centrifugation to characterize reaction kinetics and molecular concentrations within biomolecular condensates.

##### 1.3. Phase separation for protein condensate formation

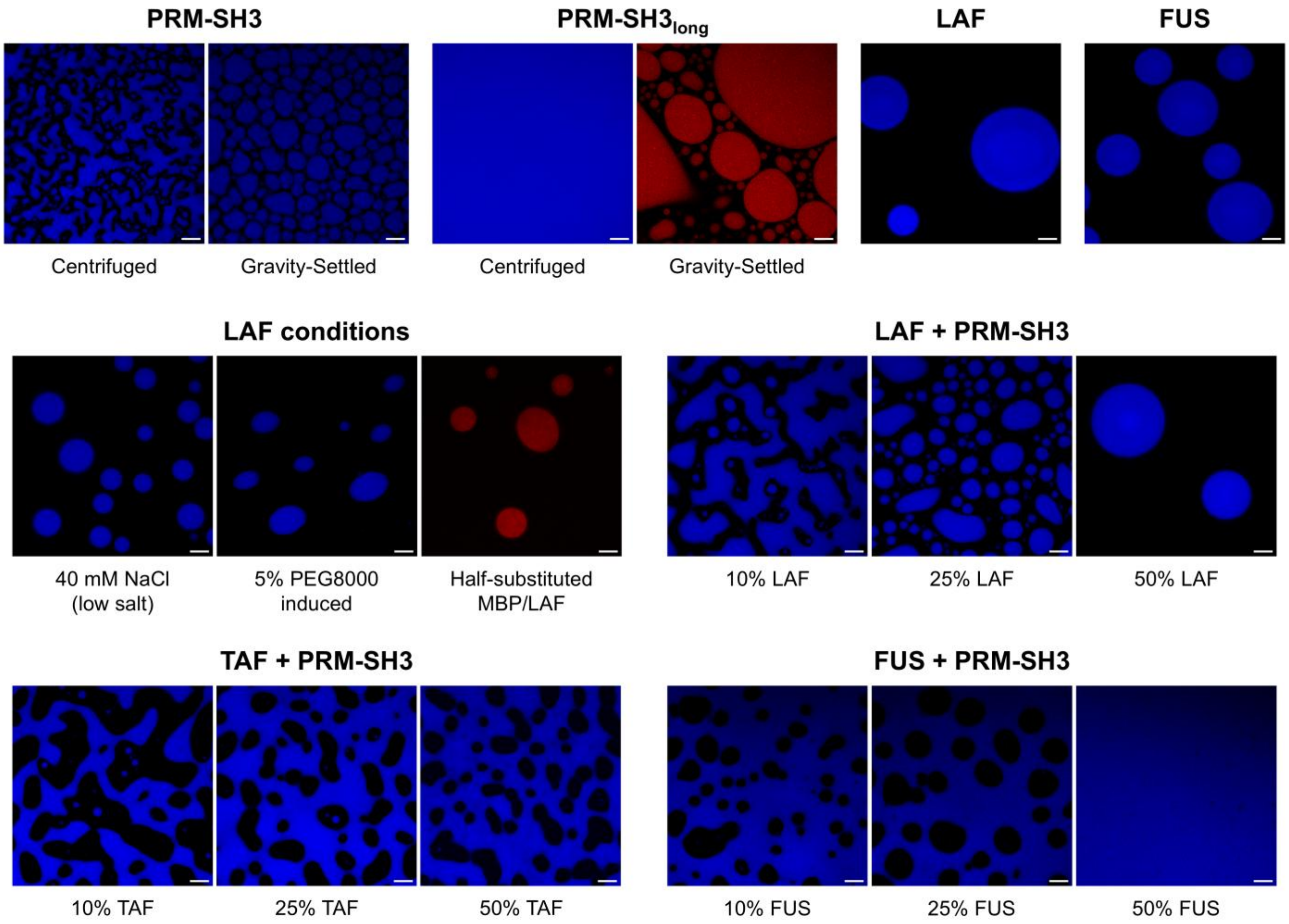

**Fig. S3.** Representative confocal laser scanning microscopy images of the diverse condensate systems characterized in this study. Condensates are visualized by Cy5-labeled scaffolds (blue) or by the client protein mCh–ST (red), as indicated. For all IDP + PRM–SH3 mixed condensates, Cy5-labeled PRM–SH3 is shown as the representative scaffold signal. In the PRM–SH3 systems, both centrifuged and gravity-settled condensates are shown, whereas in all other systems, only centrifuged condensates are displayed. Scale bars, 20 µm.

The morphological characteristics and phase-separation behavior of the various biomolecular condensate systems investigated in this work are shown in **Fig. S3**. The TAF scaffold does not undergo efficient $Ni^{2+}$-induced phase separation solely, and therefore, TAF was employed only in a mixed format with the PRM–SH3 scaffold. The multicomponent IDP + PRM–SH3 systems allowed systematic modulation of physicochemical properties within the condensate microenvironment. By adjusting the stoichiometric ratios of IDP and PRM–SH3, we implemented a gradual variation of environmental determinants, facilitating a more credible investigation of how specific internal factors influence reaction kinetics.

Unless otherwise specified, condensates were formed by adding scaffold and $Ni^{2+}$ in equimolar amounts. The total scaffold concentration was maintained at 100 µM for PRM–SH3 and IDP + PRM–SH3 mixed condensates, and 50 µM for LAF and FUS condensates. To enable client recruitment via PUMA–Bcl interaction across these systems, a PUMA-fused scaffold was incorporated at 1–10 mol%, with 5 mol% as the standard condition. For the IDP + PRM–SH3 mixed condensates, PUMA–PRM–SH3 (5 µM) was specifically utilized rather than a PUMA-fused IDP.

#### 1.4. Kinetic characterization of protein ligation pairs in bulk solution

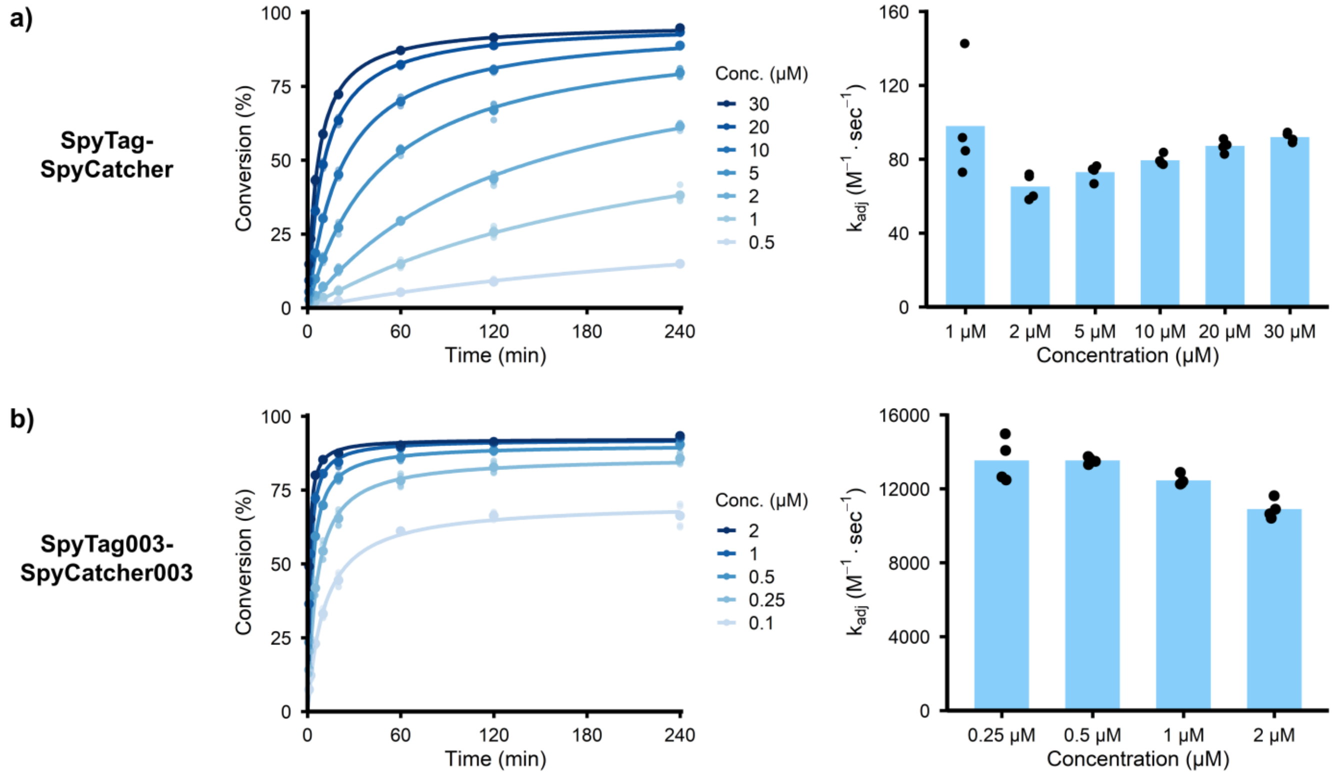

**Fig. S4.** Bulk ligation kinetics of the SpyTag–SpyCatcher and SpyTag003–SpyCatcher003 pairs. **a)** Concentration-dependent reaction conversion (left) and the corresponding $k_{adj}$ values (right) for the original SpyTag–SpyCatcher pair. This pair operates in a reaction-limited regime with relatively low rate constants. **b)** Kinetic profiles and $k_{adj}$ values for the high-affinity SpyTag003–SpyCatcher003 pair, which exhibits diffusion-influenced behavior. Solid lines in the conversion curves represent fits to the modified second-order equation, $v=k_{adj}([\text{client}]_t - F_u[\text{client}]_0)^2$.

The ligation reaction kinetics of the original SpyTag–SpyCatcher and SpyTag003–SpyCatcher003 client pairs (Bcl-fused) in bulk solutions are presented in **Fig. S4**. As these interactions follow second-order kinetics, the reaction conversion increases with increasing client concentrations. To ensure consistency with measurements within condensates where an unreacted fraction is often observed, bulk kinetic data were fitted using a modified second-order equation, $v=k_{adj}([\text{client}]_t - F_u[\text{client}]_0)^2$, where $F_u$ represents the unreacted fraction. The resulting adjusted rate constants ($k_{adj}$) are shown in the right panels of **a** and **b**.

For the original SpyTag–SpyCatcher pair shown in **a**, the rate constants remain below 100 $M^{-1} s^{-1}$, indicating a reaction-limited regime. At concentrations below 1 µM, the final conversion decreases significantly, resulting in large Fu values and reduced precision of the fitted constants. Because reactions within biomolecular condensates are generally faster than those in bulk solutions, using excessively high client concentrations can cause the reaction to terminate too quickly to capture accurate temporal profiles. Therefore, we established an optimal concentration range of 1–10 µM and adjusted the internal client concentrations of all condensates to fall within this window. The $k_{adj}$ value of 72.98 $M^{-1} s^{-1}$ obtained from the 5 µM bulk solution, which serves as the midpoint of this concentration range, was used as the reference baseline to calculate the acceleration folds reported in this study.

In the case of the SpyTag003–SpyCatcher003 pair shown in **b**, the kinetics are diffusion-influenced with rate constants exceeding $10^4$ $M^{-1} s^{-1}$. Although this pair is significantly faster, it has not yet reached a fully diffusion-limited regime. Accordingly, reactions at 0.1 µM do not reach completion, similar to the original pair. Furthermore, at concentrations above 2 µM, the reaction proceeds nearly to completion within the first 20 min. Based on these observations, a concentration range of 0.25–1 µM was selected as optimal for investigating diffusion-influenced kinetics of 003 clients in the subsequent Supplementary Note section.

#### 1.5. Differential impact of diffusivity on 003-client ligation kinetics

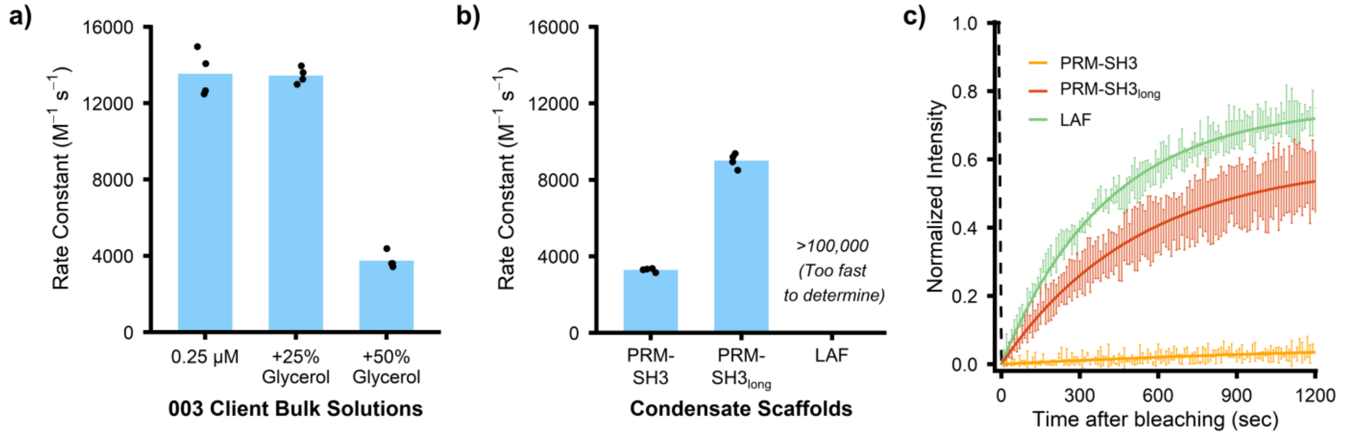

**Fig. S5.** Impact of microenvironmental diffusivity on diffusion-influenced ligation kinetics. **a)** Rate constants of the 003 client pair in bulk solutions with varying glycerol concentrations. **b)** Comparison of 003 client rate constants across different condensate scaffolds. **c)** FRAP recovery curves for different condensate systems. Error bars represent ±1 standard deviation (N=3).

To further investigate how microenvironmental properties specifically modulate diffusion-influenced reactions, we compared the kinetic responses of the SpyTag003–SpyCatcher003 (003 client) pair to the original reaction-limited pair under varying viscosity and diffusivity conditions (**Fig. S5**). As noted in the main text (Fig. 5b), the addition of glycerol, a hydrophilic small molecule, increases the reaction rate of the original ST–SC pair by creating a favorable microenvironment. However, when the same experiment is conducted with the 003 client pair in bulk solution, the results differ significantly, as shown in **a**. While the rate remains nearly unchanged in 25% glycerol, it decreases by more than half in 50% glycerol. This behavior suggests a competitive interplay between the rate-enhancing effect of the hydrophilic microenvironment and the rate-inhibiting effect of increased viscosity, which reduces molecular diffusivity. These findings confirm that, unlike the non-diffusion-limited original pair, the 003 client pair is heavily influenced by the physical diffusive constraints.

Similar trends were observed within the condensates, as illustrated in **b**. Although the original ST–SC pair exhibits slightly increased rate constants in PRM–SH3 condensates due to elevated effective concentrations (Fig. 4b), the 003 client pair shows inhibited kinetics in these diffusion-limited environments. Notably, in PRM–SH3 condensates with highly restricted internal mobility (**c**), the 003 client reaction rate decreases dramatically, whereas it decreased only slightly in the more fluidic PRM–SH3$_{long}$ condensates. In contrast, LAF condensates provide higher diffusivity and a more hydrophilic microenvironment. The reaction in LAF condensates was so rapid that it approached completion almost immediately, making it difficult to accurately determine a discrete rate constant using our method.

#### 1.6. Reversibility of reaction arrest induced by crowding agent

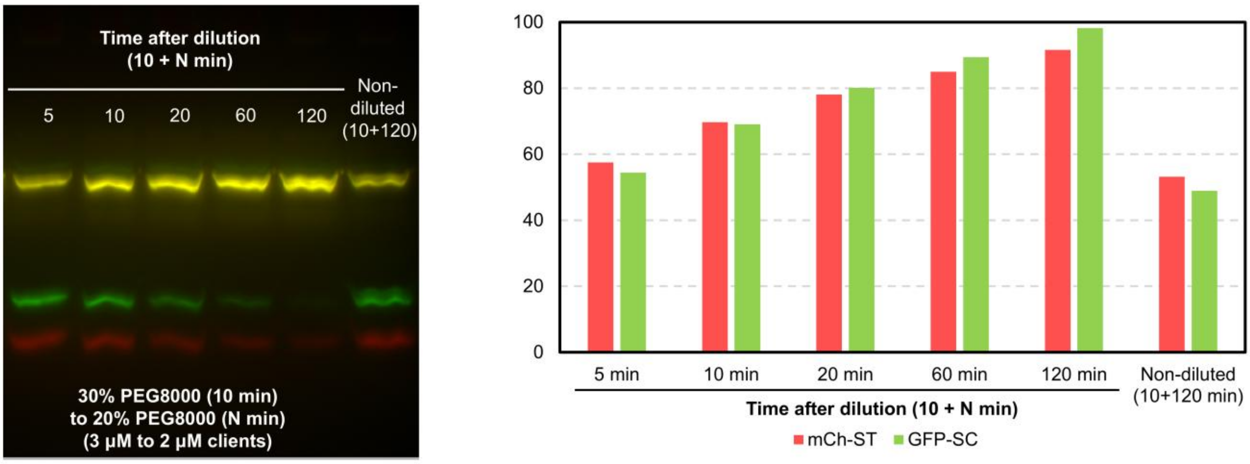

**Fig. S6.** Reversibility of ligation reactions upon dilution of the crowding agent. The left panel shows the PAGE band shift assay used to monitor reaction progress after diluting the reaction solution from 30% to 20% PEG8000. The right bar graph shows the quantitative conversion of mCh–ST (red) and GFP–SC (green) over time. While the non-diluted control (10+120 min) remains at approximately 50% conversion, the diluted samples show a progressive increase in ligation, eventually reaching near completion.

As discussed in the main text (Fig. 3a), high molecular crowding in 30% PEG8000 induces client compartmentalization and an incomplete reaction conversion of approximately 50%. **Fig. S6** demonstrates that this kinetic arrest is reversible and can be overcome by diluting the crowding agent. In the absence of dilution, the reaction conversion remains stagnant at around 50% even after 130 minutes of total incubation. However, when the crowding concentration is reduced to 20% PEG8000 after an initial 10-minute period, the reaction resumes and reaches near completion over the subsequent 120 minutes. Following the dilution step, the total client concentration is adjusted to 2 μM. Since roughly 1 μM of the client remains unreacted at the 10-minute mark, the observed recovery kinetics closely resemble the behavior of a standard 1 μM client solution in 20% PEG8000 (data for 1 μM client in 20% PEG8000 can be found in Section 5). These results confirm that reducing molecular crowding effectively reverses the compartmentalization of reacting species, allowing the protein ligation reaction to proceed toward completion.

#### 1.7. Kinetic stability across varying salt concentration in bulk solution

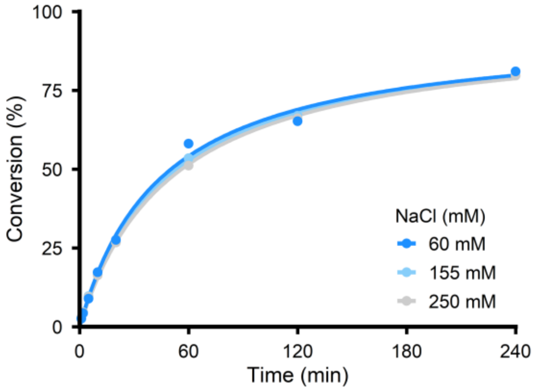

**Fig. S7.** Salt-dependent ligation kinetics in bulk solution. Conversion profiles of the ST–SC reaction measured at varying NaCl concentrations. The overall kinetic behavior remains comparable across all tested conditions, indicating that the reaction is relatively insensitive to changes in salt concentration within this range. In contrast to 155 mM standard conditions (2 samples × 2 gels), 60 mM and 250 mM data were collected from 1 sample × 2 gels.

To investigate whether variations in ionic strength affect reaction rates, we characterized the ligation kinetics of the ST–SC pair in bulk solution over a range of NaCl concentrations. As shown in **Fig. S7**, the reaction conversion profiles remain similar for 60 mM, 155 mM, and 250 mM NaCl. We observed a slight decrease in the rate constants as the salt concentration increased, with $k_{adj}$ values of 78.30, 72.98, and 68.18 $M^{-1}$ $s^{-1}$, respectively. However, such minor shifts in bulk kinetics cannot account for the dramatic acceleration of reaction rates observed in LAF condensates in low-salt environments. These results suggest that the kinetic enhancements within the condensed phase are primarily driven by specific microenvironmental factors rather than simple changes in ionic strength.

#### 1.8. Kinetic stability across varying recruiter scaffold percentages

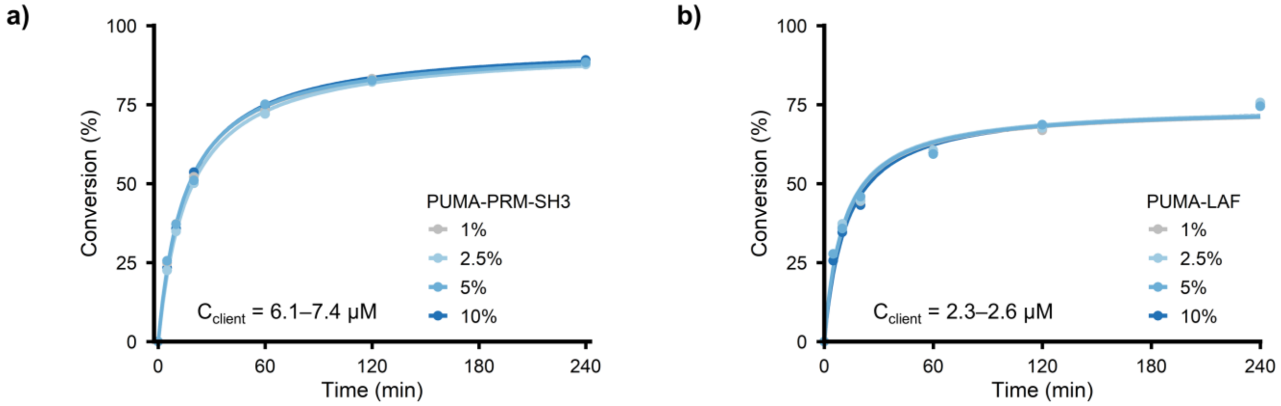

**Fig. S8.** Kinetic stability across varying recruiter scaffold percentages. **a)** Reaction conversion within PRM–SH3 condensates containing 1%, 2.5%, 5%, or 10% PUMA scaffold recruiter. **b)** Similar kinetic profiles measured within LAF condensates across the same range of PUMA scaffold percentages. The overlapping curves in both panels indicate that the reaction rate constants are insensitive to the specific fraction of the recruiter scaffold within the 1% to 10% range.

To facilitate the recruitment of clients into the condensates, we utilized a PUMA-tagged scaffold as a recruiter. To ensure that the addition of this recruiter does not fundamentally alter the internal microenvironment or the resulting reaction kinetics, we measured the ligation rates across a range of PUMA scaffold concentrations from 1% to 10%. As illustrated in **Fig. S8**, the reaction conversion profiles remain highly consistent across recruiter percentages. This stability was observed in both PRM–SH3 condensates in **a** and LAF condensates in **b**, where the client concentrations were maintained within ranges of 6.1 μM (1% PUMA) to 7.4 μM (10% PUMA) and 2.3 μM (10% PUMA) to 2.6 μM (1% PUMA), respectively. These results confirm that the recruitment strategy using a minor fraction of PUMA scaffold provides a robust platform for investigating reaction kinetics without introducing significant artifacts related to recruiter scaffold composition.

#### 1.9. Consistency of reaction rate hierarchies across varying client concentrations

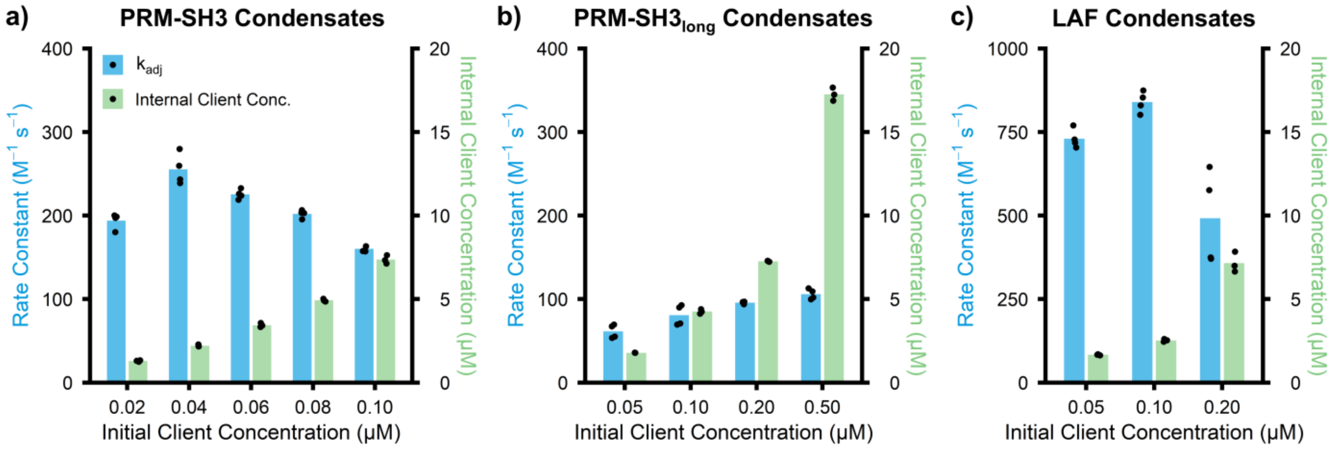

**Fig. S9.** Influence of initial and internal client concentrations on reaction rate constants. **a)** PRM–SH3, **b)** PRM–SH3$_{long}$, and **c)** LAF condensates.

To evaluate the impact of client loading on reaction efficiency, we characterized the relationship between initial and resulting internal client concentrations within the condensates, as shown in **Fig. S9**. The initial concentration represents the total amount of mCh–ST client added prior to phase separation, while the internal concentration refers to the amount recruited into the condensed phase. As expected, the internal client concentration increases with the initial concentration across all tested systems. While minor fluctuations in the rate constants are observed across different loading conditions, these variations are not significant enough to interfere with comparisons between condensate types. The clear hierarchical relationship among the reaction rates is consistently maintained, with LAF > PRM–SH3 > PRM–SH3$_{long}$ throughout the entire concentration range.

#### 1.10. Calculation of PEG excluded volume fraction

**Assuming a simple spherical shape, the excluded volume ($\varphi_{excl}$) is calculated as follows[1]:**

$\varphi_{excl}$ (%) = $(4\pi \times r_h^3 \times c \times N_A) / 3M_w$

where $r_h$ is the hydrodynamic radius of the crowder (in cm), and c is the concentration of the crowding molecule (in % w/v). Because all other parameters are constant for a given crowder, $\varphi_{excl}$ scales linearly with $c$. Therefore, the value calculated at 5% (w/v) can be scaled to other conditions by multiplying it by the corresponding concentration factor relative to 5%.

**The $r_h$ of PEG is determined by the following empirical relationship[2,3]:**

$r_h$ (nm, PEG) = $0.01912 \times M_w^{0.559}$

At a concentration of 5% (w/v), the calculated $\varphi_{excl}$ values are 5.09% for PEG400, 9.47% for PEG1000, 24.2% for PEG4000, and 38.7% for PEG8000.

**Excluded volume of Ficoll70 and Ficoll400:**

$r_h$ (Ficoll 70) = 4.06 nm, $r_h$ (Ficoll 400) = 7.26 nm[4]

At a concentration of 5% (w/v), the calculated $\varphi_{excl}$ values are 12.1% for both Ficoll70 and Ficoll400, reflecting their near-spherical geometry, which results in minimal differences in hydrodynamic radius despite the large difference in molecular weight.

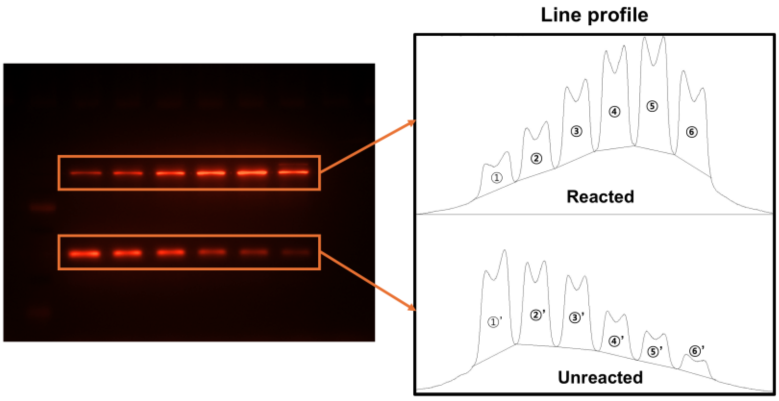

| Time (min) | Intensity (Reacted, a.u.) | Intensity (Unreacted, a.u.) | Conversion (%) |
|---|---|---|---|
| 0 | - | - | 0 |
| 5 | 13580.614 | 38869.291 | 25.89 |
| 10 | 19141.714 | 31076.016 | 38.12 |
| 20 | 29048.078 | 27185.966 | 51.66 |
| 60 | 37479.037 | 15508.915 | 70.73 |
| 120 | 43924.815 | 11856.673 | 78.74 |
| 240 | 34346.513 | 6291.137 | 84.52 |

**Fig. S10.** Representative PAGE gel and quantitative analysis of the reaction kinetics. **Left)** PAGE analysis showing the formation of reacted products (upper bands) and the depletion of unreacted clients (lower bands) over time. **Middle)** Line profiles extracted using ImageJ to quantify the fluorescence intensity of each band. Peaks 1–6 (and 1'–6') correspond to the time points from 5 to 240 min. **Right)** Summary table of the measured intensities and calculated conversion percentages. Conversion (%) is defined as the ratio of the reacted band intensity to the total band intensity.

#### 2. Supplementary Tables

**Table S1. Protein sequences of scaffolds.** The inter-motif linker, located adjacent to the His-tags, is colored purple. An underline indicates either the TEV protease site or an alpha-helical rigid linker.

| Protein names | Amino acid sequences |
|---|---|
| **PRM–SH3** | MKKKKTAPTPPKRSGGSDLNMPAYVKFNYMAEREDELSLIKGTKVIVMEKSSDGWWR<br>GSYNGQVGWFPSNYVTEEGDSPLGSENLYFQGLEHHHHHH |
| **PRM–SH3$_{long}$** | MKKKKTAPTPPKRSGGSDLNMPAYVKFNYMAEREDELSLIKGTKVIVMEKSSDGWWR<br>GSYNGQVGWFPSNYVTEEGDSPLGSEAAAKEAAAKEAAAKEAAAKEAAAKEAAAKE<br>AAAKEAAAKGSGSLEHHHHHH |
| **PUMA-PRM–SH3** | MEEQAAREIGAQLRRMADDLNAQYERGASKKKKTAPTPPKRSGGSDLNMPAYVKFN<br>YMAEREDELSLIKGTKVIVMEKSSDGWWRGSYNGQVGWFPSNYVTEEGDSPLGSENL<br>YFQGLEHHHHHH |
| **PUMA-PRM–SH3$_{long}$** | MEEQAAREIGAQLRRMADDLNAQYERGASKKKKTAPTPPKRSGGSDLNMPAYVKFN<br>YMAEREDELSLIKGTKVIVMEKSSDGWWRGSYNGQVGWFPSNYVTEEGDSPLGSEAA<br>AKEAAAKEAAAKEAAAKEAAAKEAAAKEAAAKEAAAKGSGSLEHHHHHH |
| **LAF scaffold (LAF-SUMO-His)** | MMESNQSNNGGSGNAALNRGGRYVPPHLRGGDGGAAAAASAGGDDRRGGAGGGGY<br>RRGGGNSGGGGGGGYDRGYNDNRDDRDNRGGSGGYGRDRNYEDRGYNGGGGGGG<br>NRGYNNNRGGGGGGYNRQDRGDGGSSNFSRGGYNNRDEGSDNRGSGRSYNNDRRD<br>NGGDGEFMSDSEVNQEAKPEVKPEVKPETHINLKVSDGSSEIFFKIKKTTPLRRLMEAF<br>AKRQGKEMDSLRFLYDGIRIQADQTPEDLDMEDNDIIEAHREQIGGATYLEHHHHHH |
| **TAF scaffold (His-SUMO-TAF)** | MASHHHHHHENLYFQGMSDSEVNQEAKPEVKPEVKPETHINLKVSDGSSEIFFKIKKTT<br>PLRRLMEAFAKRQGKEMDSLRFLYDGIRIQADQTPEDLDMEDNDIIEAHREQIGGATYE<br>FMSDSGSYGQSGGEQQSYSTYGNPGSQGYGQASQSYSGYGQTTDSSYGQNYSGYSSY<br>GQSQSGYSQSYGGYENQKQSSYSQQPYNNQGQQQNMESSGSQGGRAPSYDQPDYGQ<br>QDSYDQQSGYDQHQGSYDEQSNYDQQHDSYSQNQQSYHSQRENYSHHTQDDRRDVS<br>RYGEDNRGYGGSQGGGRGRGGYDKDGRGPMTGSSGGDRGG |
| **FUS scaffold (His-SUMO-FUS)** | MASHHHHHHENLYFQGMSDSEVNQEAKPEVKPEVKPETHINLKVSDGSSEIFFKIKKTT<br>PLRRLMEAFAKRQGKEMDSLRFLYDGIRIQADQTPEDLDMEDNDIIEAHREQIGGATYE<br>FASNDYTQQATQSYGAYPTQPGQGYSQQSSQPYGQQSYSGYSQSTDTSGYGQSSYSSY<br>GQSQNTGYGTQSTPQGYGSTGGYGSSQSSQSSYGQQSSYPGYGQQPAPSSTSGSYGSSS<br>QSSSYGQPQSGSYSQQPSYGGQQQSYGQQQSYNPPQGYGQQNQYNSSSGGGGGGGGG<br>GNYGQDQSSMSSGGGSGGGYGNQDQSGGGGSGGYGQQDRG |
| **PUMA-LAF-SUMO-His** | MEEQAAREIGAQLRRMADDLNAQYERGASMESNQSNNGGSGNAALNRGGRYVPPHL<br>RGGDGGAAAAASAGGDDRRGGAGGGGYRRGGGNSGGGGGGGYDRGYNDNRDDRD<br>NRGGSGGYGRDRNYEDRGYNGGGGGGGNRGYNNNRGGGGGGYNRQDRGDGGSSNF<br>SRGGYNNRDEGSDNRGSGRSYNNDRRDNGGDGEFMSDSEVNQEAKPEVKPEVKPETH<br>INLKVSDGSSEIFFKIKKTTPLRRLMEAFAKRQGKEMDSLRFLYDGIRIQADQTPEDLDM<br>EDNDIIEAHREQIGGATYLEHHHHHH |
| **PUMA-FUS-SUMO-His** | MEEQAAREIGAQLRRMADDLNAQYERGASASNDYTQQATQSYGAYPTQPGQGYSQQ<br>SSQPYGQQSYSGYSQSTDTSGYGQSSYSSYGQSQNTGYGTQSTPQGYGSTGGYGSSQS<br>SQSSYGQQSSYPGYGQQPAPSSTSGSYGSSSQSSSYGQPQSGSYSQQPSYGGQQQSYGQ<br>QQSYNPPQGYGQQNQYNSSSGGGGGGGGGGNYGQDQSSMSSGGGSGGGYGNQDQS<br>GGGGSGGYGQQDRGEFMSDSEVNQEAKPEVKPEVKPETHINLKVSDGSSEIFFKIKKTT<br>PLRRLMEAFAKRQGKEMDSLRFLYDGIRIQADQTPEDLDMEDNDIIEAHREQIGGATYL<br>EHHHHHH |

**Table S2. Protein sequences of clients.** The slash (/) denotes client proteins used after His-tag removal by TEV protease cleavage. The TEV protease recognition site is underlined.

| Protein names | Amino acid sequences |
|---|---|
| **Common Bcl domain of clients (Bcl)** | MSAMSQSNRELVVDFLSYKLSQKGYSWSQFSDVEENRTEAPEGTESEAVKQALREAG<br>DEFELRYRRAFSDLTSQLHITPGTAYQSFEQVVNELFRDGVNWGRIVAFFSFGGALSVE<br>SVDKEMQVLVSRIAAWMATYLNDHLEPWIQENGGWDTFVELYGNNAAAESRKGQER<br>LEENLYFQ/GLEHHHHHH |
| **mCh-SpyTag-Bcl** | MVSKGEEDNMAIIKEFMRFKVHMEGSVNGHEFEIEGEGEGRPYEGTQTAKLKVTKGGP<br>LPFAWDILSPQFMYGSKAYVKHPADIPDYLKLSFPEGFKWERVMNFEDGGVVTVTQDS<br>SLQDGEFIYKVKLRGTNFPSDGPVMQKKTMGWEASSERMYPEDGALKGEIKQRLKLK<br>DGGHYDAEVKTTYKAKKPVQLPGAYNVNIKLDITSHNEDYTIVEQYERAEGRHSTGG<br>MDELYKGGSGGSAHIVMVDAYKPTKGGSGS(Bcl) |
| **GFP-SpyCatcher-Bcl** | MKGEELFTGVVPILVELDGDVNGHEFSVRGEGEGDATIGKLTLKFICTTGKLPVPWPTL<br>VTTLTYGVQCFSRYPDHMKRHDFFKSAMPEGYVQERTISFKDDGKYKTRAVVKFEGD<br>TLVNRIELKGTDFKEDGNILGHKLEYNFNSHDVYITADKQENGIKAEFTVRHNVEDGSV<br>QLADHYQQNTPIGDGPVLLPDNHYLSTQTVLSKDPNEKRDHMVLHEYVNAAGITGSG<br>AMVDTLSGLSSEQGQSGDMTIEEDSATHIKFSKRDEDGKELAGATMELRDSSGKTIST<br>WISDGQVKDFYLYPGKYTFVETAAPDGYEVATAITFTVNEQGQVTVNGKATKGDAHI<br>GSGS(Bcl) |
| **mCh-SpyTag003-Bcl** | MVSKGEEDNMAIIKEFMRFKVHMEGSVNGHEFEIEGEGEGRPYEGTQTAKLKVTKGGP<br>LPFAWDILSPQFMYGSKAYVKHPADIPDYLKLSFPEGFKWERVMNFEDGGVVTVTQDS<br>SLQDGEFIYKVKLRGTNFPSDGPVMQKKTMGWEASSERMYPEDGALKGEIKQRLKLK<br>DGGHYDAEVKTTYKAKKPVQLPGAYNVNIKLDITSHNEDYTIVEQYERAEGRHSTGG<br>MDELYKGGSGGSRGVPHIVMVDAYKRYKGGSGS(Bcl) |
| **GFP-SpyCatcher003-Bcl** | MKGEELFTGVVPILVELDGDVNGHEFSVRGEGEGDATIGKLTLKFICTTGKLPVPWPTL<br>VTTLTYGVQCFSRYPDHMKRHDFFKSAMPEGYVQERTISFKDDGKYKTRAVVKFEGD<br>TLVNRIELKGTDFKEDGNILGHKLEYNFNSHDVYITADKQENGIKAEFTVRHNVEDGSV<br>QLADHYQQNTPIGDGPVLLPDNHYLSTQTVLSKDPNEKRDHMVLHEYVNAAGITGSG<br>AMVTTLSGLSGEQGPSGDMTTEEDSATHIKFSKRDEDGRELAGATMELRDSSGKTIST<br>WISDGHVKDFYLYPGKYTFVETAAPDGYEVATPIEFTVNEDGQVTVDGEATEGDAHT<br>GSGS(Bcl) |
| **mCerulean-Bcl** | MMVSKGEELFTGVVPILVELDGDVNGHKFSVSGEGEGDATYGKLTLKFICTTGKLPVP<br>WPTLVTTLSWGVQCFARYPDHMKQHDFFKSAMPEGYVQERTIFFKDDGNYKTRAEVK<br>FEGDTLVNRIELKGIDFKEDGNILGHKLEYNAIHGNVYITADKQKNGIKANFGLNCNIE<br>DGSVQLADHYQQNTPIGDGPVLLPDNHYLSTQSKLSKDPNEKRDHMVLLEFVTAAGIT<br>LGMDELYKGS(Bcl) |
| **mCitrine- Bcl** | MMVSKGEELFTGVVPILVELDGDVNGHKFSVSGEGEGDATYGKLTLKFICTTGKLPVP<br>WPTLVTTFGYGLMCFARYPDHMKQHDFFKSAMPEGYVQERTIFFKDDGNYKTRAEVK<br>FEGDTLVNRIELKGIDFKEDGNILGHKLEYNYNSHNVYIMADKQKNGIKVNFKIRHNIE<br>DGSVQLADHYQQNTPIGDGPVLLPDNHYLSYQSALSKDPNEKRDHMVLLEFVTAAGIT<br>LGMDELYKGS(Bcl) |

#### 3. Data Collection of Reaction Progress

The following datasets are the source data for reaction progress and the derived kinetic parameters for each experimental condition. In these tables, "Client Concentration" refers to the internal client concentration in the condensates, quantified by confocal laser microscopy, while the "GFP client ratio" denotes the initial input ratio, specifically adjusted to achieve an equimolar internal distribution of both clients. The results—including conversion percentages at time points, apparent rate constant $k_{obs}$, rate constant fitted using a modified equation ($k_{adj}$), and unreacted fractions—represent the mean values derived from the 2 × 2 matrix of independent samples and gels and consistently reflect the high reproducibility of the experimental measurements. For bulk client solutions at low concentrations, the reaction rates were too slow to be accurately modeled by the adjusted second-order fitting; therefore, only $k_{obs}$ values were calculated using standard second-order fitting.

##### - PRM-SH3$_{short}$ Scaffold

**PRM-SH3$_{short}$ (100 µM; 1% PUMA), 0.1 µM mCh-ST-Bcl: 197.67 M$^{-1}$ s$^{-1}$**

| Client Concentration = 6.067 µM | Conversion (%) | | | | | | | | | | |
|---|---|---|---|---|---|---|---|---|---|---|---|
| GFP client ratio = 0.7 [mCh] | Sample 1 - Gel 1 | | Sample 1 - Gel 2 | | Sample 2 - Gel 1 | | Sample 2 - Gel 2 | | Average | | |
| Reaction Time (min) | mCh-ST | GFP-SC | mCh-ST | GFP-SC | mCh-ST | GFP-SC | mCh-ST | GFP-SC | mCh-ST | GFP-SC | Total |
| 5 | 25.83 | 23.05 | 26.70 | 22.09 | 24.37 | 23.59 | 27.44 | 26.19 | 26.08 | 23.73 | 24.91 |
| 10 | 37.74 | 34.51 | 37.53 | 34.63 | 38.07 | 37.35 | 39.39 | 38.52 | 38.18 | 36.25 | 37.22 |
| 20 | 54.53 | 49.20 | 53.28 | 49.73 | 51.00 | 52.09 | 53.20 | 53.97 | 53.01 | 51.25 | 52.13 |
| 60 | 74.78 | 72.37 | 73.01 | 70.96 | 69.88 | 75.28 | 71.85 | 76.03 | 72.38 | 73.66 | 73.02 |
| 120 | 84.83 | 83.14 | 83.18 | 82.27 | 78.54 | 86.75 | 79.67 | 87.04 | 81.56 | 84.80 | 83.18 |
| 240 | 89.60 | 87.61 | 88.10 | 87.01 | 84.26 | 92.51 | 85.28 | 92.66 | 86.81 | 89.95 | 88.38 |
| $k_{obs}$ (M^-1 sec^-1) | 159 | 133 | 152 | 130 | 134 | 154 | 150 | 166 | 149 | 146 | 147 |
| $k_{adj}$ (M^-1 sec^-1) | 199 | 170 | 210 | 173 | 230 | 165 | 252 | 183 | 223 | 173 | 198 |
| Unreacted Fracrtion (%) | 5.47 | 6.01 | 7.67 | 7.07 | 12.39 | 1.63 | 11.88 | 2.39 | 9.35 | 4.28 | 6.81 |

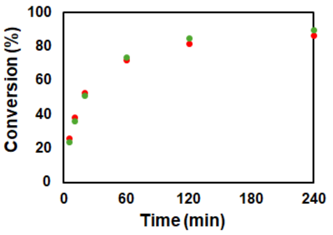

**PRM-SH3$_{short}$ (100 µM; 2.5% PUMA), 0.1 µM mCh-ST-Bcl: 161.61 M$^{-1}$ s$^{-1}$**

| Client Concentration = 6.554 µM | Conversion (%) | | | | | | | | | | |
|---|---|---|---|---|---|---|---|---|---|---|---|
| GFP client ratio = 0.7 [mCh] | Sample 1 - Gel 1 | | Sample 1 - Gel 2 | | Sample 2 - Gel 1 | | Sample 2 - Gel 2 | | Average | | |
| Reaction Time (min) | mCh-ST | GFP-SC | mCh-ST | GFP-SC | mCh-ST | GFP-SC | mCh-ST | GFP-SC | mCh-ST | GFP-SC | Total |
| 5 | 21.92 | 20.00 | 22.89 | 20.59 | 23.88 | 21.68 | 25.95 | 22.98 | 23.66 | 21.31 | 22.49 |
| 10 | 36.18 | 32.52 | 37.17 | 32.72 | 36.67 | 33.45 | 36.98 | 33.17 | 36.75 | 32.97 | 34.86 |
| 20 | 53.69 | 48.06 | 54.45 | 48.22 | 51.73 | 46.73 | 51.56 | 46.27 | 52.86 | 47.32 | 50.09 |
| 60 | 73.99 | 69.58 | 73.49 | 68.50 | 74.26 | 71.44 | 73.91 | 71.18 | 73.91 | 70.17 | 72.04 |
| 120 | 84.35 | 80.63 | 83.42 | 79.85 | 83.02 | 81.72 | 82.84 | 81.61 | 83.41 | 80.95 | 82.18 |
| 240 | 88.94 | 85.23 | 88.85 | 84.79 | 88.27 | 88.22 | 88.41 | 88.69 | 88.62 | 86.73 | 87.68 |
| $k_{obs}$ (M^-1 sec^-1) | 136.17 | 108.73 | 138.98 | 107.56 | 134.65 | 114.08 | 136.92 | 114.03 | 136.68 | 111.10 | 123.89 |
| $k_{adj}$ (M^-1 sec^-1) | 166.00 | 151.08 | 178.24 | 157.55 | 173.83 | 142.15 | 182.13 | 141.92 | 175.05 | 148.17 | 161.61 |
| Unreacted Fracrtion (%) | 4.95 | 8.06 | 6.12 | 9.22 | 6.27 | 5.47 | 6.90 | 5.41 | 6.06 | 7.04 | 6.55 |

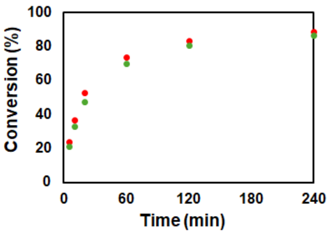

**PRM-SH3$_{short}$ (100 µM; 5% PUMA), 0.1 µM mCh-ST-Bcl: 160.17 M$^{-1}$ s$^{-1}$**

| Client Concentration = 7.360 µM | Conversion (%) | | | | | | | | | | |
|---|---|---|---|---|---|---|---|---|---|---|---|
| GFP client ratio = 0.7 [mCh] | Sample 1 - Gel 1 | | Sample 1 - Gel 2 | | Sample 2 - Gel 1 | | Sample 2 - Gel 2 | | Average | | |
| Reaction Time (min) | mCh-ST | GFP-SC | mCh-ST | GFP-SC | mCh-ST | GFP-SC | mCh-ST | GFP-SC | mCh-ST | GFP-SC | Total |
| 5 | 26.97 | 23.30 | 27.55 | 21.63 | 28.09 | 25.01 | 27.26 | 24.54 | 27.47 | 23.62 | 25.54 |
| 10 | 38.76 | 34.34 | 38.95 | 34.71 | 39.23 | 35.47 | 39.34 | 36.25 | 39.07 | 35.19 | 37.13 |
| 20 | 53.74 | 49.43 | 55.44 | 49.95 | 51.75 | 47.64 | 51.79 | 48.25 | 53.18 | 48.82 | 51.00 |
| 60 | 76.63 | 73.32 | 77.48 | 74.04 | 76.16 | 73.26 | 76.30 | 73.39 | 76.64 | 73.50 | 75.07 |
| 120 | 83.24 | 80.46 | 84.30 | 80.77 | 84.29 | 81.69 | 84.53 | 81.94 | 84.09 | 81.21 | 82.65 |
| 240 | 89.17 | 85.97 | 89.37 | 86.39 | 90.17 | 87.76 | 90.54 | 87.42 | 89.81 | 86.89 | 88.35 |
| $k_{obs}$ (M^-1 sec^-1) | 134.04 | 108.28 | 140.42 | 109.24 | 133.36 | 110.37 | 133.08 | 112.12 | 135.22 | 110.00 | 122.61 |
| $k_{adj}$ (M^-1 sec^-1) | 174.52 | 151.75 | 178.66 | 147.42 | 168.07 | 146.67 | 163.69 | 150.56 | 171.24 | 149.10 | 160.17 |
| Unreacted Fracrtion (%) | 6.40 | 8.20 | 5.87 | 7.37 | 5.60 | 6.91 | 5.05 | 7.15 | 5.73 | 7.41 | 6.57 |

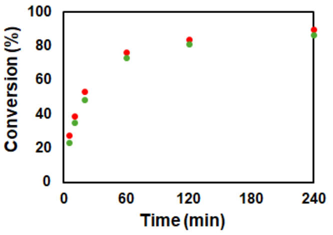

#### PRM-SH3$_{short}$ (100 µM; 10% PUMA), 0.1 µM mCh-ST-Bcl: 151.96 $M^{-1}$ $s^{-1}$

| Client Concentration = 7.405 µM | Conversion (%) | | | | | | | | | | |
|---|---|---|---|---|---|---|---|---|---|---|---|
| GFP client ratio = 0.65 [mCh] | Sample 1 - Gel 1 | | Sample 1 - Gel 2 | | Sample 2 - Gel 1 | | Sample 2 - Gel 2 | | Average | | |
| Reaction Time (min) | mCh-ST | GFP-SC | mCh-ST | GFP-SC | mCh-ST | GFP-SC | mCh-ST | GFP-SC | mCh-ST | GFP-SC | Total |
| 5 | 26.33 | 21.97 | 25.91 | 20.93 | 24.58 | 21.40 | 24.97 | 19.83 | 25.45 | 21.03 | 23.24 |
| 10 | 38.44 | 33.01 | 38.38 | 32.25 | 37.78 | 34.42 | 37.13 | 34.58 | 37.93 | 33.57 | 35.75 |
| 20 | 57.91 | 51.06 | 57.67 | 52.23 | 53.92 | 50.91 | 53.86 | 51.22 | 55.84 | 51.36 | 53.60 |
| 60 | 76.75 | 70.43 | 77.73 | 70.16 | 75.79 | 75.22 | 76.16 | 75.71 | 76.61 | 72.88 | 74.74 |
| 120 | 85.71 | 81.09 | 82.34 | 81.39 | 83.33 | 84.14 | 81.46 | 85.07 | 83.21 | 82.92 | 83.07 |
| 240 | 91.75 | 86.05 | 88.56 | 87.32 | 89.32 | 90.61 | 88.51 | 90.74 | 89.53 | 88.68 | 89.11 |
| $k_{obs}$ (M^-1 sec^-1) | 142.89 | 104.30 | 138.87 | 104.60 | 127.99 | 115.09 | 125.94 | 115.51 | 133.92 | 109.87 | 121.90 |
| $k_{adj}$ (M^-1 sec^-1) | 166.64 | 145.76 | 183.88 | 138.78 | 161.59 | 127.91 | 167.46 | 123.62 | 169.89 | 134.02 | 151.96 |
| Unreacted Fracrtion (%) | 3.83 | 8.15 | 6.85 | 6.97 | 5.74 | 2.70 | 6.95 | 1.76 | 5.84 | 4.90 | 5.37 |

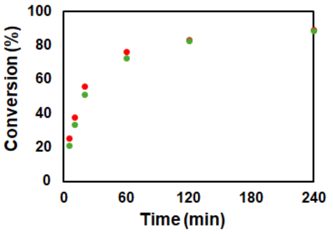

#### PRM-SH3$_{short}$ (100 µM; 5% PUMA), 0.08 µM mCh-ST-Bcl: 202.04 $M^{-1}$ $s^{-1}$

| Client Concentration = 4.924 µM | Conversion (%) | | | | | | | | | | |
|---|---|---|---|---|---|---|---|---|---|---|---|
| GFP client ratio = 0.65 [mCh] | Sample 1 - Gel 1 | | Sample 1 - Gel 2 | | Sample 2 - Gel 1 | | Sample 2 - Gel 2 | | Average | | |
| Reaction Time (min) | mCh-ST | GFP-SC | mCh-ST | GFP-SC | mCh-ST | GFP-SC | mCh-ST | GFP-SC | mCh-ST | GFP-SC | Total |
| 5 | 19.61 | 17.32 | 20.61 | 18.32 | 25.33 | 20.89 | 25.00 | 20.37 | 22.64 | 19.23 | 20.93 |
| 10 | 33.23 | 30.07 | 34.24 | 30.88 | 35.50 | 29.86 | 36.16 | 30.58 | 34.78 | 30.35 | 32.57 |
| 20 | 48.34 | 44.54 | 50.07 | 46.01 | 50.77 | 43.55 | 51.13 | 44.38 | 50.08 | 44.62 | 47.35 |
| 60 | 70.37 | 67.92 | 71.68 | 69.74 | 74.06 | 67.19 | 74.83 | 67.34 | 72.74 | 68.05 | 70.39 |
| 120 | 78.62 | 78.37 | 79.17 | 78.53 | 84.60 | 80.83 | 85.43 | 80.77 | 81.95 | 79.63 | 80.79 |
| 240 | 83.88 | 84.70 | 85.20 | 84.58 | 88.32 | 82.57 | 89.11 | 83.13 | 86.63 | 83.74 | 85.19 |
| $k_{obs}$ (M^-1 sec^-1) | 144.22 | 126.24 | 154.60 | 133.87 | 178.13 | 127.74 | 182.82 | 130.26 | 164.94 | 129.53 | 147.23 |
| $k_{adj}$ (M^-1 sec^-1) | 217.19 | 173.59 | 225.65 | 187.29 | 222.25 | 184.44 | 219.02 | 186.90 | 221.03 | 183.05 | 202.04 |
| Unreacted Fracrtion (%) | 9.89 | 7.89 | 9.16 | 8.28 | 5.45 | 8.96 | 4.49 | 8.81 | 7.25 | 8.48 | 7.87 |

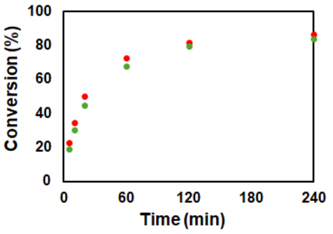

#### PRM-SH3$_{short}$ (100 µM; 5% PUMA), 0.06 µM mCh-ST-Bcl: 225.34 $M^{-1}$ $s^{-1}$

| Client Concentration = 3.432 µM | Conversion (%) | | | | | | | | | | |
|---|---|---|---|---|---|---|---|---|---|---|---|
| GFP client ratio = 0.64 [mCh] | Sample 1 - Gel 1 | | Sample 1 - Gel 2 | | Sample 2 - Gel 1 | | Sample 2 - Gel 2 | | Average | | |
| Reaction Time (min) | mCh-ST | GFP-SC | mCh-ST | GFP-SC | mCh-ST | GFP-SC | mCh-ST | GFP-SC | mCh-ST | GFP-SC | Total |
| 5 | 19.55 | 17.45 | 20.74 | 18.77 | 20.12 | 14.36 | 20.34 | 12.96 | 20.19 | 15.88 | 18.04 |
| 10 | 30.46 | 25.94 | 32.39 | 26.17 | 26.45 | 25.88 | 26.24 | 25.73 | 28.88 | 25.93 | 27.40 |
| 20 | 43.32 | 38.49 | 44.31 | 38.82 | 40.59 | 41.81 | 40.59 | 41.44 | 42.20 | 40.14 | 41.17 |
| 60 | 66.39 | 61.32 | 67.90 | 63.81 | 60.05 | 63.90 | 59.80 | 63.57 | 63.53 | 63.15 | 63.34 |
| 120 | 77.74 | 74.14 | 78.51 | 75.69 | 69.61 | 76.25 | 69.82 | 76.63 | 73.92 | 75.68 | 74.80 |
| 240 | 86.29 | 83.82 | 87.45 | 84.70 | 81.08 | 87.42 | 81.08 | 88.26 | 83.97 | 86.05 | 85.01 |
| $k_{obs}$ (M^-1 sec^-1) | 179.03 | 141.03 | 192.96 | 150.91 | 136.78 | 155.40 | 136.51 | 153.65 | 161.32 | 150.25 | 155.78 |
| $k_{adj}$ (M^-1 sec^-1) | 245.67 | 202.12 | 260.87 | 204.28 | 266.44 | 185.58 | 264.92 | 172.88 | 259.47 | 191.21 | 225.34 |
| Unreacted Fracrtion (%) | 7.75 | 8.86 | 7.36 | 7.53 | 15.28 | 4.55 | 15.20 | 3.07 | 11.40 | 6.00 | 8.70 |

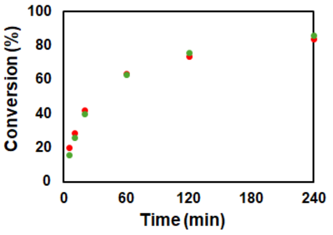

#### PRM-SH3$_{short}$ (100 µM; 5% PUMA), 0.04 µM mCh-ST-Bcl: 255.32 $M^{-1}$ $s^{-1}$

| Client Concentration = 2.204 µM | Conversion (%) | | | | | | | | | | |
|---|---|---|---|---|---|---|---|---|---|---|---|
| GFP client ratio = 0.63 [mCh] | Sample 1 - Gel 1 | | Sample 1 - Gel 2 | | Sample 2 - Gel 1 | | Sample 2 - Gel 2 | | Average | | |
| Reaction Time (min) | mCh-ST | GFP-SC | mCh-ST | GFP-SC | mCh-ST | GFP-SC | mCh-ST | GFP-SC | mCh-ST | GFP-SC | Total |
| 5 | 13.79 | 12.45 | 14.36 | 12.16 | 14.87 | 13.76 | 13.73 | 14.54 | 14.19 | 13.23 | 13.71 |
| 10 | 22.28 | 21.74 | 23.28 | 20.86 | 20.92 | 21.37 | 20.94 | 20.99 | 21.86 | 21.24 | 21.55 |
| 20 | 34.76 | 32.55 | 33.22 | 32.84 | 34.60 | 33.56 | 33.80 | 32.01 | 34.10 | 32.74 | 33.42 |
| 60 | 60.43 | 60.30 | 61.08 | 61.36 | 57.22 | 57.56 | 57.60 | 56.81 | 59.08 | 59.01 | 59.05 |
| 120 | 72.31 | 75.19 | 72.29 | 75.07 | 68.84 | 69.47 | 69.31 | 70.22 | 70.69 | 72.49 | 71.59 |
| 240 | 79.77 | 83.70 | 80.43 | 84.03 | 76.83 | 79.96 | 78.59 | 79.19 | 78.91 | 81.72 | 80.31 |
| $k_{obs}$ (M^-1 sec^-1) | 187.16 | 190.02 | 188.28 | 191.71 | 167.36 | 170.87 | 168.80 | 166.57 | 177.90 | 179.79 | 178.85 |
| $k_{adj}$ (M^-1 sec^-1) | 275.22 | 211.25 | 269.09 | 208.42 | 300.96 | 258.49 | 270.13 | 248.99 | 278.85 | 231.79 | 255.32 |
| Unreacted Fracrtion (%) | 9.72 | 2.84 | 9.04 | 2.26 | 14.22 | 10.38 | 11.69 | 10.14 | 11.17 | 6.41 | 8.79 |

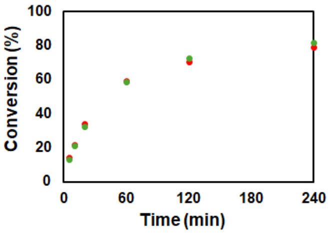

#### PRM-SH3$_{short}$ (100 µM; 5% PUMA), 0.02 µM mCh-ST-Bcl: 193.95 $M^{-1}$ $s^{-1}$

| Client Concentration = 1.293 µM | Conversion (%) | | | | | | | | | | |
|---|---|---|---|---|---|---|---|---|---|---|---|
| GFP client ratio = 0.62 [mCh] | Sample 1 - Gel 1 | | Sample 1 - Gel 2 | | Sample 2 - Gel 1 | | Sample 2 - Gel 2 | | Average | | |
| Reaction Time (min) | mCh-ST | GFP-SC | mCh-ST | GFP-SC | mCh-ST | GFP-SC | mCh-ST | GFP-SC | mCh-ST | GFP-SC | Total |
| 5 | 6.91 | 7.42 | 6.21 | 6.83 | 8.91 | 8.19 | 9.10 | 8.40 | 7.79 | 7.71 | 7.75 |
| 10 | 12.76 | 11.09 | 12.38 | 11.39 | 12.85 | 11.83 | 12.66 | 12.64 | 12.66 | 11.74 | 12.20 |
| 20 | 21.74 | 19.55 | 20.70 | 19.72 | 20.76 | 19.62 | 20.49 | 20.20 | 20.92 | 19.77 | 20.35 |
| 60 | 44.09 | 39.93 | 41.24 | 39.76 | 37.54 | 37.82 | 39.63 | 38.79 | 40.62 | 39.08 | 39.85 |
| 120 | 60.17 | 58.93 | 58.05 | 59.21 | 53.72 | 56.27 | 53.98 | 55.85 | 56.48 | 57.56 | 57.02 |
| 240 | 70.89 | 72.69 | 70.79 | 73.45 | 68.22 | 70.63 | 69.59 | 71.76 | 69.87 | 72.13 | 71.00 |
| $k_{obs}$ (M^-1 sec^-1) | 161.75 | 149.64 | 149.36 | 151.22 | 131.69 | 138.41 | 137.66 | 142.41 | 145.12 | 145.42 | 145.27 |
| $k_{adj}$ (M^-1 sec^-1) | 236.81 | 163.41 | 203.66 | 156.49 | 220.35 | 173.93 | 217.64 | 179.28 | 219.61 | 168.28 | 193.95 |
| Unreacted Fracrtion (%) | 10.58 | 2.66 | 8.86 | 1.05 | 14.21 | 6.73 | 12.73 | 6.72 | 11.59 | 4.29 | 7.94 |

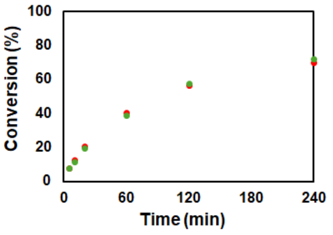

### - PRM-SH3$_{long}$ Scaffold

#### PRM-SH3$_{long}$ (100 μM; 5% PUMA), 0.5 μM mCh-ST-Bcl: 105.85 $M^{-1}$ $s^{-1}$

| Client Concentration = 17.255 μM | Conversion (%) | | | | | | | | | | |
|---|---|---|---|---|---|---|---|---|---|---|---|
| GFP client ratio = 0.8 [mCh] | Sample 1 - Gel 1 | | Sample 1 - Gel 2 | | Sample 2 - Gel 1 | | Sample 2 - Gel 2 | | Average | | |
| Reaction Time (min) | mCh-ST | GFP-SC | mCh-ST | GFP-SC | mCh-ST | GFP-SC | mCh-ST | GFP-SC | mCh-ST | GFP-SC | Total |
| 5 | 31.24 | 26.96 | 31.44 | 26.87 | 34.74 | 29.04 | 35.83 | 29.10 | 33.31 | 27.99 | 30.65 |
| 10 | 45.71 | 39.71 | 45.24 | 39.85 | 49.57 | 42.49 | 50.73 | 43.25 | 47.81 | 41.32 | 44.57 |
| 20 | 62.39 | 56.03 | 62.16 | 56.22 | 65.44 | 57.43 | 66.26 | 58.24 | 64.06 | 56.98 | 60.52 |
| 60 | 79.66 | 75.94 | 80.10 | 76.74 | 82.78 | 75.39 | 83.37 | 75.74 | 81.48 | 75.95 | 78.71 |
| 120 | 83.29 | 80.67 | 83.76 | 81.22 | 88.57 | 81.95 | 88.92 | 82.33 | 86.13 | 81.54 | 83.84 |
| 240 | 86.93 | 84.52 | 87.09 | 85.04 | 90.35 | 84.57 | 90.68 | 84.68 | 88.76 | 84.70 | 86.73 |
| $k_{obs}$ (M^-1 sec^-1) | 75.59 | 57.53 | 75.50 | 58.50 | 91.71 | 62.45 | 96.12 | 64.26 | 84.73 | 60.69 | 72.71 |
| $k_{adj}$ (M^-1 sec^-1) | 112.40 | 91.11 | 110.02 | 89.43 | 116.21 | 102.13 | 121.28 | 104.23 | 114.98 | 96.73 | 105.85 |
| Unreacted Fracrtion (%) | 9.24 | 10.78 | 8.81 | 10.03 | 5.58 | 11.32 | 5.45 | 11.15 | 7.27 | 10.82 | 9.04 |

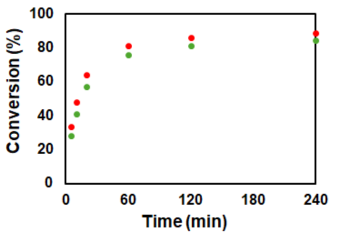

#### PRM-SH3$_{long}$ (100 μM; 5% PUMA), 0.2 μM mCh-ST-Bcl: 95.68 $M^{-1}$ $s^{-1}$

| Client Concentration = 7.258 μM | Conversion (%) | | | | | | | | | | |
|---|---|---|---|---|---|---|---|---|---|---|---|
| GFP client ratio = 0.75 [mCh] | Sample 1 - Gel 1 | | Sample 1 - Gel 2 | | Sample 2 - Gel 1 | | Sample 2 - Gel 2 | | Average | | |
| Reaction Time (min) | mCh-ST | GFP-SC | mCh-ST | GFP-SC | mCh-ST | GFP-SC | mCh-ST | GFP-SC | mCh-ST | GFP-SC | Total |
| 5 | 15.80 | 14.20 | 15.49 | 14.19 | 17.57 | 15.11 | 17.88 | 15.74 | 16.69 | 14.81 | 15.75 |
| 10 | 25.03 | 22.73 | 24.51 | 22.77 | 26.81 | 22.88 | 27.21 | 23.43 | 25.89 | 22.95 | 24.42 |
| 20 | 40.96 | 37.65 | 40.68 | 37.93 | 41.16 | 36.36 | 41.74 | 37.10 | 41.14 | 37.26 | 39.20 |
| 60 | 64.24 | 64.35 | 63.98 | 64.52 | 65.60 | 61.35 | 66.14 | 61.59 | 64.99 | 62.95 | 63.97 |
| 120 | 72.44 | 76.08 | 72.58 | 75.55 | 76.23 | 73.85 | 76.79 | 74.15 | 74.51 | 74.91 | 74.71 |
| 240 | 79.81 | 84.22 | 79.39 | 84.35 | 83.72 | 82.80 | 83.93 | 82.69 | 81.71 | 83.52 | 82.61 |
| $k_{obs}$ (M^-1 sec^-1) | 67.38 | 66.58 | 66.33 | 66.70 | 75.14 | 61.61 | 77.19 | 63.07 | 71.51 | 64.49 | 68.00 |
| $k_{adj}$ (M^-1 sec^-1) | 113.00 | 79.79 | 111.02 | 80.79 | 105.31 | 82.00 | 107.38 | 86.13 | 109.18 | 82.18 | 95.68 |
| Unreacted Fracrtion (%) | 12.50 | 4.73 | 12.48 | 4.99 | 8.37 | 7.28 | 8.19 | 7.86 | 10.38 | 6.22 | 8.30 |

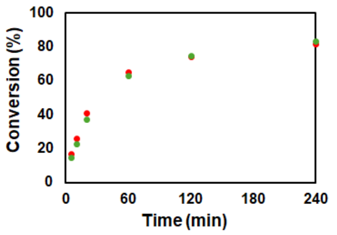

#### PRM-SH3$_{long}$ (100 μM; 5% PUMA), 0.1 μM mCh-ST-Bcl: 80.70 $M^{-1}$ $s^{-1}$

| Client Concentration = 4.252 μM | Conversion (%) | | | | | | | | | | |
|---|---|---|---|---|---|---|---|---|---|---|---|
| GFP client ratio = 0.66 [mCh] | Sample 1 - Gel 1 | | Sample 1 - Gel 2 | | Sample 2 - Gel 1 | | Sample 2 - Gel 2 | | Average | | |
| Reaction Time (min) | mCh-ST | GFP-SC | mCh-ST | GFP-SC | mCh-ST | GFP-SC | mCh-ST | GFP-SC | mCh-ST | GFP-SC | Total |
| 5 | 8.60 | 8.21 | 7.27 | 7.16 | 8.24 | 6.24 | 7.33 | 6.26 | 7.86 | 6.97 | 7.41 |
| 10 | 15.33 | 14.20 | 14.14 | 14.02 | 13.98 | 12.28 | 13.66 | 12.29 | 14.27 | 13.20 | 13.74 |
| 20 | 23.46 | 22.88 | 23.22 | 23.67 | 22.39 | 19.90 | 23.44 | 20.07 | 23.13 | 21.63 | 22.38 |
| 60 | 46.89 | 45.45 | 47.63 | 48.10 | 51.21 | 47.22 | 53.53 | 48.89 | 49.81 | 47.41 | 48.61 |
| 120 | 57.00 | 58.62 | 57.91 | 59.88 | 63.17 | 59.64 | 62.50 | 59.33 | 60.15 | 59.37 | 59.76 |
| 240 | 69.84 | 70.75 | 67.98 | 74.28 | 75.63 | 72.65 | 77.77 | 74.41 | 72.81 | 73.02 | 72.91 |
| $k_{obs}$ (M^-1 sec^-1) | 50.18 | 50.10 | 49.67 | 54.85 | 59.81 | 50.76 | 62.65 | 52.66 | 55.58 | 52.09 | 53.84 |
| $k_{adj}$ (M^-1 sec^-1) | 100.11 | 85.23 | 102.25 | 77.58 | 75.40 | 66.17 | 74.29 | 64.56 | 88.01 | 73.38 | 80.70 |
| Unreacted Fracrtion (%) | 17.65 | 14.12 | 18.46 | 9.49 | 6.47 | 7.55 | 4.80 | 5.85 | 11.85 | 9.25 | 10.55 |

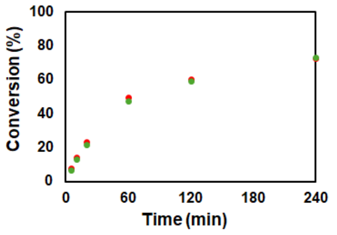

#### PRM-SH3$_{long}$ (100 μM; 5% PUMA), 0.05 μM mCh-ST-Bcl: 61.27 $M^{-1}$ $s^{-1}$

| Client Concentration = 1.784 μM | Conversion (%) | | | | | | | | | | |
|---|---|---|---|---|---|---|---|---|---|---|---|
| GFP client ratio = 0.6 [mCh] | Sample 1 - Gel 1 | | Sample 1 - Gel 2 | | Sample 2 - Gel 1 | | Sample 2 - Gel 2 | | Average | | |
| Reaction Time (min) | mCh-ST | GFP-SC | mCh-ST | GFP-SC | mCh-ST | GFP-SC | mCh-ST | GFP-SC | mCh-ST | GFP-SC | Total |
| 10 | 5.93 | 5.60 | 5.47 | 5.65 | 6.05 | 5.64 | 5.02 | 5.65 | 5.62 | 5.64 | 5.63 |
| 20 | 11.02 | 11.33 | 11.91 | 11.11 | 10.70 | 9.64 | 10.81 | 9.88 | 11.11 | 10.49 | 10.80 |
| 60 | 27.04 | 26.65 | 28.53 | 26.57 | 26.56 | 24.89 | 25.48 | 24.50 | 26.90 | 25.65 | 26.28 |
| 120 | 44.63 | 44.96 | 43.61 | 45.47 | 39.65 | 38.85 | 39.34 | 39.66 | 41.81 | 42.24 | 42.02 |
| 240 | 57.59 | 60.37 | 58.47 | 61.18 | 58.22 | 60.10 | 57.81 | 61.11 | 58.02 | 60.69 | 59.36 |
| $k_{obs}$ (M^-1 sec^-1) | 57.71 | 60.08 | 58.81 | 61.07 | 53.77 | 53.21 | 52.40 | 54.46 | 55.67 | 57.20 | 56.44 |
| $k_{adj}$ (M^-1 sec^-1) | 74.58 | 60.08 | 77.58 | 61.07 | 56.76 | 53.21 | 52.40 | 54.46 | 65.33 | 57.20 | 61.27 |
| Unreacted Fracrtion (%) | 8.41 | 0.00 | 8.99 | 0.00 | 1.89 | 0.00 | 0.00 | 0.00 | 4.82 | 0.00 | 2.41 |

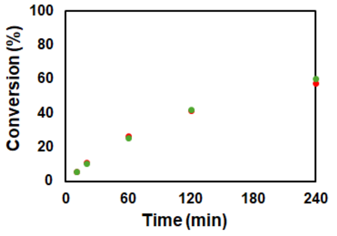

### - LAF Scaffold

#### LAF (50 μM; 1% PUMA), 0.1 μM mCh-ST-Bcl: 804.93 $M^{-1}$ $s^{-1}$

| Client Concentration = 2.618 μM | Conversion (%) | | | | | | | | | | |
|---|---|---|---|---|---|---|---|---|---|---|---|
| GFP client ratio = 1.0 [mCh] | Sample 1 - Gel 1 | | Sample 1 - Gel 2 | | Sample 2 - Gel 1 | | Sample 2 - Gel 2 | | Average | | |
| Reaction Time (min) | mCh-ST | GFP-SC | mCh-ST | GFP-SC | mCh-ST | GFP-SC | mCh-ST | GFP-SC | mCh-ST | GFP-SC | Total |
| 5 | 27.92 | 26.63 | 28.44 | 25.04 | 28.54 | 27.69 | 29.90 | 27.87 | 28.70 | 26.81 | 27.76 |
| 10 | 37.12 | 34.81 | 36.54 | 33.73 | 36.74 | 36.09 | 38.22 | 36.29 | 37.16 | 35.23 | 36.19 |
| 20 | 44.90 | 42.36 | 45.97 | 42.88 | 44.21 | 44.67 | 45.63 | 44.59 | 45.18 | 43.63 | 44.40 |
| 60 | 60.14 | 60.03 | 61.23 | 61.41 | 58.22 | 61.88 | 59.21 | 61.66 | 59.70 | 61.24 | 60.47 |
| 120 | 66.39 | 67.71 | 66.50 | 67.81 | 64.57 | 68.82 | 64.89 | 68.19 | 65.59 | 68.13 | 66.86 |
| 240 | 75.38 | 75.82 | 75.12 | 77.30 | 71.50 | 76.73 | 72.28 | 76.54 | 73.57 | 76.60 | 75.08 |
| $k_{obs}$ (M^-1 sec^-1) | 214.61 | 200.59 | 221.62 | 203.59 | 195.19 | 224.05 | 212.68 | 222.19 | 211.03 | 212.60 | 211.81 |
| $k_{adj}$ (M^-1 sec^-1) | 829.55 | 676.56 | 842.80 | 601.28 | 972.66 | 720.42 | 1054.61 | 741.55 | 924.91 | 684.95 | 804.93 |
| Unreacted Fracrtion (%) | 26.19 | 24.34 | 26.00 | 22.37 | 29.76 | 23.50 | 29.50 | 24.05 | 27.86 | 23.56 | 25.71 |

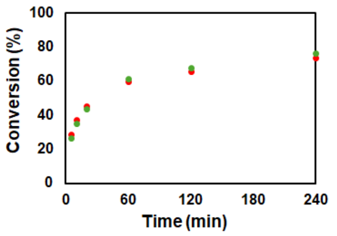

#### LAF (50 µM; 2.5% PUMA), 0.1 µM mCh-ST-Bcl: 846.68 $M^{-1}$ $s^{-1}$

| Client Concentration = 2.500 µM | Conversion (%) | | | | | | | | | | |
|---|---|---|---|---|---|---|---|---|---|---|---|
| GFP client ratio = 0.98 [mCh] | Sample 1 - Gel 1 | | Sample 1 - Gel 2 | | Sample 2 - Gel 1 | | Sample 2 - Gel 2 | | Average | | |
| Reaction Time (min) | mCh-ST | GFP-SC | mCh-ST | GFP-SC | mCh-ST | GFP-SC | mCh-ST | GFP-SC | mCh-ST | GFP-SC | Total |
| 5 | 26.43 | 26.01 | 28.40 | 27.13 | 29.77 | 27.45 | 29.17 | 27.67 | 28.44 | 27.07 | 27.75 |
| 10 | 39.16 | 36.31 | 40.45 | 38.10 | 35.77 | 36.09 | 36.19 | 35.81 | 37.89 | 36.58 | 37.24 |
| 20 | 48.01 | 43.43 | 46.12 | 42.93 | 43.87 | 45.17 | 43.51 | 44.67 | 45.38 | 44.05 | 44.72 |
| 60 | 60.67 | 58.69 | 61.06 | 60.27 | 59.03 | 63.08 | 58.92 | 62.83 | 59.92 | 61.22 | 60.57 |
| 120 | 67.29 | 66.27 | 67.00 | 65.79 | 65.34 | 72.10 | 65.21 | 72.03 | 66.21 | 69.05 | 67.63 |
| 240 | 76.31 | 77.02 | 75.74 | 77.15 | 70.61 | 78.47 | 71.49 | 78.40 | 73.54 | 77.76 | 75.65 |
| $k_{obs}$ (M^-1 sec^-1) | 246.80 | 210.61 | 247.49 | 220.27 | 206.08 | 251.25 | 205.57 | 247.58 | 226.49 | 232.43 | 229.46 |
| $k_{adj}$ (M^-1 sec^-1) | 892.73 | 747.46 | 969.67 | 806.82 | 1024.93 | 674.20 | 989.28 | 668.39 | 969.15 | 724.22 | 846.68 |
| Unreacted Fracrtion (%) | 25.24 | 24.98 | 26.23 | 25.34 | 29.75 | 20.56 | 29.30 | 20.65 | 27.63 | 22.88 | 25.26 |

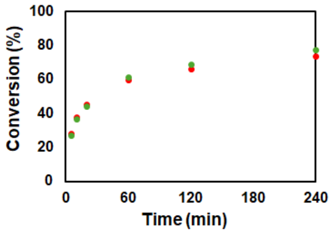

#### LAF (50 µM; 5% PUMA), 0.1 µM mCh-ST-Bcl: 839.65 $M^{-1}$ $s^{-1}$

| Client Concentration = 2.523 µM | Conversion (%) | | | | | | | | | | |
|---|---|---|---|---|---|---|---|---|---|---|---|
| GFP client ratio = 0.96 [mCh] | Sample 1 - Gel 1 | | Sample 1 - Gel 2 | | Sample 2 - Gel 1 | | Sample 2 - Gel 2 | | Average | | |
| Reaction Time (min) | mCh-ST | GFP-SC | mCh-ST | GFP-SC | mCh-ST | GFP-SC | mCh-ST | GFP-SC | mCh-ST | GFP-SC | Total |
| 5 | 30.35 | 26.38 | 28.28 | 25.15 | 28.13 | 28.46 | 28.27 | 26.99 | 28.76 | 26.74 | 27.75 |
| 10 | 37.06 | 34.90 | 37.36 | 34.41 | 35.59 | 36.01 | 35.13 | 35.55 | 36.28 | 35.22 | 35.75 |
| 20 | 46.81 | 45.90 | 46.31 | 45.38 | 45.88 | 46.33 | 44.96 | 45.55 | 45.99 | 45.79 | 45.89 |
| 60 | 59.12 | 56.91 | 57.63 | 56.35 | 59.79 | 63.62 | 58.50 | 62.91 | 58.76 | 59.95 | 59.35 |
| 120 | 66.50 | 69.24 | 68.72 | 70.67 | 65.89 | 71.67 | 65.23 | 71.05 | 66.59 | 70.66 | 68.62 |
| 240 | 73.52 | 76.66 | 73.68 | 75.75 | 69.74 | 78.42 | 70.49 | 77.67 | 71.85 | 77.13 | 74.49 |
| $k_{obs}$ (M^-1 sec^-1) | 228.73 | 214.97 | 223.94 | 209.84 | 210.82 | 256.49 | 201.41 | 243.34 | 216.23 | 231.16 | 223.69 |
| $k_{adj}$ (M^-1 sec^-1) | 1021.41 | 727.56 | 919.07 | 684.08 | 999.77 | 706.45 | 976.53 | 682.35 | 979.19 | 700.11 | 839.65 |
| Unreacted Fracrtion (%) | 28.08 | 24.27 | 27.00 | 23.81 | 29.33 | 20.96 | 29.56 | 21.37 | 28.50 | 22.60 | 25.55 |

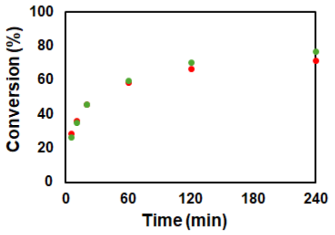

#### LAF (50 µM; 10% PUMA), 0.1 µM mCh-ST-Bcl: 790.45 $M^{-1}$ $s^{-1}$

| Client Concentration = 2.296 µM | Conversion (%) | | | | | | | | | | |
|---|---|---|---|---|---|---|---|---|---|---|---|
| GFP client ratio = 0.94 [mCh] | Sample 1 - Gel 1 | | Sample 1 - Gel 2 | | Sample 2 - Gel 1 | | Sample 2 - Gel 2 | | Average | | |
| Reaction Time (min) | mCh-ST | GFP-SC | mCh-ST | GFP-SC | mCh-ST | GFP-SC | mCh-ST | GFP-SC | mCh-ST | GFP-SC | Total |
| 5 | 25.36 | 24.08 | 26.17 | 25.63 | 26.01 | 25.44 | 25.97 | 25.95 | 25.88 | 25.27 | 25.58 |
| 10 | 36.10 | 33.94 | 36.44 | 34.94 | 33.40 | 33.79 | 33.33 | 34.68 | 34.82 | 34.34 | 34.58 |
| 20 | 44.33 | 42.64 | 43.44 | 41.10 | 41.98 | 43.34 | 43.34 | 45.23 | 43.27 | 43.08 | 43.17 |
| 60 | 57.78 | 59.31 | 60.19 | 59.50 | 58.49 | 62.01 | 57.97 | 62.91 | 58.61 | 60.93 | 59.77 |
| 120 | 65.03 | 69.19 | 67.27 | 69.40 | 62.68 | 69.85 | 63.69 | 71.00 | 64.67 | 69.86 | 67.27 |
| 240 | 73.96 | 78.15 | 74.88 | 78.01 | 69.33 | 77.65 | 71.52 | 77.71 | 72.42 | 77.88 | 75.15 |
| $k_{obs}$ (M^-1 sec^-1) | 219.07 | 227.67 | 234.72 | 228.61 | 193.35 | 242.70 | 202.19 | 260.58 | 212.33 | 239.89 | 226.11 |
| $k_{adj}$ (M^-1 sec^-1) | 909.95 | 640.06 | 848.87 | 664.04 | 970.24 | 667.60 | 917.20 | 705.64 | 911.57 | 669.33 | 790.45 |
| Unreacted Fracrtion (%) | 27.43 | 21.51 | 25.39 | 21.94 | 30.38 | 21.18 | 28.89 | 20.88 | 28.02 | 21.38 | 24.70 |

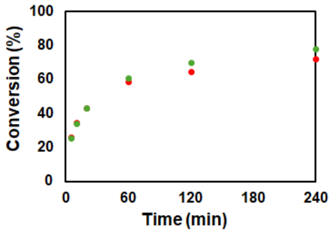

#### LAF (50 µM; 5% PUMA), 0.2 µM mCh-ST-Bcl: 491.74 $M^{-1}$ $s^{-1}$

| Client Concentration = 7.154 µM | Conversion (%) | | | | | | | | | | |
|---|---|---|---|---|---|---|---|---|---|---|---|
| GFP client ratio = 1.02 [mCh] | Sample 1 - Gel 1 | | Sample 1 - Gel 2 | | Sample 2 - Gel 1 | | Sample 2 - Gel 2 | | Average | | |
| Reaction Time (min) | mCh-ST | GFP-SC | mCh-ST | GFP-SC | mCh-ST | GFP-SC | mCh-ST | GFP-SC | mCh-ST | GFP-SC | Total |
| 5 | 45.48 | 44.86 | 43.19 | 39.31 | 34.99 | 31.96 | 34.82 | 33.73 | 39.62 | 37.47 | 38.54 |
| 10 | 53.09 | 49.54 | 53.34 | 51.09 | 43.89 | 41.36 | 43.99 | 41.85 | 48.58 | 45.96 | 47.27 |
| 20 | 58.90 | 56.35 | 60.17 | 57.08 | 53.74 | 52.34 | 54.39 | 53.06 | 56.80 | 54.71 | 55.75 |
| 60 | 71.95 | 72.26 | 71.86 | 72.81 | 66.66 | 68.54 | 68.02 | 69.78 | 69.62 | 70.85 | 70.24 |
| 120 | 77.05 | 78.29 | 77.24 | 78.36 | 72.31 | 75.70 | 73.23 | 76.13 | 74.96 | 77.12 | 76.04 |
| 240 | 76.62 | 78.98 | 76.61 | 79.56 | 76.37 | 80.15 | 75.90 | 80.64 | 76.37 | 79.83 | 78.10 |
| $k_{obs}$ (M^-1 sec^-1) | 205.60 | 187.02 | 203.75 | 181.31 | 128.01 | 124.20 | 132.28 | 131.61 | 167.41 | 156.04 | 161.72 |
| $k_{adj}$ (M^-1 sec^-1) | 718.72 | 572.29 | 667.70 | 484.41 | 435.52 | 306.02 | 429.14 | 320.08 | 562.77 | 420.70 | 491.74 |
| Unreacted Fracrtion (%) | 22.82 | 21.07 | 22.19 | 19.48 | 23.79 | 18.84 | 23.15 | 18.51 | 22.99 | 19.48 | 21.23 |

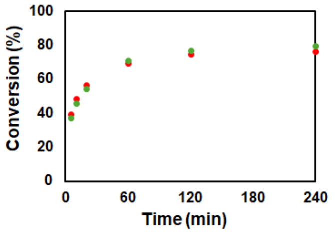

#### LAF (50 µM; 5% PUMA), 0.05 µM mCh-ST-Bcl: 729.69 $M^{-1}$ $s^{-1}$

| Client Concentration = 1.661 µM | Conversion (%) | | | | | | | | | | |
|---|---|---|---|---|---|---|---|---|---|---|---|
| GFP client ratio = 0.95 [mCh] | Sample 1 - Gel 1 | | Sample 1 - Gel 2 | | Sample 2 - Gel 1 | | Sample 2 - Gel 2 | | Average | | |
| Reaction Time (min) | mCh-ST | GFP-SC | mCh-ST | GFP-SC | mCh-ST | GFP-SC | mCh-ST | GFP-SC | mCh-ST | GFP-SC | Total |
| 5 | 21.01 | 18.15 | 24.07 | 20.45 | 21.32 | 18.56 | 20.61 | 19.01 | 21.75 | 19.04 | 20.40 |
| 10 | 22.90 | 22.04 | 23.75 | 22.22 | 26.56 | 24.81 | 26.67 | 25.20 | 24.97 | 23.57 | 24.27 |
| 20 | 32.58 | 30.48 | 31.52 | 29.45 | 32.51 | 29.33 | 30.57 | 30.05 | 31.79 | 29.83 | 30.81 |
| 60 | 50.43 | 48.82 | 48.16 | 48.37 | 49.75 | 49.85 | 51.09 | 50.61 | 49.86 | 49.41 | 49.63 |
| 120 | 53.74 | 56.32 | 55.20 | 56.49 | 57.89 | 61.21 | 58.52 | 61.11 | 56.34 | 58.78 | 57.56 |
| 240 | 68.30 | 65.84 | 66.18 | 68.14 | 66.17 | 71.23 | 65.66 | 71.07 | 66.58 | 69.07 | 67.82 |
| $k_{obs}$ (M^-1 sec^-1) | 154.21 | 145.50 | 149.27 | 148.43 | 163.31 | 169.52 | 163.30 | 173.39 | 157.52 | 159.21 | 158.37 |
| $k_{adj}$ (M^-1 sec^-1) | 785.13 | 670.18 | 909.60 | 629.37 | 903.07 | 533.02 | 836.29 | 570.90 | 858.52 | 600.87 | 729.69 |
| Unreacted Fracrtion (%) | 32.20 | 31.33 | 34.50 | 29.90 | 33.04 | 24.84 | 32.19 | 25.51 | 32.98 | 27.90 | 30.44 |

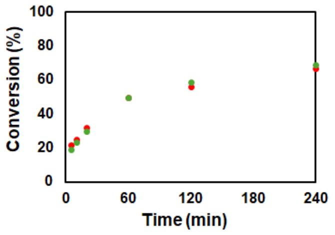

#### LAF (50 µM; 5% PUMA), 0.2 µM mCh-ST-Bcl, 40mM NaCl: 4212.37 M$^{-1}$ s$^{-1}$

| Client Concentration = 1.921 µM | Conversion (%) | | | | | | | | | | |
|---|---|---|---|---|---|---|---|---|---|---|---|
| GFP client ratio = 1.00 [mCh] | Sample 1 - Gel 1 | | Sample 1 - Gel 2 | | Sample 2 - Gel 1 | | Sample 2 - Gel 2 | | Average | | |
| Reaction Time (min) | mCh-ST | GFP-SC | mCh-ST | GFP-SC | mCh-ST | GFP-SC | mCh-ST | GFP-SC | mCh-ST | GFP-SC | Total |
| 5 | 53.28 | 50.11 | 55.45 | 51.25 | 46.76 | 45.22 | 46.51 | 45.73 | 50.50 | 48.08 | 49.29 |
| 10 | 58.59 | 54.79 | 57.57 | 54.16 | 53.46 | 51.90 | 54.21 | 52.53 | 55.96 | 53.34 | 54.65 |
| 20 | 66.20 | 62.21 | 64.27 | 61.42 | 57.43 | 54.95 | 57.56 | 55.09 | 61.37 | 58.42 | 59.89 |
| 60 | 73.53 | 69.60 | 72.76 | 69.11 | 63.34 | 62.88 | 62.51 | 63.83 | 68.03 | 66.35 | 67.19 |
| 120 | 76.03 | 75.38 | 77.06 | 74.80 | 68.31 | 69.52 | 67.69 | 70.39 | 72.27 | 72.52 | 72.40 |
| 240 | 80.09 | 77.52 | 79.43 | 77.64 | 71.15 | 70.15 | 71.52 | 70.40 | 75.54 | 73.93 | 74.74 |
| $k_{obs}$ (M^-1 sec^-1) | 1092.77 | 870.71 | 1079.14 | 859.61 | 648.99 | 584.58 | 649.09 | 610.25 | 867.50 | 731.29 | 799.39 |
| $k_{adj}$ (M^-1 sec^-1) | 4072.08 | 3649.55 | 4407.41 | 3842.45 | 4683.43 | 4115.76 | 4825.24 | 4103.06 | 4497.04 | 3927.71 | 4212.37 |
| Unreacted Fracrtion (%) | 22.14 | 24.42 | 22.95 | 25.05 | 31.28 | 31.39 | 31.55 | 30.79 | 26.98 | 27.91 | 27.45 |

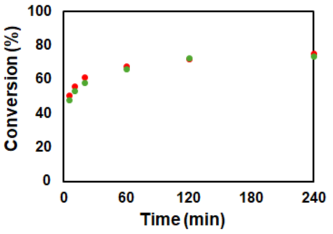

#### LAF (50 µM; 5% PUMA), 0.1 µM mCh-ST-Bcl, 3% PEG8000 induced: 156.30 M$^{-1}$ s$^{-1}$

| Client Concentration = 5.374 µM | Conversion (%) | | | | | | | | | | |
|---|---|---|---|---|---|---|---|---|---|---|---|
| GFP client ratio = 1.1 [mCh] | Sample 1 - Gel 1 | | Sample 1 - Gel 2 | | Sample 2 - Gel 1 | | Sample 2 - Gel 2 | | Average | | |
| Reaction Time (min) | mCh-ST | GFP-SC | mCh-ST | GFP-SC | mCh-ST | GFP-SC | mCh-ST | GFP-SC | mCh-ST | GFP-SC | Total |
| 5 | 15.12 | 15.96 | 16.16 | 14.13 | 14.38 | 12.89 | 13.42 | 13.05 | 14.77 | 14.01 | 14.39 |
| 10 | 24.70 | 23.63 | 23.77 | 22.57 | 19.33 | 20.63 | 18.83 | 20.59 | 21.66 | 21.86 | 21.76 |
| 20 | 35.92 | 33.55 | 33.44 | 31.73 | 28.69 | 30.19 | 29.05 | 30.31 | 31.78 | 31.45 | 31.61 |
| 60 | 54.67 | 52.42 | 52.88 | 51.69 | 46.76 | 52.42 | 47.73 | 52.31 | 50.51 | 52.21 | 51.36 |
| 120 | 64.45 | 64.35 | 62.56 | 62.75 | 56.92 | 65.86 | 57.97 | 65.62 | 60.47 | 64.64 | 62.56 |
| 240 | 69.13 | 70.50 | 67.71 | 69.73 | 63.11 | 75.20 | 65.08 | 75.77 | 66.26 | 72.80 | 69.53 |
| $k_{obs}$ (M^-1 sec^-1) | 61.74 | 58.20 | 55.76 | 53.78 | 40.41 | 57.26 | 42.48 | 57.37 | 50.10 | 56.65 | 53.37 |
| $k_{adj}$ (M^-1 sec^-1) | 195.80 | 170.04 | 194.81 | 155.38 | 171.45 | 105.92 | 152.54 | 104.45 | 178.65 | 133.95 | 156.30 |
| Unreacted Fracrtion (%) | 24.92 | 23.58 | 26.69 | 23.68 | 30.90 | 15.06 | 28.16 | 14.70 | 27.67 | 19.25 | 23.46 |

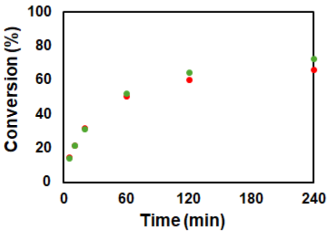

#### LAF (50 µM; 5% PUMA), 0.2 µM mCh-ST-Bcl, 3% PEG8000 induced: 133.76 M$^{-1}$ s$^{-1}$

| Client Concentration = 11.51 µM | Conversion (%) | | | | | | | | | | |
|---|---|---|---|---|---|---|---|---|---|---|---|
| GFP client ratio = 1.18 [mCh] | Sample 1 - Gel 1 | | Sample 1 - Gel 2 | | Sample 2 - Gel 1 | | Sample 2 - Gel 2 | | Average | | |
| Reaction Time (min) | mCh-ST | GFP-SC | mCh-ST | GFP-SC | mCh-ST | GFP-SC | mCh-ST | GFP-SC | mCh-ST | GFP-SC | Total |
| 5 | 20.94 | 21.63 | 20.83 | 20.70 | 22.09 | 20.34 | 23.48 | 20.46 | 21.83 | 20.78 | 21.31 |
| 10 | 31.50 | 33.43 | 32.04 | 33.64 | 31.03 | 30.68 | 30.82 | 30.20 | 31.35 | 31.99 | 31.67 |
| 20 | 43.38 | 46.77 | 42.86 | 46.62 | 43.04 | 42.77 | 43.65 | 42.73 | 43.23 | 44.72 | 43.98 |
| 60 | 58.79 | 64.97 | 57.66 | 63.78 | 60.84 | 62.28 | 61.49 | 62.51 | 59.69 | 63.38 | 61.54 |
| 120 | 63.85 | 73.92 | 63.66 | 73.54 | 68.90 | 72.33 | 68.96 | 72.53 | 66.34 | 73.08 | 69.71 |
| 240 | 66.44 | 75.93 | 65.31 | 75.58 | 71.77 | 78.38 | 72.40 | 77.65 | 68.98 | 76.89 | 72.93 |
| $k_{obs}$ (M^-1 sec^-1) | 36.99 | 52.97 | 35.72 | 51.45 | 42.40 | 46.03 | 43.81 | 45.82 | 39.73 | 49.07 | 44.40 |
| $k_{adj}$ (M^-1 sec^-1) | 171.08 | 124.71 | 178.87 | 124.57 | 134.44 | 99.21 | 137.46 | 99.75 | 155.46 | 112.06 | 133.76 |
| Unreacted Fracrtion (%) | 29.90 | 18.83 | 30.99 | 19.35 | 24.03 | 17.20 | 23.78 | 17.43 | 27.17 | 18.20 | 22.69 |

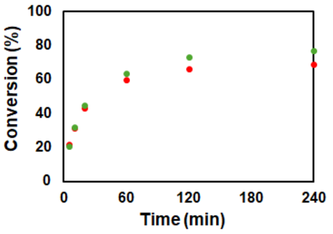

#### LAF (50 µM; 5% PUMA), 0.1 µM mCh-ST-Bcl, 5% PEG8000 induced: 89.07 M$^{-1}$ s$^{-1}$

| Client Concentration = 9.764 µM | Conversion (%) | | | | | | | | | | |
|---|---|---|---|---|---|---|---|---|---|---|---|
| GFP client ratio = 0.95 [mCh] | Sample 1 - Gel 1 | | Sample 1 - Gel 2 | | Sample 2 - Gel 1 | | Sample 2 - Gel 2 | | Average | | |
| Reaction Time (min) | mCh-ST | GFP-SC | mCh-ST | GFP-SC | mCh-ST | GFP-SC | mCh-ST | GFP-SC | mCh-ST | GFP-SC | Total |
| 5 | 24.73 | 18.56 | 24.05 | 18.82 | 17.49 | 15.04 | 16.50 | 14.35 | 20.69 | 16.69 | 18.69 |
| 10 | 34.35 | 27.20 | 33.84 | 27.65 | 36.21 | 30.91 | 35.73 | 30.24 | 35.03 | 29.00 | 32.02 |
| 20 | 45.54 | 37.86 | 44.73 | 38.29 | 43.93 | 38.65 | 41.90 | 36.79 | 44.02 | 37.90 | 40.96 |
| 60 | 70.14 | 63.44 | 69.51 | 63.32 | 66.10 | 62.04 | 64.64 | 61.08 | 67.60 | 62.47 | 65.03 |
| 120 | 80.04 | 75.45 | 80.09 | 75.84 | 76.55 | 74.86 | 76.41 | 74.16 | 78.27 | 75.08 | 76.68 |
| 240 | 86.14 | 84.77 | 86.87 | 85.84 | 83.80 | 83.47 | 82.77 | 82.82 | 84.90 | 84.23 | 84.56 |
| $k_{obs}$ (M^-1 sec^-1) | 75.33 | 52.50 | 73.32 | 53.58 | 64.96 | 52.18 | 60.74 | 49.16 | 68.59 | 51.86 | 60.22 |
| $k_{adj}$ (M^-1 sec^-1) | 109.19 | 71.48 | 102.28 | 70.87 | 106.04 | 78.35 | 100.61 | 73.71 | 104.53 | 73.60 | 89.07 |
| Unreacted Fracrtion (%) | 8.85 | 7.65 | 8.01 | 6.94 | 11.54 | 9.86 | 11.88 | 9.88 | 10.07 | 8.58 | 9.33 |

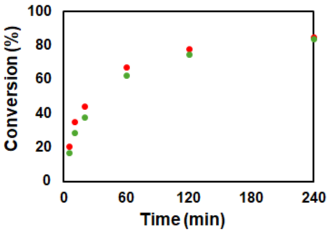

#### LAF (50 µM; 5% PUMA), 0.1 µM mCh-ST-Bcl, 100 µM $NiCl_2$: 287.89 M$^{-1}$ s$^{-1}$

| Client Concentration = 5.543 µM | Conversion (%) | | | | | | | | | | |
|---|---|---|---|---|---|---|---|---|---|---|---|
| GFP client ratio = 0.96 [mCh] | Sample 1 - Gel 1 | | Sample 1 - Gel 2 | | Sample 2 - Gel 1 | | Sample 2 - Gel 2 | | Average | | |
| Reaction Time (min) | mCh-ST | GFP-SC | mCh-ST | GFP-SC | mCh-ST | GFP-SC | mCh-ST | GFP-SC | mCh-ST | GFP-SC | Total |
| 5 | 29.26 | 27.68 | 28.96 | 27.37 | 31.76 | 25.42 | 30.31 | 26.54 | 30.07 | 26.75 | 28.41 |
| 10 | 38.89 | 37.14 | 38.77 | 37.30 | 40.10 | 35.07 | 44.03 | 35.40 | 40.45 | 36.23 | 38.34 |
| 20 | 49.55 | 48.81 | 50.24 | 49.36 | 52.60 | 46.04 | 52.18 | 46.13 | 51.14 | 47.58 | 49.36 |
| 60 | 64.96 | 68.95 | 67.12 | 70.12 | 73.85 | 66.02 | 75.53 | 67.00 | 70.36 | 68.02 | 69.19 |
| 120 | 73.98 | 79.92 | 74.75 | 80.42 | 78.90 | 75.11 | 83.84 | 75.99 | 77.87 | 77.86 | 77.86 |
| 240 | 78.83 | 86.37 | 79.23 | 85.72 | 83.94 | 80.94 | 85.35 | 80.69 | 81.84 | 83.43 | 82.63 |
| $k_{obs}$ (M^-1 sec^-1) | 134.61 | 146.03 | 140.52 | 149.20 | 173.81 | 120.83 | 189.56 | 125.04 | 159.62 | 135.28 | 147.45 |
| $k_{adj}$ (M^-1 sec^-1) | 357.08 | 236.38 | 344.30 | 239.51 | 318.99 | 250.65 | 298.71 | 257.53 | 329.77 | 246.02 | 287.89 |
| Unreacted Fracrtion (%) | 20.25 | 11.05 | 18.97 | 10.92 | 13.52 | 16.13 | 10.42 | 15.97 | 15.79 | 13.52 | 14.65 |

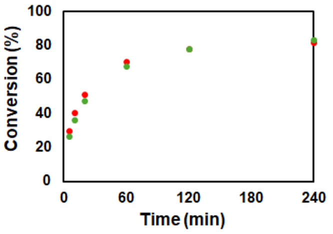

##### LAF (100 μM; 5% PUMA), 0.1 μM mCh-ST-Bcl, 100 μM $NiCl_2$: 1060.12 $M^{-1}$ $s^{-1}$

| Client Concentration = 2.029 μM | Conversion (%) | | | | | | | | | | |
|---|---|---|---|---|---|---|---|---|---|---|---|
| GFP client ratio = 0.9 [mCh] | Sample 1 - Gel 1 | | Sample 1 - Gel 2 | | Sample 2 - Gel 1 | | Sample 2 - Gel 2 | | Average | | |
| Reaction Time (min) | mCh-ST | GFP-SC | mCh-ST | GFP-SC | mCh-ST | GFP-SC | mCh-ST | GFP-SC | mCh-ST | GFP-SC | Total |
| 5 | 27.92 | 22.44 | 29.30 | 22.48 | 27.03 | 25.39 | 30.71 | 25.37 | 28.74 | 23.92 | 26.33 |
| 10 | 31.35 | 25.16 | 31.27 | 24.89 | 34.28 | 32.04 | 35.64 | 33.11 | 33.14 | 28.80 | 30.97 |
| 20 | 49.82 | 43.06 | 46.11 | 41.21 | 49.17 | 49.47 | 44.66 | 45.73 | 47.44 | 44.87 | 46.16 |
| 60 | 53.42 | 48.35 | 52.48 | 47.11 | 60.31 | 66.30 | 59.71 | 66.26 | 56.48 | 57.00 | 56.74 |
| 120 | 61.57 | 59.21 | 63.05 | 58.88 | 61.99 | 71.03 | 63.74 | 71.86 | 62.59 | 65.25 | 63.92 |
| 240 | 68.16 | 65.20 | 68.87 | 65.58 | 73.41 | 81.78 | 73.27 | 81.86 | 70.93 | 73.60 | 72.27 |
| $k_{obs}$ (M^-1 sec^-1) | 220.53 | 150.84 | 210.69 | 143.22 | 267.38 | 325.39 | 262.45 | 313.65 | 240.26 | 233.27 | 236.77 |
| $k_{adj}$ (M^-1 sec^-1) | 1410.23 | 994.91 | 1315.91 | 941.37 | 1192.40 | 708.15 | 1255.10 | 662.89 | 1293.41 | 826.83 | 1060.12 |
| Unreacted Fracrtion (%) | 33.22 | 34.73 | 32.79 | 34.79 | 28.52 | 17.12 | 29.07 | 16.52 | 30.90 | 25.79 | 28.34 |

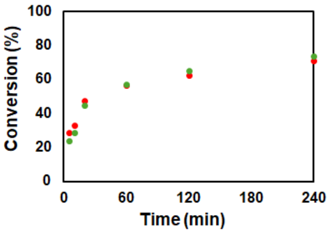

#### - MBP–LAF Scaffold

##### 50% MBP–LAF + 50% LAF (total 50 μM; 5% PUMA), 0.1 μM mCh-ST-Bcl: 497.38 $M^{-1}$ $s^{-1}$

| Client Concentration = 1.836 μM | Conversion (%) | | | | | | | | | | |
|---|---|---|---|---|---|---|---|---|---|---|---|
| GFP client ratio = 1.4 [mCh] | Sample 1 - Gel 1 | | Sample 1 - Gel 2 | | Sample 2 - Gel 1 | | Sample 2 - Gel 2 | | Average | | |
| Reaction Time (min) | mCh-ST | GFP-SC | mCh-ST | GFP-SC | mCh-ST | GFP-SC | mCh-ST | GFP-SC | mCh-ST | GFP-SC | Total |
| 5 | 14.79 | 12.89 | 14.07 | 12.57 | 10.00 | 9.75 | 9.55 | 9.23 | 12.10 | 11.11 | 11.61 |
| 10 | 20.51 | 18.88 | 21.11 | 18.91 | 24.30 | 22.99 | 23.22 | 23.02 | 22.28 | 20.95 | 21.62 |
| 20 | 30.98 | 28.93 | 30.74 | 28.25 | 27.62 | 27.21 | 32.08 | 32.79 | 30.36 | 29.29 | 29.83 |
| 60 | 50.53 | 49.75 | 51.35 | 49.96 | 43.64 | 41.48 | 41.84 | 40.93 | 46.84 | 45.53 | 46.19 |
| 120 | 59.81 | 60.88 | 59.09 | 58.42 | 55.97 | 56.34 | 55.90 | 59.37 | 57.69 | 58.75 | 58.22 |
| 240 | 67.41 | 66.24 | 64.31 | 65.21 | 61.50 | 64.77 | 60.60 | 63.41 | 63.46 | 64.90 | 64.18 |
| $k_{obs}$ (M^-1 sec^-1) | 141.99 | 134.95 | 137.58 | 128.23 | 109.52 | 109.40 | 133.08 | 112.12 | 130.54 | 121.18 | 125.86 |
| $k_{adj}$ (M^-1 sec^-1) | 476.74 | 409.85 | 541.01 | 430.72 | 544.75 | 423.02 | 627.77 | 525.16 | 547.57 | 447.19 | 497.38 |
| Unreacted Fracrtion (%) | 26.53 | 25.20 | 29.25 | 27.08 | 33.44 | 29.66 | 35.34 | 31.42 | 31.14 | 28.34 | 29.74 |

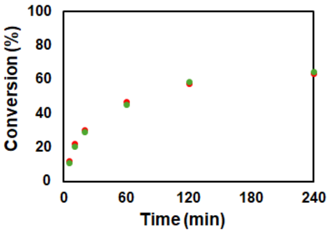

##### MBP–LAF (50 μM; 5% PUMA), 0.5 μM mCh-ST-Bcl: 199.13 $M^{-1}$ $s^{-1}$

| Client Concentration = 5.415 μM | Conversion (%) | | | | | | | | | | |
|---|---|---|---|---|---|---|---|---|---|---|---|
| GFP client ratio = 1.5 [mCh] | Sample 1 - Gel 1 | | Sample 1 - Gel 2 | | Sample 2 - Gel 1 | | Sample 2 - Gel 2 | | Average | | |
| Reaction Time (min) | mCh-ST | GFP-SC | mCh-ST | GFP-SC | mCh-ST | GFP-SC | mCh-ST | GFP-SC | mCh-ST | GFP-SC | Total |
| 5 | 13.77 | 11.81 | 15.06 | 12.99 | 13.08 | 11.09 | 14.22 | 11.93 | 14.03 | 11.96 | 12.99 |
| 10 | 21.32 | 18.23 | 22.62 | 18.76 | 20.12 | 15.82 | 20.20 | 15.94 | 21.07 | 17.19 | 19.13 |
| 20 | 29.88 | 29.13 | 31.75 | 29.32 | 28.94 | 26.97 | 28.78 | 26.50 | 29.84 | 27.98 | 28.91 |
| 60 | 45.66 | 44.15 | 45.34 | 43.91 | 51.40 | 43.46 | 50.56 | 43.06 | 48.24 | 43.65 | 45.94 |
| 120 | 54.79 | 53.43 | 55.17 | 53.90 | 55.30 | 52.04 | 55.26 | 51.46 | 55.13 | 52.71 | 53.92 |
| 240 | 61.12 | 59.77 | 61.01 | 60.05 | 58.22 | 58.32 | 59.06 | 57.57 | 59.85 | 58.93 | 59.39 |
| $k_{obs}$ (M^-1 sec^-1) | 38.12 | 34.67 | 39.38 | 35.23 | 32.76 | 31.79 | 39.83 | 30.96 | 37.52 | 33.16 | 35.34 |
| $k_{adj}$ (M^-1 sec^-1) | 210.92 | 185.50 | 244.97 | 192.53 | 217.74 | 151.47 | 216.57 | 173.37 | 222.55 | 175.72 | 199.13 |
| Unreacted Fracrtion (%) | 34.74 | 34.93 | 35.95 | 35.07 | 34.77 | 34.29 | 34.61 | 36.29 | 35.02 | 35.14 | 35.08 |

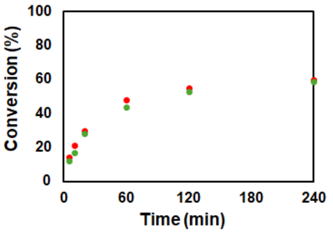

#### - FUS Scaffold

##### FUS (50 μM; 5% PUMA), 0.5 μM mCh-ST-Bcl: 180.50 $M^{-1}$ $s^{-1}$

| Client Concentration = 8.887 μM | Conversion (%) | | | | | | | | | | |
|---|---|---|---|---|---|---|---|---|---|---|---|
| GFP client ratio = 1.75 [mCh] | Sample 1 - Gel 1 | | Sample 1 - Gel 2 | | Sample 2 - Gel 1 | | Sample 2 - Gel 2 | | Average | | |
| Reaction Time (min) | mCh-ST | GFP-SC | mCh-ST | GFP-SC | mCh-ST | GFP-SC | mCh-ST | GFP-SC | mCh-ST | GFP-SC | Total |
| 5 | 9.57 | 10.30 | 9.60 | 11.35 | 9.87 | 13.21 | 8.53 | 11.58 | 9.40 | 11.61 | 10.50 |
| 10 | 15.39 | 15.05 | 15.31 | 16.33 | 15.95 | 17.59 | 14.62 | 15.93 | 15.32 | 16.23 | 15.77 |
| 20 | 22.71 | 21.75 | 25.32 | 26.48 | 22.86 | 24.36 | 21.10 | 21.22 | 23.00 | 23.46 | 23.23 |
| 60 | 34.08 | 29.76 | 33.24 | 33.35 | 30.98 | 32.51 | 30.98 | 30.25 | 32.32 | 31.47 | 31.90 |
| 120 | 46.12 | 38.31 | 47.14 | 41.62 | 42.33 | 44.61 | 41.17 | 41.81 | 44.19 | 41.59 | 42.89 |
| 240 | 48.43 | 42.43 | 49.10 | 44.89 | 43.35 | 44.75 | 42.05 | 43.08 | 45.73 | 43.79 | 44.76 |
| $k_{obs}$ (M^-1 sec^-1) | 12.77 | 9.43 | 13.22 | 11.40 | 10.56 | 11.77 | 9.92 | 10.23 | 11.61 | 10.71 | 11.16 |
| $k_{adj}$ (M^-1 sec^-1) | 126.21 | 200.15 | 133.49 | 225.68 | 185.73 | 218.33 | 166.66 | 187.77 | 153.02 | 207.98 | 180.50 |
| Unreacted Fracrtion (%) | 45.92 | 55.49 | 45.80 | 52.96 | 52.93 | 52.21 | 53.23 | 53.58 | 49.47 | 53.56 | 51.51 |

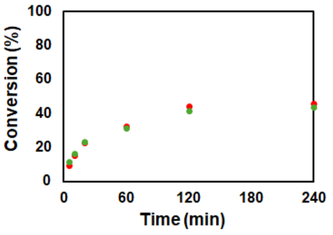

#### - PRM-SH3$_{short}$ + IDPs

##### 10% LAF + 90% PRM-SH3$_{short}$ (total 100 µM; 5% PUMA), 0.1 µM mCh-ST-Bcl: 187.61 $M^{-1}$ $s^{-1}$

| Client Concentration = 4.756 µM | Conversion (%) | | | | | | | | | | |
|---|---|---|---|---|---|---|---|---|---|---|---|
| GFP client ratio = 0.66 [mCh] | Sample 1 - Gel 1 | | Sample 1 - Gel 2 | | Sample 2 - Gel 1 | | Sample 2 - Gel 2 | | Average | | |
| Reaction Time (min) | mCh-ST | GFP-SC | mCh-ST | GFP-SC | mCh-ST | GFP-SC | mCh-ST | GFP-SC | mCh-ST | GFP-SC | Total |
| 5 | 21.06 | 19.42 | 22.07 | 19.16 | 19.20 | 17.49 | 19.17 | 17.47 | 20.38 | 18.39 | 19.38 |
| 10 | 30.05 | 27.04 | 28.69 | 26.79 | 29.38 | 26.35 | 29.26 | 26.06 | 29.34 | 26.56 | 27.95 |
| 20 | 42.00 | 38.97 | 41.19 | 38.81 | 40.95 | 38.03 | 40.53 | 38.09 | 41.17 | 38.47 | 39.82 |
| 60 | 63.67 | 63.06 | 63.06 | 62.32 | 62.59 | 60.41 | 62.29 | 60.97 | 62.90 | 61.69 | 62.30 |
| 120 | 71.15 | 73.17 | 71.24 | 73.12 | 69.48 | 72.55 | 70.35 | 72.83 | 70.55 | 72.92 | 71.74 |
| 240 | 81.74 | 84.53 | 80.13 | 83.22 | 79.41 | 83.02 | 78.38 | 83.00 | 79.92 | 83.44 | 81.68 |
| $k_{obs}$ (M^-1 sec^-1) | 113.09 | 106.79 | 108.98 | 104.19 | 104.11 | 98.32 | 103.16 | 99.09 | 107.34 | 102.10 | 104.72 |
| $k_{adj}$ (M^-1 sec^-1) | 218.31 | 159.14 | 219.97 | 162.28 | 219.23 | 151.79 | 219.33 | 150.82 | 219.21 | 156.01 | 187.61 |
| Unreacted Fracrtion (%) | 15.00 | 9.67 | 15.91 | 10.67 | 16.83 | 10.52 | 17.05 | 10.22 | 16.20 | 10.27 | 13.23 |

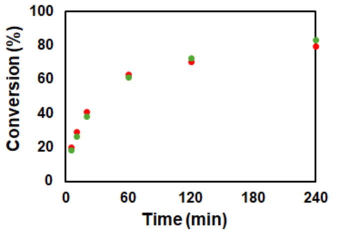

##### 25% LAF + 75% PRM-SH3$_{short}$ (total 100 µM; 5% PUMA), 0.1 µM mCh-ST-Bcl: 303.93 $M^{-1}$ $s^{-1}$

| Client Concentration = 3.106 µM | Conversion (%) | | | | | | | | | | |
|---|---|---|---|---|---|---|---|---|---|---|---|
| GFP client ratio = 0.7 [mCh] | Sample 1 - Gel 1 | | Sample 1 - Gel 2 | | Sample 2 - Gel 1 | | Sample 2 - Gel 2 | | Average | | |
| Reaction Time (min) | mCh-ST | GFP-SC | mCh-ST | GFP-SC | mCh-ST | GFP-SC | mCh-ST | GFP-SC | mCh-ST | GFP-SC | Total |
| 5 | 22.10 | 20.86 | 20.86 | 19.08 | 17.56 | 17.49 | 17.69 | 17.16 | 19.55 | 18.65 | 19.10 |
| 10 | 29.16 | 27.24 | 27.24 | 26.31 | 26.30 | 24.11 | 26.03 | 24.04 | 27.18 | 25.42 | 26.30 |
| 20 | 39.39 | 37.79 | 37.79 | 36.95 | 34.93 | 33.10 | 35.58 | 33.04 | 36.92 | 35.22 | 36.07 |
| 60 | 59.61 | 59.01 | 59.01 | 57.33 | 53.75 | 53.54 | 53.71 | 53.23 | 56.52 | 55.78 | 56.15 |
| 120 | 66.83 | 69.69 | 69.69 | 68.71 | 65.59 | 67.63 | 63.96 | 66.49 | 66.52 | 68.13 | 67.32 |
| 240 | 74.71 | 79.13 | 79.13 | 78.33 | 72.71 | 77.29 | 70.98 | 76.19 | 74.38 | 77.74 | 76.06 |
| $k_{obs}$ (M^-1 sec^-1) | 142.48 | 143.24 | 143.24 | 133.14 | 111.91 | 113.94 | 108.55 | 110.52 | 126.55 | 125.21 | 125.88 |
| $k_{adj}$ (M^-1 sec^-1) | 412.25 | 299.18 | 299.18 | 281.47 | 318.60 | 228.90 | 352.88 | 238.96 | 345.73 | 262.13 | 303.93 |
| Unreacted Fracrtion (%) | 22.51 | 16.67 | 16.67 | 17.03 | 22.74 | 16.28 | 25.03 | 17.79 | 21.74 | 16.94 | 19.34 |

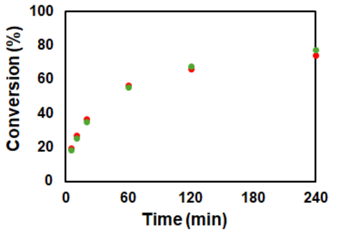

##### 50% LAF + 50% PRM-SH3$_{short}$ (total 100 µM; 5% PUMA), 0.1 µM mCh-ST-Bcl: 761.43 $M^{-1}$ $s^{-1}$

| Client Concentration = 1.558 µM | Conversion (%) | | | | | | | | | | |
|---|---|---|---|---|---|---|---|---|---|---|---|
| GFP client ratio = 0.76 [mCh] | Sample 1 - Gel 1 | | Sample 1 - Gel 2 | | Sample 2 - Gel 1 | | Sample 2 - Gel 2 | | Average | | |
| Reaction Time (min) | mCh-ST | GFP-SC | mCh-ST | GFP-SC | mCh-ST | GFP-SC | mCh-ST | GFP-SC | mCh-ST | GFP-SC | Total |
| 5 | 23.36 | 19.38 | 22.95 | 20.45 | 18.51 | 17.56 | 17.67 | 17.55 | 20.62 | 18.74 | 19.68 |
| 10 | 25.59 | 24.48 | 27.30 | 26.02 | 25.08 | 24.77 | 23.25 | 23.71 | 25.31 | 24.74 | 25.03 |
| 20 | 33.48 | 31.75 | 33.59 | 32.65 | 33.70 | 32.36 | 32.65 | 31.53 | 33.35 | 32.07 | 32.71 |
| 60 | 51.43 | 50.16 | 50.87 | 51.07 | 51.48 | 53.22 | 51.45 | 52.84 | 51.31 | 51.82 | 51.57 |
| 120 | 57.77 | 59.42 | 56.85 | 60.62 | 60.18 | 66.44 | 59.70 | 65.56 | 58.63 | 63.01 | 60.82 |
| 240 | 67.27 | 70.41 | 65.74 | 70.40 | 66.61 | 73.91 | 65.85 | 74.96 | 66.37 | 72.42 | 69.39 |
| $k_{obs}$ (M^-1 sec^-1) | 183.61 | 181.05 | 178.86 | 192.31 | 184.38 | 215.89 | 175.86 | 210.13 | 180.68 | 199.84 | 190.26 |
| $k_{adj}$ (M^-1 sec^-1) | 982.70 | 673.41 | 1123.88 | 744.86 | 816.34 | 523.11 | 749.86 | 477.31 | 918.20 | 604.67 | 761.43 |
| Unreacted Fracrtion (%) | 32.40 | 27.44 | 34.45 | 27.81 | 30.14 | 20.00 | 29.89 | 18.84 | 31.72 | 23.52 | 27.62 |

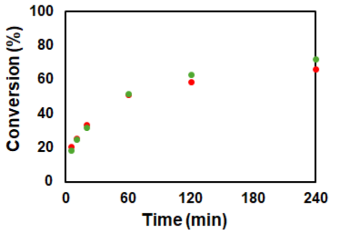

##### 10% TAF + 90% PRM-SH3$_{short}$ (total 100 µM; 5% PUMA), 0.1 µM mCh-ST-Bcl: 471.33 $M^{-1}$ $s^{-1}$

| Client Concentration = 4.593 µM | Conversion (%) | | | | | | | | | | |
|---|---|---|---|---|---|---|---|---|---|---|---|
| GFP client ratio = 0.8 [mCh] | Sample 1 - Gel 1 | | Sample 1 - Gel 2 | | Sample 2 - Gel 1 | | Sample 2 - Gel 2 | | Average | | |
| Reaction Time (min) | mCh-ST | GFP-SC | mCh-ST | GFP-SC | mCh-ST | GFP-SC | mCh-ST | GFP-SC | mCh-ST | GFP-SC | Total |
| 5 | 38.25 | 37.85 | 39.02 | 37.26 | 38.55 | 34.37 | 39.44 | 34.08 | 38.82 | 35.89 | 37.35 |
| 10 | 44.65 | 44.07 | 44.38 | 43.86 | 45.05 | 39.52 | 44.61 | 39.57 | 44.67 | 41.75 | 43.21 |
| 20 | 52.93 | 53.34 | 53.47 | 53.45 | 57.08 | 51.88 | 57.85 | 52.04 | 55.33 | 52.68 | 54.00 |
| 60 | 69.98 | 72.90 | 70.34 | 73.17 | 73.55 | 69.76 | 74.68 | 69.55 | 72.14 | 71.34 | 71.74 |
| 120 | 77.87 | 83.03 | 77.32 | 83.10 | 81.50 | 78.51 | 81.69 | 78.89 | 79.59 | 80.88 | 80.24 |
| 240 | 83.12 | 89.23 | 83.63 | 89.20 | 87.55 | 84.21 | 87.08 | 84.39 | 85.34 | 86.76 | 86.05 |
| $k_{obs}$ (M^-1 sec^-1) | 230.50 | 249.04 | 234.25 | 247.67 | 267.43 | 202.04 | 274.47 | 202.23 | 251.66 | 225.24 | 238.45 |
| $k_{adj}$ (M^-1 sec^-1) | 558.27 | 419.11 | 569.16 | 409.86 | 493.27 | 410.82 | 504.92 | 405.20 | 531.40 | 411.25 | 471.33 |
| Unreacted Fracrtion (%) | 17.97 | 11.33 | 17.98 | 11.04 | 13.11 | 15.25 | 13.05 | 14.98 | 15.53 | 13.15 | 14.34 |

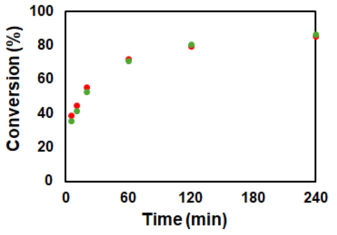

##### 25% TAF + 75% PRM-SH3$_{short}$ (total 100 µM; 5% PUMA), 0.1 µM mCh-ST-Bcl: 871.02 $M^{-1}$ $s^{-1}$

| Client Concentration = 2.147 µM | Conversion (%) | | | | | | | | | | |
|---|---|---|---|---|---|---|---|---|---|---|---|
| GFP client ratio = 1.05 [mCh] | Sample 1 - Gel 1 | | Sample 1 - Gel 2 | | Sample 2 - Gel 1 | | Sample 2 - Gel 2 | | Average | | |
| Reaction Time (min) | mCh-ST | GFP-SC | mCh-ST | GFP-SC | mCh-ST | GFP-SC | mCh-ST | GFP-SC | mCh-ST | GFP-SC | Total |
| 5 | 30.30 | 30.55 | 28.15 | 31.41 | 34.28 | 28.84 | 34.13 | 28.16 | 31.72 | 29.74 | 30.73 |
| 10 | 37.62 | 37.47 | 37.15 | 39.07 | 40.81 | 34.17 | 41.66 | 34.49 | 39.31 | 36.30 | 37.81 |
| 20 | 45.75 | 46.47 | 45.87 | 47.41 | 52.93 | 45.20 | 53.96 | 45.88 | 49.63 | 46.24 | 47.93 |
| 60 | 59.75 | 63.05 | 61.22 | 64.23 | 70.59 | 62.16 | 70.36 | 61.88 | 65.48 | 62.83 | 64.15 |
| 120 | 68.47 | 74.87 | 69.34 | 75.71 | 78.24 | 70.88 | 78.58 | 70.29 | 73.66 | 72.94 | 73.30 |
| 240 | 75.85 | 81.34 | 74.98 | 82.41 | 87.85 | 80.40 | 87.33 | 78.98 | 81.50 | 80.78 | 81.14 |
| $k_{obs}$ (M^-1 sec^-1) | 278.94 | 326.62 | 279.54 | 351.97 | 456.91 | 284.96 | 468.23 | 282.29 | 370.91 | 311.46 | 341.18 |
| $k_{adj}$ (M^-1 sec^-1) | 1068.45 | 805.17 | 983.03 | 838.22 | 844.94 | 750.23 | 883.23 | 794.88 | 944.91 | 797.12 | 871.02 |
| Unreacted Fracrtion (%) | 25.83 | 18.80 | 24.84 | 18.11 | 13.38 | 20.14 | 13.77 | 21.33 | 19.46 | 19.60 | 19.53 |

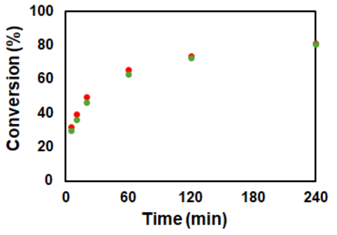

#### 50% TAF + 50% PRM-SH3$_{short}$ (total 100 µM; 5% PUMA), 0.1 µM mCh-ST-Bcl: 783.40 M$^{-1}$ s$^{-1}$

| Client Concentration = 1.099 µM | Conversion (%) | | | | | | | | | | |
|---|---|---|---|---|---|---|---|---|---|---|---|
| GFP client ratio = 1.4 [mCh] | Sample 1 - Gel 1 | | Sample 1 - Gel 2 | | Sample 2 - Gel 1 | | Sample 2 - Gel 2 | | Average | | |
| Reaction Time (min) | mCh-ST | GFP-SC | mCh-ST | GFP-SC | mCh-ST | GFP-SC | mCh-ST | GFP-SC | mCh-ST | GFP-SC | Total |
| 5 | 16.99 | 17.54 | 16.55 | 16.75 | 18.11 | 15.35 | 16.46 | 15.62 | 17.03 | 16.31 | 16.67 |
| 10 | 25.81 | 25.71 | 23.69 | 23.45 | 24.78 | 22.12 | 23.66 | 22.16 | 24.48 | 23.36 | 23.92 |
| 20 | 32.34 | 34.13 | 33.11 | 34.06 | 35.28 | 32.76 | 34.11 | 31.71 | 33.71 | 33.16 | 33.44 |
| 60 | 49.45 | 52.69 | 49.81 | 51.18 | 53.78 | 49.76 | 52.02 | 48.17 | 51.26 | 50.45 | 50.86 |
| 120 | 59.07 | 65.21 | 60.76 | 64.52 | 67.06 | 62.71 | 65.47 | 62.03 | 63.09 | 63.62 | 63.35 |
| 240 | 70.77 | 73.02 | 71.32 | 72.17 | 77.60 | 71.40 | 77.27 | 71.76 | 74.24 | 72.09 | 73.16 |
| $k_{obs}$ (M^-1 sec^-1) | 255.12 | 305.68 | 261.68 | 286.61 | 333.87 | 262.92 | 307.68 | 252.41 | 289.59 | 276.91 | 283.25 |
| $k_{adj}$ (M^-1 sec^-1) | 947.26 | 852.21 | 846.46 | 798.78 | 722.94 | 738.90 | 657.16 | 703.49 | 793.45 | 773.35 | 783.40 |
| Unreacted Fracrtion (%) | 27.42 | 22.43 | 25.28 | 22.63 | 17.62 | 23.04 | 17.52 | 22.94 | 21.96 | 22.76 | 22.36 |

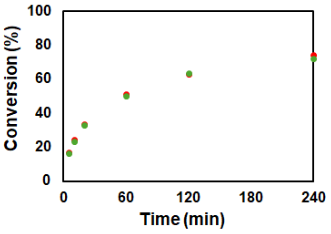

#### 50% TAF + 50% PRM-SH3$_{short}$ (total 100 µM; 5% PUMA), 0.5 µM mCh-ST-Bcl: 710.85 M$^{-1}$ s$^{-1}$

| Client Concentration = 4.347 µM | Conversion (%) | | | | | | | | | | |
|---|---|---|---|---|---|---|---|---|---|---|---|
| GFP client ratio = 1.45 [mCh] | Sample 1 - Gel 1 | | Sample 1 - Gel 2 | | Sample 2 - Gel 1 | | Sample 2 - Gel 2 | | Average | | |
| Reaction Time (min) | mCh-ST | GFP-SC | mCh-ST | GFP-SC | mCh-ST | GFP-SC | mCh-ST | GFP-SC | mCh-ST | GFP-SC | Total |
| 5 | 38.43 | 35.05 | 38.75 | 35.03 | 38.01 | 35.00 | 37.22 | 36.01 | 38.10 | 35.27 | 36.69 |
| 10 | 51.04 | 45.96 | 51.74 | 46.48 | 50.00 | 46.35 | 51.00 | 46.82 | 50.94 | 46.40 | 48.67 |
| 20 | 62.38 | 55.89 | 62.34 | 55.79 | 60.57 | 58.10 | 60.96 | 58.32 | 61.56 | 57.03 | 59.29 |
| 60 | 73.95 | 69.57 | 73.71 | 69.32 | 72.73 | 70.40 | 73.82 | 71.23 | 73.55 | 70.13 | 71.84 |
| 120 | 76.65 | 75.77 | 78.46 | 75.32 | 79.18 | 76.45 | 80.10 | 77.36 | 78.60 | 76.23 | 77.41 |
| 240 | 80.69 | 79.63 | 79.60 | 78.04 | 83.56 | 79.94 | 83.96 | 80.44 | 81.95 | 79.51 | 80.73 |
| $k_{obs}$ (M^-1 sec^-1) | 321.33 | 243.10 | 327.58 | 241.71 | 309.15 | 257.01 | 317.62 | 267.31 | 318.92 | 252.28 | 285.60 |
| $k_{adj}$ (M^-1 sec^-1) | 811.06 | 654.87 | 827.63 | 695.46 | 696.23 | 660.29 | 675.24 | 665.98 | 752.54 | 669.15 | 710.85 |
| Unreacted Fracrtion (%) | 18.71 | 20.07 | 18.68 | 21.16 | 16.73 | 19.33 | 15.76 | 18.72 | 17.47 | 19.82 | 18.64 |

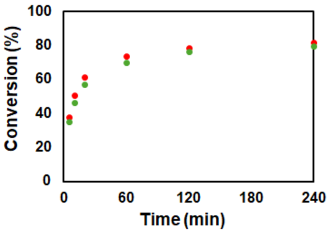

#### 10% FUS + 90% PRM-SH3$_{short}$ (total 100 µM; 5% PUMA), 0.1 µM mCh-ST-Bcl: 237.11 M$^{-1}$ s$^{-1}$

| Client Concentration = 5.602 µM | Conversion (%) | | | | | | | | | | |
|---|---|---|---|---|---|---|---|---|---|---|---|
| GFP client ratio = 0.725 [mCh] | Sample 1 - Gel 1 | | Sample 1 - Gel 2 | | Sample 2 - Gel 1 | | Sample 2 - Gel 2 | | Average | | |
| Reaction Time (min) | mCh-ST | GFP-SC | mCh-ST | GFP-SC | mCh-ST | GFP-SC | mCh-ST | GFP-SC | mCh-ST | GFP-SC | Total |
| 5 | 28.78 | 27.21 | 31.47 | 28.46 | 29.45 | 27.09 | 29.18 | 25.30 | 29.72 | 27.01 | 28.37 |
| 10 | 35.34 | 33.96 | 36.62 | 34.18 | 37.76 | 34.98 | 38.11 | 33.63 | 36.96 | 34.19 | 35.57 |
| 20 | 47.07 | 45.68 | 48.14 | 46.43 | 48.61 | 44.36 | 48.00 | 43.28 | 47.95 | 44.94 | 46.45 |
| 60 | 67.84 | 68.22 | 68.21 | 68.46 | 67.91 | 59.28 | 67.50 | 64.83 | 67.86 | 65.20 | 66.53 |
| 120 | 77.71 | 80.56 | 78.29 | 80.64 | 77.66 | 75.68 | 77.03 | 77.83 | 77.67 | 78.68 | 78.17 |
| 240 | 82.92 | 87.63 | 83.72 | 88.28 | 83.88 | 83.09 | 83.67 | 83.97 | 83.55 | 85.74 | 84.64 |
| $k_{obs}$ (M^-1 sec^-1) | 132.65 | 132.26 | 142.49 | 136.40 | 141.88 | 109.75 | 139.30 | 115.35 | 139.08 | 123.44 | 131.26 |
| $k_{adj}$ (M^-1 sec^-1) | 250.69 | 190.53 | 275.13 | 196.96 | 274.35 | 235.57 | 276.68 | 197.00 | 269.21 | 205.01 | 237.11 |
| Unreacted Fracrtion (%) | 14.19 | 8.61 | 14.45 | 8.62 | 14.53 | 16.45 | 15.03 | 12.26 | 14.55 | 11.49 | 13.02 |

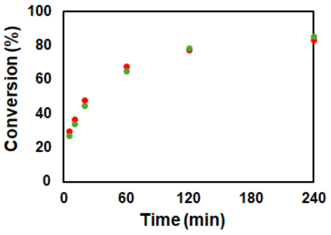

#### 25% FUS + 75% PRM-SH3$_{short}$ (total 100 µM; 5% PUMA), 0.1 µM mCh-ST-Bcl: 253.87 M$^{-1}$ s$^{-1}$

| Client Concentration = 2.831 µM | Conversion (%) | | | | | | | | | | |
|---|---|---|---|---|---|---|---|---|---|---|---|
| GFP client ratio = 1.0 [mCh] | Sample 1 - Gel 1 | | Sample 1 - Gel 2 | | Sample 2 - Gel 1 | | Sample 2 - Gel 2 | | Average | | |
| Reaction Time (min) | mCh-ST | GFP-SC | mCh-ST | GFP-SC | mCh-ST | GFP-SC | mCh-ST | GFP-SC | mCh-ST | GFP-SC | Total |
| 5 | 15.62 | 15.14 | 15.64 | 15.19 | 21.89 | 16.20 | 21.20 | 15.74 | 18.59 | 15.57 | 17.08 |
| 10 | 23.35 | 22.51 | 23.23 | 22.67 | 29.21 | 21.74 | 29.04 | 21.40 | 26.21 | 22.08 | 24.14 |
| 20 | 34.06 | 33.10 | 34.11 | 33.28 | 40.93 | 30.57 | 40.89 | 30.19 | 37.50 | 31.79 | 34.64 |
| 60 | 55.32 | 57.27 | 54.72 | 56.38 | 65.10 | 49.59 | 63.84 | 49.18 | 59.74 | 53.11 | 56.43 |
| 120 | 67.52 | 72.40 | 67.18 | 72.60 | 78.15 | 63.44 | 77.44 | 62.83 | 72.57 | 67.82 | 70.19 |
| 240 | 74.97 | 81.43 | 77.22 | 80.15 | 86.03 | 69.59 | 85.28 | 69.81 | 80.88 | 75.24 | 78.06 |
| $k_{obs}$ (M^-1 sec^-1) | 125.22 | 139.86 | 125.87 | 137.73 | 207.88 | 98.57 | 201.05 | 96.51 | 165.01 | 118.17 | 141.59 |
| $k_{adj}$ (M^-1 sec^-1) | 268.21 | 192.13 | 247.59 | 200.35 | 283.33 | 282.51 | 285.28 | 271.58 | 271.10 | 236.65 | 253.87 |
| Unreacted Fracrtion (%) | 17.71 | 8.07 | 15.94 | 9.42 | 7.56 | 23.56 | 8.49 | 23.30 | 12.43 | 16.09 | 14.26 |

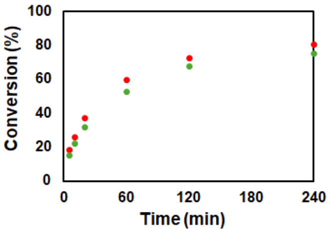

#### 50% FUS + 50% PRM-SH3$_{short}$ (total 100 µM; 5% PUMA), 0.1 µM mCh-ST-Bcl: 303.51 M$^{-1}$ s$^{-1}$

| Client Concentration = 1.464 µM | Conversion (%) | | | | | | | | | | |
|---|---|---|---|---|---|---|---|---|---|---|---|
| GFP client ratio = 1.5 [mCh] | Sample 1 - Gel 1 | | Sample 1 - Gel 2 | | Sample 2 - Gel 1 | | Sample 2 - Gel 2 | | Average | | |
| Reaction Time (min) | mCh-ST | GFP-SC | mCh-ST | GFP-SC | mCh-ST | GFP-SC | mCh-ST | GFP-SC | mCh-ST | GFP-SC | Total |
| 5 | 10.81 | 9.24 | 10.43 | 10.59 | 12.34 | 10.03 | 11.25 | 10.13 | 11.21 | 10.00 | 10.60 |
| 10 | 13.82 | 12.98 | 14.31 | 13.48 | 17.34 | 13.55 | 17.92 | 14.63 | 15.85 | 13.66 | 14.75 |
| 20 | 20.63 | 19.47 | 19.78 | 19.87 | 26.81 | 21.01 | 27.78 | 21.62 | 23.75 | 20.49 | 22.12 |
| 60 | 35.34 | 36.37 | 35.16 | 37.44 | 46.45 | 36.96 | 47.39 | 37.93 | 41.08 | 37.17 | 39.13 |
| 120 | 45.70 | 50.10 | 46.95 | 50.66 | 58.37 | 49.63 | 59.49 | 49.57 | 52.63 | 49.99 | 51.31 |
| 240 | 59.90 | 61.42 | 62.17 | 62.26 | 76.36 | 60.97 | 75.68 | 62.09 | 68.53 | 61.68 | 65.11 |
| $k_{obs}$ (M^-1 sec^-1) | 91.13 | 99.15 | 95.07 | 103.43 | 166.81 | 100.29 | 172.03 | 104.05 | 131.26 | 101.73 | 116.49 |
| $k_{adj}$ (M^-1 sec^-1) | 346.58 | 271.56 | 279.57 | 287.77 | 277.29 | 324.22 | 301.57 | 339.51 | 301.25 | 305.76 | 303.51 |
| Unreacted Fracrtion (%) | 31.78 | 25.57 | 26.99 | 25.64 | 13.05 | 28.54 | 14.21 | 28.46 | 21.51 | 27.05 | 24.28 |

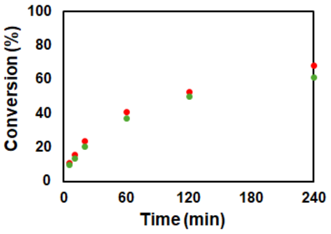

##### 50% FUS + 50% PRM-SH3$_{short}$ (total 100 µM; 5% PUMA), 0.5 µM mCh-ST-Bcl: 283.37 $M^{-1}$ $s^{-1}$

| Client Concentration = 6.533 µM | Conversion (%) | | | | | | | | | | |
|---|---|---|---|---|---|---|---|---|---|---|---|
| GFP client ratio = 1.5 [mCh] | Sample 1 - Gel 1 | | Sample 1 - Gel 2 | | Sample 2 - Gel 1 | | Sample 2 - Gel 2 | | Average | | |
| Reaction Time (min) | mCh-ST | GFP-SC | mCh-ST | GFP-SC | mCh-ST | GFP-SC | mCh-ST | GFP-SC | mCh-ST | GFP-SC | Total |
| 5 | 24.77 | 24.15 | 23.42 | 22.69 | 17.88 | 22.22 | 17.32 | 21.73 | 20.85 | 22.70 | 21.77 |
| 10 | 30.72 | 31.47 | 31.02 | 31.25 | 25.38 | 29.18 | 25.02 | 29.12 | 28.03 | 30.26 | 29.14 |
| 20 | 40.21 | 41.13 | 40.67 | 41.03 | 33.40 | 37.68 | 32.78 | 37.90 | 36.77 | 39.43 | 38.10 |
| 60 | 54.98 | 57.68 | 55.29 | 57.60 | 47.59 | 56.47 | 46.95 | 55.92 | 51.20 | 56.92 | 54.06 |
| 120 | 61.40 | 63.23 | 62.27 | 63.87 | 52.55 | 60.62 | 52.06 | 60.18 | 57.07 | 61.98 | 59.52 |
| 240 | 68.82 | 68.71 | 70.68 | 69.25 | 59.42 | 68.63 | 58.87 | 67.82 | 64.45 | 68.60 | 66.53 |
| kobs (M^-1 sec^-1) | 58.29 | 63.83 | 60.44 | 63.67 | 33.43 | 55.01 | 32.17 | 53.70 | 46.08 | 59.05 | 52.57 |
| kadj (M^-1 sec^-1) | 307.90 | 300.60 | 278.77 | 277.25 | 292.95 | 257.49 | 287.76 | 264.22 | 291.85 | 274.89 | 283.37 |
| Unreacted Fracrtion (%) | 31.31 | 29.81 | 29.46 | 28.81 | 39.58 | 30.10 | 40.02 | 30.85 | 35.09 | 29.90 | 32.49 |

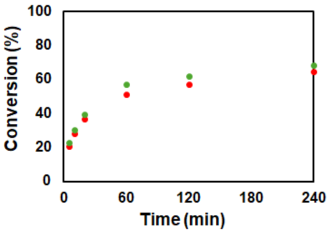

#### - Bulk Solutions

##### 30 µM Clients: 91.99 $M^{-1}$ $s^{-1}$

| | Conversion (%) | | | | | | | | | | |
|---|---|---|---|---|---|---|---|---|---|---|---|
| | Sample 1 - Gel 1 | | Sample 1 - Gel 2 | | Sample 2 - Gel 1 | | Sample 2 - Gel 2 | | Average | | |
| Reaction Time (min) | mCh-ST | GFP-SC | mCh-ST | GFP-SC | mCh-ST | GFP-SC | mCh-ST | GFP-SC | mCh-ST | GFP-SC | Total |
| 1 | 14.71 | 13.06 | 15.34 | 13.64 | 16.12 | 13.69 | 17.16 | 13.91 | 15.83 | 13.57 | 14.70 |
| 2 | 25.38 | 22.12 | 24.59 | 21.69 | 25.27 | 21.45 | 24.73 | 21.76 | 24.99 | 21.75 | 23.37 |
| 5 | 46.64 | 41.36 | 46.51 | 41.58 | 44.90 | 38.98 | 45.43 | 39.95 | 45.87 | 40.47 | 43.17 |
| 10 | 62.60 | 56.56 | 62.17 | 56.16 | 60.85 | 55.20 | 61.32 | 55.42 | 61.73 | 55.83 | 58.78 |
| 20 | 75.50 | 70.05 | 75.23 | 70.07 | 74.24 | 68.91 | 74.93 | 69.47 | 74.98 | 69.62 | 72.30 |
| 60 | 89.97 | 85.12 | 89.82 | 85.44 | 88.06 | 85.17 | 88.45 | 85.42 | 89.07 | 85.29 | 87.18 |
| 120 | 93.37 | 89.24 | 92.87 | 89.97 | 92.57 | 90.56 | 92.79 | 90.95 | 92.90 | 90.18 | 91.54 |
| 240 | 96.06 | 92.73 | 96.05 | 93.01 | 95.62 | 94.28 | 95.50 | 94.79 | 95.81 | 93.70 | 94.76 |
| kobs (M^-1 sec^-1) | 92.55 | 71.11 | 91.25 | 71.23 | 87.49 | 67.46 | 89.41 | 69.30 | 90.17 | 69.78 | 79.97 |
| kadj (M^-1 sec^-1) | 100.14 | 89.16 | 99.54 | 87.49 | 99.06 | 79.28 | 100.69 | 80.54 | 99.86 | 84.12 | 91.99 |
| Unreacted Fracrtion (%) | 1.98 | 5.55 | 2.18 | 5.06 | 3.07 | 4.01 | 2.94 | 3.73 | 2.54 | 4.59 | 3.57 |

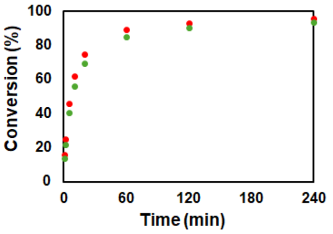

##### 20 µM Clients: 87.20 $M^{-1}$ $s^{-1}$

| | Conversion (%) | | | | | | | | | | |
|---|---|---|---|---|---|---|---|---|---|---|---|
| | Sample 1 - Gel 1 | | Sample 1 - Gel 2 | | Sample 2 - Gel 1 | | Sample 2 - Gel 2 | | Average | | |
| Reaction Time (min) | mCh-ST | GFP-SC | mCh-ST | GFP-SC | mCh-ST | GFP-SC | mCh-ST | GFP-SC | mCh-ST | GFP-SC | Total |
| 1 | 9.49 | 8.94 | 9.69 | 9.76 | 9.08 | 8.68 | 9.86 | 8.67 | 9.53 | 9.01 | 9.27 |
| 2 | 16.62 | 15.59 | 17.16 | 16.19 | 17.36 | 15.37 | 17.69 | 15.40 | 17.21 | 15.64 | 16.42 |
| 5 | 34.39 | 31.24 | 35.35 | 32.39 | 33.92 | 29.89 | 34.92 | 30.36 | 34.64 | 30.97 | 32.81 |
| 10 | 50.49 | 45.99 | 51.71 | 47.53 | 49.47 | 44.68 | 51.94 | 46.14 | 50.90 | 46.08 | 48.49 |
| 20 | 66.02 | 61.61 | 65.52 | 61.70 | 64.00 | 59.82 | 67.00 | 62.11 | 65.63 | 61.31 | 63.47 |
| 60 | 83.44 | 81.27 | 83.65 | 81.84 | 82.35 | 80.64 | 83.91 | 81.59 | 83.34 | 81.33 | 82.33 |
| 120 | 89.17 | 88.11 | 89.43 | 88.43 | 89.08 | 88.20 | 89.77 | 88.76 | 89.36 | 88.38 | 88.87 |
| 240 | 93.57 | 93.10 | 93.39 | 92.86 | 93.41 | 93.15 | 93.47 | 93.66 | 93.46 | 93.19 | 93.33 |
| kobs (M^-1 sec^-1) | 81.88 | 69.23 | 84.09 | 72.33 | 78.33 | 65.35 | 85.91 | 69.41 | 82.55 | 69.08 | 75.81 |
| kadj (M^-1 sec^-1) | 93.82 | 79.92 | 97.57 | 84.86 | 91.30 | 74.28 | 98.47 | 77.39 | 95.29 | 79.11 | 87.20 |
| Unreacted Fracrtion (%) | 3.47 | 3.65 | 3.77 | 4.03 | 3.87 | 3.26 | 3.48 | 2.79 | 3.65 | 3.43 | 3.54 |

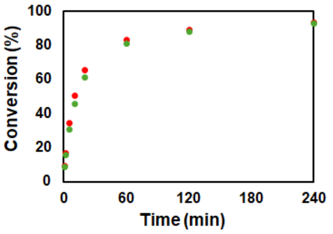

##### 10 µM Clients: 79.45 $M^{-1}$ $s^{-1}$

| | Conversion (%) | | | | | | | | | | |
|---|---|---|---|---|---|---|---|---|---|---|---|
| | Sample 1 - Gel 1 | | Sample 1 - Gel 2 | | Sample 2 - Gel 1 | | Sample 2 - Gel 2 | | Average | | |
| Reaction Time (min) | mCh-ST | GFP-SC | mCh-ST | GFP-SC | mCh-ST | GFP-SC | mCh-ST | GFP-SC | mCh-ST | GFP-SC | Total |
| 1 | 5.35 | 6.29 | 5.73 | 5.81 | 4.74 | 5.18 | 4.81 | 5.20 | 5.16 | 5.62 | 5.39 |
| 2 | 9.07 | 9.34 | 9.50 | 8.84 | 9.22 | 8.98 | 10.11 | 9.40 | 9.47 | 9.14 | 9.31 |
| 5 | 19.39 | 18.15 | 19.98 | 18.15 | 18.77 | 17.05 | 19.33 | 17.39 | 19.37 | 17.68 | 18.52 |
| 10 | 32.46 | 29.82 | 32.09 | 28.38 | 31.24 | 28.14 | 31.42 | 28.23 | 31.80 | 28.64 | 30.22 |
| 20 | 48.76 | 44.67 | 47.27 | 43.17 | 45.96 | 41.73 | 46.94 | 42.12 | 47.23 | 42.92 | 45.08 |
| 60 | 73.01 | 69.88 | 71.21 | 68.92 | 69.87 | 66.77 | 71.38 | 67.59 | 71.37 | 68.29 | 69.83 |
| 120 | 81.60 | 80.71 | 81.79 | 80.91 | 80.77 | 78.53 | 81.63 | 79.04 | 81.45 | 79.80 | 80.62 |
| 240 | 88.44 | 89.59 | 88.88 | 89.67 | 88.37 | 87.97 | 89.09 | 88.57 | 88.69 | 88.95 | 88.82 |
| kobs (M^-1 sec^-1) | 76.22 | 67.43 | 73.83 | 64.41 | 69.35 | 59.43 | 72.75 | 61.00 | 73.04 | 63.07 | 68.05 |
| kadj (M^-1 sec^-1) | 91.14 | 76.26 | 88.01 | 70.22 | 83.61 | 70.60 | 84.99 | 70.75 | 86.94 | 71.96 | 79.45 |
| Unreacted Fracrtion (%) | 4.61 | 3.19 | 4.50 | 2.26 | 4.79 | 4.43 | 4.01 | 3.83 | 4.48 | 3.43 | 3.95 |

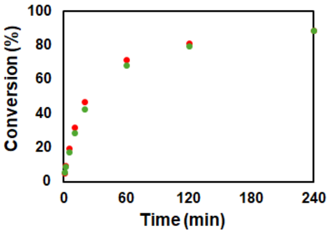

##### 5 µM Clients: 72.98 $M^{-1}$ $s^{-1}$

| | Conversion (%) | | | | | | | | | | |
|---|---|---|---|---|---|---|---|---|---|---|---|
| | Sample 1 - Gel 1 | | Sample 1 - Gel 2 | | Sample 2 - Gel 1 | | Sample 2 - Gel 2 | | Average | | |
| Reaction Time (min) | mCh-ST | GFP-SC | mCh-ST | GFP-SC | mCh-ST | GFP-SC | mCh-ST | GFP-SC | mCh-ST | GFP-SC | Total |
| 1 | 2.72 | 3.67 | 2.72 | 3.53 | 1.88 | 2.75 | 2.25 | 2.63 | 2.39 | 3.15 | 2.77 |
| 2 | 4.44 | 4.88 | 4.54 | 4.83 | 4.27 | 4.80 | 4.96 | 4.03 | 4.55 | 4.64 | 4.59 |
| 5 | 10.70 | 9.77 | 10.60 | 9.73 | 9.57 | 8.85 | 10.42 | 8.09 | 10.32 | 9.11 | 9.72 |
| 10 | 19.07 | 16.13 | 18.69 | 16.85 | 16.43 | 13.88 | 18.41 | 13.55 | 18.15 | 15.10 | 16.63 |
| 20 | 30.40 | 26.13 | 30.96 | 27.07 | 26.84 | 22.83 | 30.43 | 22.59 | 29.66 | 24.65 | 27.16 |
| 60 | 56.98 | 51.35 | 56.91 | 51.68 | 54.51 | 48.33 | 59.64 | 48.24 | 57.01 | 49.90 | 53.46 |
| 120 | 70.63 | 67.02 | 70.96 | 67.31 | 66.37 | 60.76 | 71.68 | 60.72 | 69.91 | 63.95 | 66.93 |
| 240 | 82.20 | 79.85 | 82.13 | 80.16 | 80.26 | 76.04 | 81.58 | 74.98 | 81.54 | 77.76 | 79.65 |
| kobs (M^-1 sec^-1) | 71.81 | 58.76 | 72.18 | 60.20 | 61.37 | 48.16 | 74.40 | 47.22 | 69.94 | 53.59 | 61.76 |
| kadj (M^-1 sec^-1) | 82.95 | 66.60 | 83.37 | 69.56 | 71.78 | 61.60 | 85.81 | 62.12 | 80.98 | 64.97 | 72.98 |
| Unreacted Fracrtion (%) | 3.91 | 3.49 | 3.91 | 3.99 | 4.33 | 6.87 | 3.89 | 7.65 | 4.01 | 5.50 | 4.75 |

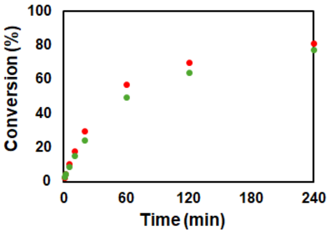

#### 2 µM Clients: 65.18 $M^{-1}$ $s^{-1}$

| | Conversion (%) | | | | | | | | | | |
|---|---|---|---|---|---|---|---|---|---|---|---|
| | Sample 1 - Gel 1 | | Sample 1 - Gel 2 | | Sample 2 - Gel 1 | | Sample 2 - Gel 2 | | Average | | |
| Reaction Time (min) | mCh-ST | GFP-SC | mCh-ST | GFP-SC | mCh-ST | GFP-SC | mCh-ST | GFP-SC | mCh-ST | GFP-SC | Total |
| 2 | 2.16 | 3.43 | 2.45 | 3.54 | 1.71 | 2.27 | 1.73 | 2.36 | 2.01 | 2.90 | 2.46 |
| 5 | 4.25 | 4.74 | 4.05 | 5.18 | 3.28 | 3.63 | 3.74 | 3.79 | 3.83 | 4.34 | 4.08 |
| 10 | 7.81 | 7.71 | 7.65 | 7.92 | 6.48 | 6.37 | 6.67 | 6.12 | 7.15 | 7.03 | 7.09 |
| 20 | 14.46 | 13.47 | 14.50 | 13.17 | 12.23 | 11.28 | 12.38 | 11.37 | 13.39 | 12.32 | 12.86 |
| 60 | 31.54 | 28.31 | 31.73 | 28.31 | 30.39 | 27.15 | 30.33 | 27.57 | 31.00 | 27.84 | 29.42 |
| 120 | 48.03 | 42.82 | 47.81 | 43.13 | 43.33 | 39.35 | 44.32 | 40.74 | 45.87 | 41.51 | 43.69 |
| 240 | 64.76 | 60.29 | 64.61 | 60.19 | 61.43 | 58.46 | 62.48 | 59.40 | 63.32 | 59.59 | 61.45 |
| kobs (M^-1 sec^-1) | 64.75 | 54.22 | 64.54 | 54.37 | 56.01 | 48.56 | 57.71 | 50.48 | 60.75 | 51.91 | 56.33 |
| kadj (M^-1 sec^-1) | 70.74 | 70.59 | 71.83 | 71.79 | 64.47 | 55.40 | 61.60 | 55.02 | 67.16 | 63.20 | 65.18 |
| Unreacted Fracrtion (%) | 2.89 | 8.52 | 3.48 | 8.94 | 4.66 | 4.50 | 2.20 | 2.95 | 3.31 | 6.23 | 4.77 |

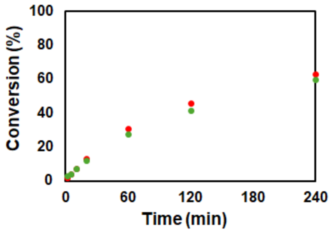

#### 1 µM Client (low concentration)

| | Conversion (%) | | | | | | | | | | |
|---|---|---|---|---|---|---|---|---|---|---|---|
| | Sample 1 - Gel 1 | | Sample 1 - Gel 2 | | Sample 2 - Gel 1 | | Sample 2 - Gel 2 | | Average | | |
| Reaction Time (min) | mCh-ST | GFP-SC | mCh-ST | GFP-SC | mCh-ST | GFP-SC | mCh-ST | GFP-SC | mCh-ST | GFP-SC | Total |
| 5 | 2.02 | 3.08 | 2.16 | 2.89 | 2.72 | 2.62 | 1.63 | 1.80 | 2.13 | 2.60 | 2.36 |
| 10 | 3.66 | 4.15 | 3.60 | 3.87 | 3.76 | 3.39 | 2.84 | 2.91 | 3.47 | 3.58 | 3.52 |
| 20 | 6.69 | 6.55 | 6.69 | 6.52 | 5.51 | 5.61 | 5.33 | 4.44 | 6.05 | 5.78 | 5.92 |
| 60 | 17.61 | 14.92 | 17.34 | 14.94 | 13.18 | 13.29 | 15.14 | 11.83 | 15.82 | 13.74 | 14.78 |
| 120 | 29.98 | 25.25 | 30.22 | 25.18 | 22.84 | 22.78 | 26.52 | 21.04 | 27.39 | 23.56 | 25.48 |
| 240 | 44.71 | 38.59 | 43.70 | 38.20 | 32.75 | 32.69 | 39.99 | 32.32 | 40.29 | 35.45 | 37.87 |
| kobs (M^-1 sec^-1) | 57.93 | 45.92 | 56.86 | 45.44 | 37.39 | 37.32 | 47.98 | 34.94 | 50.04 | 40.90 | 45.47 |
| kadj (M^-1 sec^-1) | 75.32 | 93.54 | 86.05 | 97.54 | 142.13 | 142.91 | 67.72 | 78.47 | 92.81 | 103.12 | 97.96 |
| Unreacted Fracrtion (%) | 9.57 | 24.13 | 14.60 | 25.63 | 40.40 | 40.57 | 12.70 | 27.94 | 19.32 | 29.57 | 24.44 |

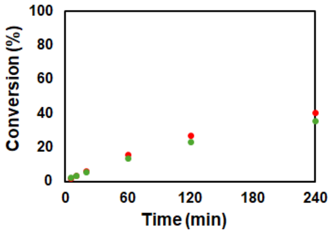

#### 0.5 µM Clients (low concentration)

| | Conversion (%) | | | | | | | | | | |
|---|---|---|---|---|---|---|---|---|---|---|---|
| | Sample 1 - Gel 1 | | Sample 1 - Gel 2 | | Sample 2 - Gel 1 | | Sample 2 - Gel 2 | | Average | | |
| Reaction Time (min) | mCh-ST | GFP-SC | mCh-ST | GFP-SC | mCh-ST | GFP-SC | mCh-ST | GFP-SC | mCh-ST | GFP-SC | Total |
| 10 | 1.63 | 1.39 | 1.06 | 1.75 | 1.25 | 1.88 | 1.47 | 1.66 | 1.35 | 1.67 | 1.51 |
| 20 | 2.03 | 2.61 | 2.62 | 2.38 | 2.28 | 2.40 | 1.93 | 2.01 | 2.21 | 2.35 | 2.28 |
| 60 | 6.90 | 4.44 | 5.80 | 5.27 | 5.20 | 4.71 | 5.25 | 4.24 | 5.79 | 4.66 | 5.23 |
| 120 | 9.99 | 7.89 | 9.46 | 10.09 | 8.96 | 7.49 | 9.23 | 7.50 | 9.41 | 8.24 | 8.83 |
| 240 | 14.48 | 14.97 | 14.58 | 15.82 | 16.48 | 12.92 | 16.20 | 13.54 | 15.43 | 14.31 | 14.87 |
| kobs (M^-1 sec^-1) | 26.61 | 24.60 | 25.87 | 27.86 | 27.75 | 21.74 | 27.57 | 22.29 | 26.95 | 24.12 | 25.54 |

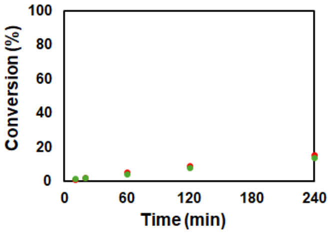

#### 0.25 µM Clients (low concentration)

| | Conversion (%) | | | | | | | | | | |
|---|---|---|---|---|---|---|---|---|---|---|---|
| | Sample 1 - Gel 1 | | Sample 1 - Gel 2 | | Sample 2 - Gel 1 | | Sample 2 - Gel 2 | | Average | | |
| Reaction Time (min) | mCh-ST | GFP-SC | mCh-ST | GFP-SC | mCh-ST | GFP-SC | mCh-ST | GFP-SC | mCh-ST | GFP-SC | Total |
| 60 | 1.88 | 1.94 | 2.69 | 2.37 | 2.62 | 2.01 | 1.57 | 1.66 | 2.19 | 2.00 | 2.09 |
| 120 | 3.56 | 3.26 | 4.24 | 3.71 | 4.07 | 3.99 | 2.48 | 2.65 | 3.58 | 3.40 | 3.49 |
| 240 | 6.15 | 5.93 | 7.13 | 6.56 | 6.60 | 5.75 | 5.55 | 5.54 | 6.36 | 5.95 | 6.15 |
| kobs (M^-1 sec^-1) | 18.84 | 18.01 | 22.56 | 20.33 | 21.03 | 18.53 | 15.94 | 16.19 | 19.59 | 18.27 | 18.93 |

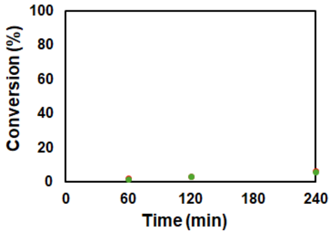

#### 5 µM Clients + 1% PEG8000: 91.06 $M^{-1}$ $s^{-1}$

| | Conversion (%) | | | | | | | | | | |
|---|---|---|---|---|---|---|---|---|---|---|---|
| | Sample 1 - Gel 1 | | Sample 1 - Gel 2 | | Sample 2 - Gel 1 | | Sample 2 - Gel 2 | | Average | | |
| Reaction Time (min) | mCh-ST | GFP-SC | mCh-ST | GFP-SC | mCh-ST | GFP-SC | mCh-ST | GFP-SC | mCh-ST | GFP-SC | Total |
| 1 | 3.24 | 4.05 | 3.24 | 3.85 | 2.44 | 3.20 | 2.23 | 2.86 | 2.79 | 3.49 | 3.14 |
| 2 | 6.15 | 6.26 | 5.99 | 5.92 | 4.78 | 5.18 | 5.03 | 5.12 | 5.49 | 5.62 | 5.55 |
| 5 | 12.93 | 11.75 | 12.50 | 11.55 | 11.63 | 10.57 | 11.68 | 10.69 | 12.18 | 11.14 | 11.66 |
| 10 | 23.39 | 20.31 | 22.34 | 19.61 | 20.92 | 18.54 | 20.97 | 18.84 | 21.90 | 19.32 | 20.61 |
| 20 | 35.93 | 31.77 | 35.36 | 31.49 | 33.08 | 29.74 | 34.61 | 30.25 | 34.74 | 30.81 | 32.78 |
| 60 | 61.96 | 57.19 | 61.52 | 56.57 | 58.15 | 54.67 | 59.72 | 56.07 | 60.34 | 56.13 | 58.23 |
| 120 | 75.03 | 71.92 | 74.96 | 72.48 | 72.38 | 70.65 | 72.40 | 70.76 | 73.69 | 71.45 | 72.57 |
| 240 | 85.04 | 83.90 | 84.83 | 84.06 | 82.63 | 82.44 | 82.86 | 82.75 | 83.84 | 83.29 | 83.56 |
| kobs (M^-1 sec^-1) | 91.54 | 76.55 | 89.16 | 75.65 | 78.30 | 69.14 | 81.57 | 71.21 | 85.14 | 73.14 | 79.14 |
| kadj (M^-1 sec^-1) | 106.65 | 86.69 | 101.69 | 81.99 | 94.66 | 76.65 | 100.59 | 79.58 | 100.89 | 81.23 | 91.06 |
| Unreacted Fracrtion (%) | 4.03 | 3.34 | 3.50 | 2.19 | 5.04 | 2.82 | 5.53 | 3.03 | 4.53 | 2.85 | 3.69 |

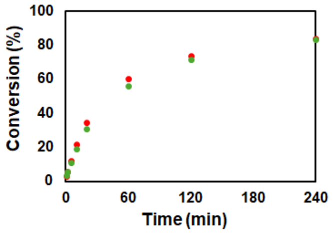

##### 5 µM Clients + 2.5% PEG8000: 115.41 $M^{-1}$ $s^{-1}$

| | Conversion (%) | | | | | | | | | | |
|---|---|---|---|---|---|---|---|---|---|---|---|
| | Sample 1 - Gel 1 | | Sample 1 - Gel 2 | | Sample 2 - Gel 1 | | Sample 2 - Gel 2 | | Average | | |
| Reaction Time (min) | mCh-ST | GFP-SC | mCh-ST | GFP-SC | mCh-ST | GFP-SC | mCh-ST | GFP-SC | mCh-ST | GFP-SC | Total |
| 1 | 4.30 | 4.64 | 4.33 | 4.77 | 3.46 | 4.42 | 3.56 | 4.20 | 3.91 | 4.51 | 4.21 |
| 2 | 7.27 | 6.95 | 7.57 | 7.50 | 6.74 | 6.50 | 6.58 | 6.63 | 7.04 | 6.89 | 6.97 |
| 5 | 15.92 | 13.78 | 16.66 | 14.37 | 14.03 | 12.77 | 14.85 | 13.45 | 15.37 | 13.59 | 14.48 |
| 10 | 28.38 | 24.42 | 28.94 | 23.99 | 25.12 | 22.15 | 25.37 | 22.28 | 26.95 | 23.21 | 25.08 |
| 20 | 40.94 | 36.07 | 41.67 | 36.20 | 39.34 | 34.86 | 40.26 | 35.23 | 40.56 | 35.59 | 38.07 |
| 60 | 67.41 | 62.99 | 67.93 | 62.23 | 65.91 | 62.27 | 65.49 | 61.50 | 66.69 | 62.25 | 64.47 |
| 120 | 79.55 | 76.69 | 80.82 | 77.74 | 77.82 | 75.93 | 77.34 | 76.03 | 78.88 | 76.60 | 77.74 |
| 240 | 88.02 | 87.23 | 88.38 | 87.56 | 85.16 | 86.15 | 84.89 | 86.73 | 86.61 | 86.92 | 86.76 |
| $k_{obs}$ (M^-1 sec^-1) | 118.07 | 97.31 | 122.74 | 97.78 | 105.36 | 91.26 | 106.22 | 91.65 | 113.10 | 94.50 | 103.80 |
| $k_{adj}$ (M^-1 sec^-1) | 132.47 | 104.61 | 135.56 | 102.34 | 124.13 | 96.83 | 130.77 | 96.55 | 130.73 | 100.08 | 115.41 |
| Unreacted Fracrtion (%) | 3.02 | 1.93 | 2.61 | 1.22 | 4.31 | 1.60 | 5.40 | 1.40 | 3.83 | 1.54 | 2.69 |

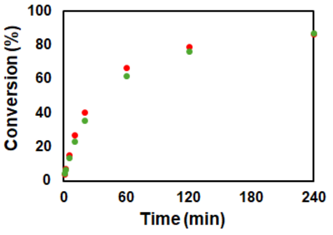

##### 5 µM Clients + 5% PEG8000: 170.44 $M^{-1}$ $s^{-1}$

| | Conversion (%) | | | | | | | | | | |
|---|---|---|---|---|---|---|---|---|---|---|---|
| | Sample 1 - Gel 1 | | Sample 1 - Gel 2 | | Sample 2 - Gel 1 | | Sample 2 - Gel 2 | | Average | | |
| Reaction Time (min) | mCh-ST | GFP-SC | mCh-ST | GFP-SC | mCh-ST | GFP-SC | mCh-ST | GFP-SC | mCh-ST | GFP-SC | Total |
| 1 | 5.61 | 5.65 | 6.56 | 6.81 | 4.93 | 5.60 | 4.82 | 5.34 | 5.48 | 5.85 | 5.66 |
| 2 | 10.75 | 8.90 | 13.88 | 11.30 | 8.92 | 8.34 | 8.34 | 8.17 | 10.47 | 9.18 | 9.83 |
| 5 | 21.90 | 17.74 | 25.47 | 20.37 | 19.56 | 17.15 | 18.82 | 16.40 | 21.44 | 17.92 | 19.68 |
| 10 | 35.91 | 29.30 | 39.69 | 32.59 | 31.19 | 27.23 | 31.08 | 26.97 | 34.47 | 29.02 | 31.74 |
| 20 | 50.30 | 43.08 | 53.36 | 45.97 | 47.05 | 42.05 | 45.97 | 40.65 | 49.17 | 42.94 | 46.05 |
| 60 | 74.39 | 69.43 | 80.03 | 74.61 | 71.34 | 68.36 | 70.77 | 67.50 | 74.13 | 69.98 | 72.05 |
| 120 | 83.65 | 80.80 | 86.62 | 83.76 | 80.74 | 79.58 | 79.96 | 79.30 | 82.74 | 80.86 | 81.80 |
| 240 | 89.44 | 89.46 | 90.73 | 91.18 | 87.84 | 88.91 | 87.03 | 88.19 | 88.76 | 89.43 | 89.10 |
| $k_{obs}$ (M^-1 sec^-1) | 171.55 | 130.02 | 210.41 | 156.54 | 143.25 | 121.89 | 137.81 | 116.88 | 165.76 | 131.33 | 148.54 |
| $k_{adj}$ (M^-1 sec^-1) | 204.23 | 143.41 | 237.41 | 165.37 | 175.55 | 134.73 | 173.03 | 129.81 | 197.55 | 143.33 | 170.44 |
| Unreacted Fracrtion (%) | 4.46 | 2.56 | 3.11 | 1.44 | 5.21 | 2.63 | 5.81 | 2.75 | 4.65 | 2.34 | 3.50 |

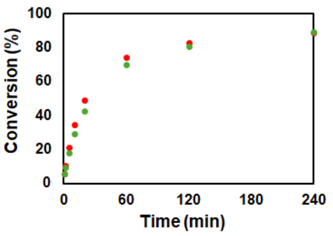

##### 5 µM Clients + 10% PEG8000: 295.63 $M^{-1}$ $s^{-1}$

| | Conversion (%) | | | | | | | | | | |
|---|---|---|---|---|---|---|---|---|---|---|---|
| | Sample 1 - Gel 1 | | Sample 1 - Gel 2 | | Sample 2 - Gel 1 | | Sample 2 - Gel 2 | | Average | | |
| Reaction Time (min) | mCh-ST | GFP-SC | mCh-ST | GFP-SC | mCh-ST | GFP-SC | mCh-ST | GFP-SC | mCh-ST | GFP-SC | Total |
| 1 | 11.04 | 9.69 | 11.16 | 9.74 | 8.88 | 7.16 | 9.16 | 7.32 | 10.06 | 8.48 | 9.27 |
| 2 | 19.62 | 15.73 | 20.27 | 16.15 | 14.94 | 12.64 | 15.49 | 12.30 | 17.58 | 14.21 | 15.89 |
| 5 | 31.91 | 28.75 | 32.70 | 29.46 | 29.32 | 25.37 | 31.17 | 26.03 | 31.28 | 27.40 | 29.34 |
| 10 | 49.31 | 40.04 | 50.07 | 40.93 | 45.57 | 38.60 | 46.46 | 39.35 | 47.85 | 39.73 | 43.79 |
| 20 | 59.10 | 58.26 | 59.37 | 58.64 | 61.86 | 53.00 | 62.75 | 54.66 | 60.77 | 56.14 | 58.46 |
| 60 | 83.69 | 79.41 | 83.95 | 80.10 | 81.58 | 76.10 | 82.52 | 77.52 | 82.93 | 78.28 | 80.61 |
| 120 | 89.42 | 87.19 | 89.53 | 87.21 | 86.72 | 86.84 | 87.37 | 87.38 | 88.26 | 87.16 | 87.71 |
| 240 | 92.98 | 92.94 | 92.90 | 92.97 | 91.02 | 92.57 | 91.58 | 93.17 | 92.12 | 92.91 | 92.52 |
| $k_{obs}$ (M^-1 sec^-1) | 298.34 | 237.20 | 306.97 | 244.65 | 267.46 | 200.16 | 282.73 | 210.77 | 288.88 | 223.20 | 256.04 |
| $k_{adj}$ (M^-1 sec^-1) | 346.87 | 269.80 | 361.00 | 280.08 | 322.25 | 221.15 | 336.35 | 227.54 | 341.62 | 249.65 | 295.63 |
| Unreacted Fracrtion (%) | 3.76 | 3.27 | 4.02 | 3.43 | 4.74 | 2.56 | 4.42 | 1.98 | 4.23 | 2.81 | 3.52 |

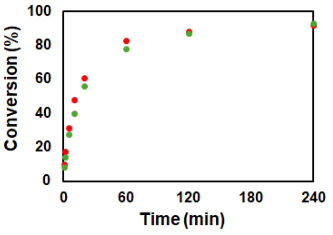

##### 5 µM Clients + 20% PEG8000: 1143.17 $M^{-1}$ $s^{-1}$

| | Conversion (%) | | | | | | | | | | |
|---|---|---|---|---|---|---|---|---|---|---|---|
| | Sample 1 - Gel 1 | | Sample 1 - Gel 2 | | Sample 2 - Gel 1 | | Sample 2 - Gel 2 | | Average | | |
| Reaction Time (min) | mCh-ST | GFP-SC | mCh-ST | GFP-SC | mCh-ST | GFP-SC | mCh-ST | GFP-SC | mCh-ST | GFP-SC | Total |
| 5 | 67.13 | 55.69 | 67.01 | 55.83 | 61.67 | 52.40 | 64.00 | 53.52 | 64.95 | 54.36 | 59.66 |
| 10 | 77.10 | 66.40 | 77.97 | 66.63 | 75.86 | 67.37 | 78.41 | 69.51 | 77.33 | 67.48 | 72.40 |
| 20 | 87.58 | 77.11 | 88.27 | 77.38 | 84.24 | 77.25 | 87.61 | 80.67 | 86.92 | 78.10 | 82.51 |
| 60 | 95.31 | 88.25 | 95.83 | 88.41 | 93.20 | 88.33 | 94.92 | 90.08 | 94.82 | 88.77 | 91.79 |
| 120 | 97.05 | 89.81 | 96.85 | 89.72 | 95.40 | 89.68 | 96.54 | 90.23 | 96.46 | 89.86 | 93.16 |
| 240 | 98.09 | 88.69 | 97.98 | 88.13 | 97.11 | 91.64 | 98.10 | 92.04 | 97.82 | 90.13 | 93.97 |
| $k_{obs}$ (M^-1 sec^-1) | 1252.14 | 676.15 | 1280.61 | 682.49 | 1021.72 | 650.71 | 1184.41 | 722.94 | 1184.72 | 683.07 | 933.90 |
| $k_{adj}$ (M^-1 sec^-1) | 1338.05 | 1066.63 | 1354.97 | 1090.05 | 1154.93 | 922.30 | 1244.36 | 974.06 | 1273.07 | 1013.26 | 1143.17 |
| Unreacted Fracrtion (%) | 1.11 | 8.61 | 0.94 | 8.80 | 2.21 | 6.90 | 0.86 | 5.84 | 1.28 | 7.54 | 4.41 |

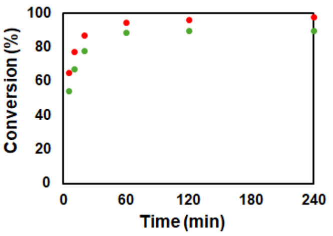

##### 2.5 µM Clients + 30% PEG8000: Reaction ended in 5 min (unreacted fraction ~ 40%)

| | Conversion (%) | | | | | | | | | | |
|---|---|---|---|---|---|---|---|---|---|---|---|
| | Sample 1 - Gel 1 | | Sample 1 - Gel 2 | | Sample 2 - Gel 1 | | Sample 2 - Gel 2 | | Average | | |
| Reaction Time (min) | mCh-ST | GFP-SC | mCh-ST | GFP-SC | mCh-ST | GFP-SC | mCh-ST | GFP-SC | mCh-ST | GFP-SC | Total |
| 5 | 56.19 | 53.58 | 61.11 | 53.66 | 60.52 | 55.34 | 54.93 | 53.18 | 58.19 | 53.94 | 56.07 |
| 10 | 58.29 | 55.68 | 61.29 | 55.58 | 59.35 | 55.44 | 58.14 | 56.66 | 59.27 | 55.84 | 57.55 |
| 20 | 60.21 | 57.33 | 60.04 | 55.30 | 59.54 | 54.28 | 58.40 | 57.72 | 59.55 | 56.16 | 57.85 |
| 60 | 60.29 | 57.67 | 59.94 | 54.97 | 58.25 | 53.50 | 59.69 | 57.27 | 59.54 | 55.85 | 57.70 |
| 120 | 58.74 | 56.07 | 60.05 | 54.42 | 57.09 | 53.65 | 58.20 | 55.93 | 58.52 | 55.02 | 56.77 |
| 240 | 55.81 | 51.89 | 54.86 | 49.72 | 56.28 | 50.40 | 52.64 | 49.91 | 54.89 | 50.48 | 52.69 |
| $k_{obs}$ (M^-1 sec^-1) | 586.88 | 445.32 | 700.03 | 383.11 | 627.46 | 368.08 | 531.61 | 448.75 | 611.50 | 411.31 | 511.41 |

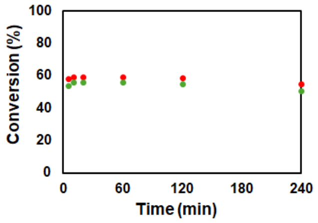

#### 1 µM Clients + 10% PEG8000: 341.22 $M^{-1}$ $s^{-1}$

| | Conversion (%) | | | | | | | | | | |
|---|---|---|---|---|---|---|---|---|---|---|---|
| | Sample 1 - Gel 1 | | Sample 1 - Gel 2 | | Sample 2 - Gel 1 | | Sample 2 - Gel 2 | | Average | | |
| Reaction Time (min) | mCh-ST | GFP-SC | mCh-ST | GFP-SC | mCh-ST | GFP-SC | mCh-ST | GFP-SC | mCh-ST | GFP-SC | Total |
| 5 | 7.72 | 7.40 | 9.26 | 8.74 | 8.44 | 7.90 | 8.43 | 7.09 | 8.46 | 7.78 | 8.12 |
| 10 | 17.12 | 13.72 | 15.69 | 14.98 | 16.64 | 13.06 | 15.59 | 12.02 | 16.26 | 13.44 | 14.85 |
| 20 | 26.45 | 21.29 | 25.49 | 22.17 | 22.30 | 20.21 | 21.10 | 19.19 | 23.83 | 20.72 | 22.27 |
| 60 | 55.97 | 48.04 | 49.74 | 45.98 | 45.08 | 41.41 | 44.72 | 40.22 | 48.88 | 43.91 | 46.39 |
| 120 | 65.05 | 59.87 | 62.36 | 59.66 | 57.18 | 57.45 | 55.99 | 55.26 | 60.15 | 58.06 | 59.10 |
| 240 | 74.45 | 76.24 | 72.99 | 72.56 | 70.67 | 71.28 | 70.67 | 70.07 | 72.19 | 72.54 | 72.37 |
| kobs (M^-1 sec^-1) | 291.26 | 232.07 | 251.77 | 221.50 | 209.67 | 193.83 | 201.89 | 180.07 | 238.65 | 206.87 | 222.76 |
| kadj (M^-1 sec^-1) | 451.69 | 272.10 | 417.17 | 336.44 | 385.66 | 266.13 | 352.54 | 247.99 | 401.76 | 280.67 | 341.22 |
| Unreacted Fracrtion (%) | 11.49 | 4.57 | 13.17 | 11.33 | 15.87 | 9.01 | 14.81 | 9.22 | 13.83 | 8.53 | 11.18 |

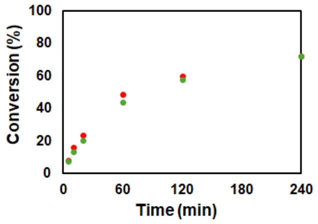

#### 1 µM Clients + 20% PEG8000: 1408.58 $M^{-1}$ $s^{-1}$

| | Conversion (%) | | | | | | | | | | |
|---|---|---|---|---|---|---|---|---|---|---|---|
| | Sample 1 - Gel 1 | | Sample 1 - Gel 2 | | Sample 2 - Gel 1 | | Sample 2 - Gel 2 | | Average | | |
| Reaction Time (min) | mCh-ST | GFP-SC | mCh-ST | GFP-SC | mCh-ST | GFP-SC | mCh-ST | GFP-SC | mCh-ST | GFP-SC | Total |
| 5 | 28.91 | 24.58 | 29.97 | 26.01 | 31.00 | 23.73 | 32.31 | 24.05 | 30.55 | 24.59 | 27.57 |
| 10 | 39.92 | 35.74 | 39.03 | 34.84 | 38.92 | 31.11 | 42.23 | 33.19 | 40.02 | 33.72 | 36.87 |
| 20 | 50.74 | 46.54 | 51.84 | 47.60 | 51.78 | 43.07 | 54.43 | 44.15 | 52.19 | 45.34 | 48.77 |
| 60 | 69.02 | 68.95 | 70.65 | 70.15 | 71.57 | 65.08 | 74.54 | 66.84 | 71.45 | 67.75 | 69.60 |
| 120 | 77.14 | 80.31 | 76.76 | 80.79 | 79.97 | 76.25 | 80.35 | 76.57 | 78.55 | 78.48 | 78.52 |
| 240 | 82.39 | 87.83 | 83.02 | 87.80 | 85.57 | 82.91 | 85.74 | 82.18 | 84.18 | 85.18 | 84.68 |
| kobs (M^-1 sec^-1) | 843.65 | 754.95 | 872.85 | 780.68 | 918.86 | 607.30 | 1050.03 | 647.67 | 921.35 | 697.65 | 809.50 |
| kadj (M^-1 sec^-1) | 1740.57 | 1075.03 | 1749.91 | 1100.44 | 1587.66 | 1042.41 | 1816.18 | 1156.42 | 1723.58 | 1093.58 | 1408.58 |
| Unreacted Fracrtion (%) | 15.81 | 8.40 | 15.27 | 8.18 | 12.30 | 12.51 | 12.27 | 13.30 | 13.91 | 10.60 | 12.26 |

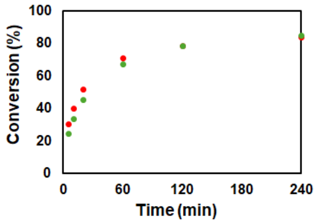

#### 1 µM Clients + 30% PEG8000

| | Conversion (%) | | | | | | | | | | |
|---|---|---|---|---|---|---|---|---|---|---|---|
| | Sample 1 - Gel 1 | | Sample 1 - Gel 2 | | Sample 2 - Gel 1 | | Sample 2 - Gel 2 | | Average | | |
| Reaction Time (min) | mCh-ST | GFP-SC | mCh-ST | GFP-SC | mCh-ST | GFP-SC | mCh-ST | GFP-SC | mCh-ST | GFP-SC | Total |
| 5 | 42.42 | 42.76 | 43.25 | 44.14 | 37.81 | 40.56 | 43.97 | 42.47 | 41.86 | 42.48 | 42.17 |
| 10 | 45.52 | 46.23 | 45.60 | 46.42 | 40.58 | 43.30 | 39.06 | 43.59 | 42.69 | 44.89 | 43.79 |
| 20 | 46.95 | 48.99 | 47.47 | 49.27 | 44.03 | 47.83 | 45.62 | 49.58 | 46.02 | 48.92 | 47.47 |
| 60 | 49.49 | 51.37 | 50.11 | 52.05 | 47.41 | 53.94 | 48.14 | 55.35 | 48.79 | 53.18 | 50.98 |
| 120 | 48.89 | 51.41 | 48.79 | 52.56 | 50.57 | 58.93 | 52.53 | 60.00 | 50.20 | 55.73 | 52.96 |
| 240 | 52.88 | 53.56 | 51.47 | 55.72 | 58.61 | 61.42 | 56.57 | 63.61 | 54.88 | 58.58 | 56.73 |
| kobs (M^-1 sec^-1) | 354.65 | 437.22 | 361.08 | 488.34 | 315.95 | 534.13 | 348.13 | 616.00 | 344.95 | 518.92 | 431.94 |

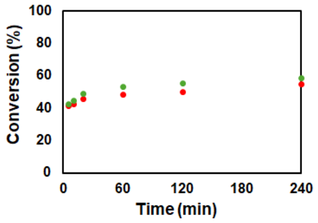

#### 5 µM Clients + 10% Ficoll 70: 268.10 $M^{-1}$ $s^{-1}$

| | Conversion (%) | | | | | | | | | | |
|---|---|---|---|---|---|---|---|---|---|---|---|
| | Sample 1 - Gel 1 | | Sample 1 - Gel 2 | | Sample 2 - Gel 1 | | Sample 2 - Gel 2 | | Average | | |
| Reaction Time (min) | mCh-ST | GFP-SC | mCh-ST | GFP-SC | mCh-ST | GFP-SC | mCh-ST | GFP-SC | mCh-ST | GFP-SC | Total |
| 1 | 9.23 | 7.66 | 8.36 | 7.62 | 7.19 | 6.53 | 7.26 | 7.38 | 8.01 | 7.30 | 7.66 |
| 2 | 15.39 | 12.77 | 14.84 | 12.40 | 14.30 | 12.41 | 14.49 | 12.91 | 14.76 | 12.62 | 13.69 |
| 5 | 30.41 | 25.49 | 28.64 | 24.16 | 28.92 | 24.89 | 29.33 | 25.67 | 29.33 | 25.05 | 27.19 |
| 10 | 45.04 | 38.61 | 42.98 | 36.61 | 42.15 | 36.56 | 42.71 | 37.89 | 43.22 | 37.42 | 40.32 |
| 20 | 60.48 | 53.66 | 58.35 | 51.78 | 57.31 | 51.05 | 57.84 | 52.14 | 58.50 | 52.16 | 55.33 |
| 60 | 80.64 | 75.20 | 79.53 | 73.88 | 78.27 | 73.42 | 77.89 | 73.21 | 79.08 | 73.93 | 76.50 |
| 120 | 87.52 | 83.30 | 86.75 | 82.40 | 86.50 | 82.42 | 86.35 | 82.42 | 86.78 | 82.64 | 84.71 |
| 240 | 92.81 | 89.14 | 92.23 | 88.74 | 91.88 | 88.59 | 91.27 | 88.93 | 92.05 | 88.85 | 90.45 |
| kobs (M^-1 sec^-1) | 265.59 | 194.83 | 243.01 | 179.39 | 233.64 | 177.35 | 236.86 | 184.93 | 244.77 | 184.12 | 214.45 |
| kadj (M^-1 sec^-1) | 310.49 | 254.93 | 285.64 | 235.99 | 279.04 | 235.44 | 291.68 | 251.62 | 291.71 | 244.50 | 268.10 |
| Unreacted Fracrtion (%) | 3.96 | 6.69 | 4.10 | 6.81 | 4.48 | 7.01 | 5.22 | 7.55 | 4.44 | 7.02 | 5.73 |

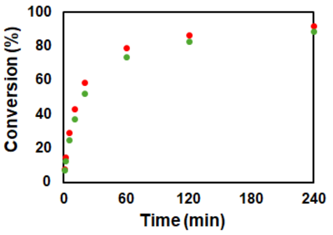

#### 5 µM Clients + 20% Ficoll 70: 788.14 $M^{-1}$ $s^{-1}$

| | Conversion (%) | | | | | | | | | | |
|---|---|---|---|---|---|---|---|---|---|---|---|
| | Sample 1 - Gel 1 | | Sample 1 - Gel 2 | | Sample 2 - Gel 1 | | Sample 2 - Gel 2 | | Average | | |
| Reaction Time (min) | mCh-ST | GFP-SC | mCh-ST | GFP-SC | mCh-ST | GFP-SC | mCh-ST | GFP-SC | mCh-ST | GFP-SC | Total |
| 1 | 20.60 | 19.05 | 19.62 | 18.58 | 18.38 | 16.59 | 19.09 | 17.23 | 19.42 | 17.86 | 18.64 |
| 2 | 30.25 | 29.19 | 30.53 | 29.16 | 30.17 | 27.09 | 31.05 | 28.27 | 30.50 | 28.43 | 29.46 |
| 5 | 50.56 | 50.47 | 50.49 | 49.76 | 48.79 | 45.31 | 50.19 | 46.84 | 50.01 | 48.10 | 49.05 |
| 10 | 64.74 | 66.12 | 64.71 | 65.96 | 63.29 | 60.84 | 64.70 | 62.55 | 64.36 | 63.87 | 64.11 |
| 20 | 75.35 | 79.34 | 76.03 | 80.29 | 74.98 | 74.12 | 75.42 | 75.16 | 75.45 | 77.23 | 76.34 |
| 60 | 85.48 | 91.62 | 85.45 | 91.56 | 85.71 | 88.14 | 86.82 | 88.91 | 85.87 | 90.06 | 87.96 |
| 120 | 87.73 | 94.92 | 87.69 | 94.45 | 88.94 | 91.79 | 89.93 | 92.32 | 88.57 | 93.37 | 90.97 |
| 240 | 90.05 | 96.12 | 89.92 | 96.19 | 91.67 | 94.90 | 91.86 | 95.11 | 90.88 | 95.58 | 93.23 |
| kobs (M^-1 sec^-1) | 628.28 | 674.14 | 628.61 | 668.85 | 594.53 | 536.21 | 631.48 | 574.53 | 620.73 | 613.43 | 617.08 |
| kadj (M^-1 sec^-1) | 937.25 | 723.63 | 931.08 | 715.02 | 831.76 | 631.66 | 863.71 | 670.99 | 890.95 | 685.32 | 788.14 |
| Unreacted Fracrtion (%) | 9.20 | 1.75 | 9.09 | 1.66 | 7.84 | 3.99 | 7.32 | 3.78 | 8.37 | 2.79 | 5.58 |

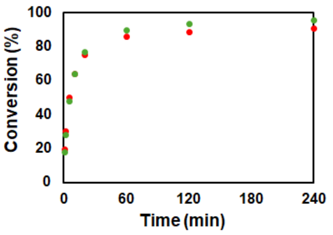

#### 5 μM Clients + 10% PEG400: 112.94 M$^{-1}$ s$^{-1}$

| | Conversion (%) | | | | | | | | | | |
|---|---|---|---|---|---|---|---|---|---|---|---|
| | Sample 1 - Gel 1 | | Sample 1 - Gel 2 | | Sample 2 - Gel 1 | | Sample 2 - Gel 2 | | Average | | |
| Reaction Time (min) | mCh-ST | GFP-SC | mCh-ST | GFP-SC | mCh-ST | GFP-SC | mCh-ST | GFP-SC | mCh-ST | GFP-SC | Total |
| 1 | 2.95 | 3.19 | 3.62 | 3.88 | 3.22 | 3.61 | 3.40 | 3.73 | 3.30 | 3.60 | 3.45 |
| 2 | 6.39 | 5.92 | 6.61 | 6.44 | 6.61 | 6.52 | 6.95 | 6.61 | 6.64 | 6.37 | 6.51 |
| 5 | 13.88 | 12.20 | 13.91 | 12.35 | 15.46 | 13.63 | 15.83 | 14.39 | 14.77 | 13.14 | 13.96 |
| 10 | 24.79 | 22.01 | 24.85 | 21.45 | 26.87 | 23.72 | 25.79 | 22.79 | 25.58 | 22.49 | 24.03 |
| 20 | 39.20 | 34.95 | 39.33 | 34.80 | 41.04 | 36.76 | 39.72 | 35.70 | 39.82 | 35.55 | 37.69 |
| 60 | 64.32 | 61.06 | 63.53 | 60.11 | 64.90 | 61.11 | 65.72 | 62.31 | 64.62 | 61.15 | 62.88 |
| 120 | 82.82 | 80.35 | 82.24 | 79.54 | 76.72 | 74.62 | 77.06 | 75.21 | 79.71 | 77.43 | 78.57 |
| 240 | 85.01 | 85.21 | 84.53 | 84.61 | 85.58 | 85.48 | 85.33 | 85.06 | 85.11 | 85.09 | 85.10 |
| $k_{obs}$ (M^-1 sec^-1) | 107.02 | 92.55 | 105.66 | 90.18 | 108.30 | 92.18 | 106.96 | 92.31 | 106.99 | 91.81 | 99.40 |
| $k_{adj}$ (M^-1 sec^-1) | 114.12 | 92.55 | 116.88 | 92.77 | 137.46 | 110.08 | 131.88 | 107.78 | 125.09 | 100.80 | 112.94 |
| Unreacted Fracrtion (%) | 1.73 | 0.00 | 2.69 | 0.77 | 6.11 | 4.63 | 5.42 | 4.08 | 3.99 | 2.37 | 3.18 |

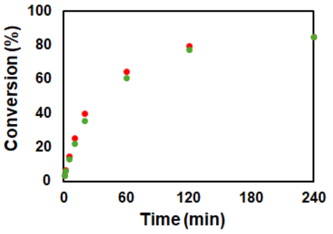

#### 5 μM Clients + 25% PEG400: 680.38 M$^{-1}$ s$^{-1}$

| | Conversion (%) | | | | | | | | | | |
|---|---|---|---|---|---|---|---|---|---|---|---|
| | Sample 1 - Gel 1 | | Sample 1 - Gel 2 | | Sample 2 - Gel 1 | | Sample 2 - Gel 2 | | Average | | |
| Reaction Time (min) | mCh-ST | GFP-SC | mCh-ST | GFP-SC | mCh-ST | GFP-SC | mCh-ST | GFP-SC | mCh-ST | GFP-SC | Total |
| 1 | 18.90 | 15.17 | 20.19 | 16.40 | 24.03 | 21.47 | 24.30 | 21.31 | 21.85 | 18.59 | 20.22 |
| 2 | 23.57 | 19.76 | 26.52 | 22.31 | 32.32 | 29.07 | 32.53 | 29.64 | 28.74 | 25.20 | 26.97 |
| 5 | 38.33 | 33.53 | 40.96 | 35.65 | 47.14 | 44.27 | 47.24 | 44.57 | 43.42 | 39.50 | 41.46 |
| 10 | 53.31 | 48.15 | 54.02 | 48.54 | 59.58 | 57.61 | 59.24 | 57.68 | 56.53 | 53.00 | 54.76 |
| 20 | 66.51 | 61.77 | 67.99 | 63.32 | 70.01 | 70.58 | 69.55 | 70.74 | 68.52 | 66.60 | 67.56 |
| 60 | 82.09 | 79.83 | 83.15 | 80.83 | 82.10 | 85.92 | 82.40 | 86.47 | 82.43 | 83.26 | 82.85 |
| 120 | 87.01 | 85.65 | 87.79 | 86.33 | 85.82 | 91.02 | 85.76 | 91.29 | 86.60 | 88.58 | 87.59 |
| 240 | 90.22 | 90.03 | 90.65 | 90.41 | 88.08 | 94.82 | 87.99 | 94.60 | 89.23 | 92.46 | 90.85 |
| $k_{obs}$ (M^-1 sec^-1) | 384.46 | 299.51 | 425.47 | 325.37 | 545.65 | 509.45 | 544.12 | 516.64 | 474.93 | 412.74 | 443.83 |
| $k_{adj}$ (M^-1 sec^-1) | 566.24 | 428.47 | 632.45 | 468.82 | 1004.43 | 662.47 | 1010.61 | 669.53 | 803.43 | 557.32 | 680.38 |
| Unreacted Fracrtion (%) | 8.98 | 8.50 | 9.04 | 8.56 | 13.02 | 6.06 | 13.14 | 5.98 | 11.05 | 7.27 | 9.16 |

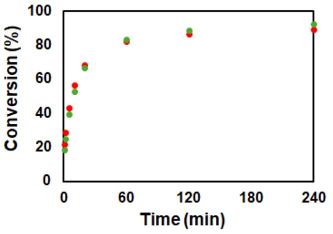

#### 5 μM Clients + 50% PEG400

| | Conversion (%) | | | | | | | | | | |
|---|---|---|---|---|---|---|---|---|---|---|---|
| | Sample 1 - Gel 1 | | Sample 1 - Gel 2 | | Sample 2 - Gel 1 | | Sample 2 - Gel 2 | | Average | | |
| Reaction Time (min) | mCh-ST | GFP-SC | mCh-ST | GFP-SC | mCh-ST | GFP-SC | mCh-ST | GFP-SC | mCh-ST | GFP-SC | Total |
| 1 | 46.07 | 37.10 | 46.43 | 37.66 | 43.05 | 36.94 | 43.69 | 37.14 | 44.81 | 37.21 | 41.01 |
| 2 | 50.49 | 41.48 | 49.90 | 40.48 | 46.99 | 40.67 | 46.71 | 40.14 | 48.52 | 40.69 | 44.61 |
| 5 | 54.21 | 45.18 | 53.60 | 44.41 | 50.16 | 44.37 | 50.03 | 44.28 | 52.00 | 44.56 | 48.28 |
| 10 | 59.21 | 50.43 | 58.59 | 49.94 | 56.67 | 51.09 | 56.08 | 51.02 | 57.64 | 50.62 | 54.13 |
| 20 | 61.62 | 53.86 | 61.67 | 53.33 | 58.81 | 53.69 | 57.86 | 53.15 | 59.99 | 53.51 | 56.75 |
| 60 | 61.80 | 54.33 | 61.41 | 52.97 | 61.20 | 56.81 | 61.06 | 57.49 | 61.37 | 55.40 | 58.38 |
| 120 | 63.98 | 57.09 | 65.56 | 57.14 | 62.40 | 58.04 | 62.30 | 58.41 | 63.56 | 57.67 | 60.62 |
| 240 | 61.73 | 58.80 | 62.06 | 58.64 | 64.31 | 60.80 | 64.73 | 61.21 | 63.21 | 59.86 | 61.54 |
| $k_{obs}$ (M^-1 sec^-1) | 729.02 | 258.74 | 706.44 | 237.01 | 527.93 | 270.90 | 510.26 | 268.84 | 618.41 | 258.88 | 438.64 |
| $k_{adj}$ (M^-1 sec^-1) | 13397.82 | 9638.75 | 12984.48 | 9960.61 | 10259.45 | 8147.98 | 10680.32 | 7875.15 | 11830.52 | 8905.62 | 10368.07 |
| Unreacted Fracrtion (%) | 37.99 | 44.43 | 37.86 | 45.06 | 38.89 | 43.14 | 39.22 | 42.93 | 38.49 | 43.89 | 41.19 |

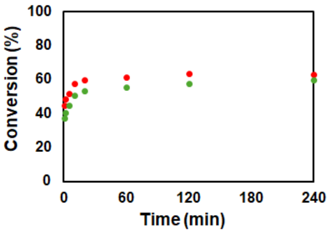

#### 1 μM Clients + 10% PEG400: 136.50 M$^{-1}$ s$^{-1}$

| | Conversion (%) | | | | | | | | | | |
|---|---|---|---|---|---|---|---|---|---|---|---|
| | Sample 1 - Gel 1 | | Sample 1 - Gel 2 | | Sample 2 - Gel 1 | | Sample 2 - Gel 2 | | Average | | |
| Reaction Time (min) | mCh-ST | GFP-SC | mCh-ST | GFP-SC | mCh-ST | GFP-SC | mCh-ST | GFP-SC | mCh-ST | GFP-SC | Total |
| 5 | 3.74 | 3.55 | 3.41 | 3.62 | 3.46 | 3.65 | 3.56 | 4.15 | 3.54 | 3.74 | 3.64 |
| 10 | 6.32 | 5.71 | 6.20 | 5.70 | 5.51 | 5.42 | 5.69 | 6.13 | 5.93 | 5.74 | 5.84 |
| 20 | 11.28 | 9.61 | 10.29 | 8.99 | 10.23 | 9.17 | 10.74 | 9.81 | 10.63 | 9.40 | 10.02 |
| 60 | 27.79 | 24.21 | 27.44 | 23.72 | 23.43 | 21.81 | 23.20 | 22.26 | 25.47 | 23.00 | 24.23 |
| 120 | 41.28 | 37.40 | 40.99 | 36.91 | 37.41 | 33.62 | 37.69 | 34.47 | 39.34 | 35.60 | 37.47 |
| 240 | 55.53 | 51.61 | 55.42 | 50.84 | 52.23 | 49.22 | 52.41 | 49.36 | 53.90 | 50.26 | 52.08 |
| $k_{obs}$ (M^-1 sec^-1) | 96.31 | 80.99 | 94.71 | 78.67 | 81.31 | 71.18 | 81.95 | 73.11 | 88.57 | 75.99 | 82.28 |
| $k_{adj}$ (M^-1 sec^-1) | 160.76 | 138.78 | 146.68 | 136.08 | 125.19 | 118.65 | 127.00 | 138.89 | 139.91 | 133.10 | 136.50 |
| Unreacted Fracrtion (%) | 15.97 | 17.24 | 13.94 | 17.61 | 14.17 | 16.86 | 14.34 | 20.40 | 14.61 | 18.03 | 16.32 |

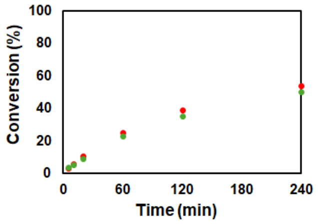

#### 1 μM Clients + 25% PEG400: 550.84 M$^{-1}$ s$^{-1}$

| | Conversion (%) | | | | | | | | | | |
|---|---|---|---|---|---|---|---|---|---|---|---|
| | Sample 1 - Gel 1 | | Sample 1 - Gel 2 | | Sample 2 - Gel 1 | | Sample 2 - Gel 2 | | Average | | |
| Reaction Time (min) | mCh-ST | GFP-SC | mCh-ST | GFP-SC | mCh-ST | GFP-SC | mCh-ST | GFP-SC | mCh-ST | GFP-SC | Total |
| 5 | 9.77 | 8.37 | 10.63 | 8.28 | 14.40 | 12.24 | 15.03 | 13.04 | 12.46 | 10.48 | 11.47 |
| 10 | 17.54 | 15.01 | 17.99 | 15.38 | 23.11 | 19.63 | 23.43 | 20.79 | 20.52 | 17.70 | 19.11 |
| 20 | 28.34 | 24.07 | 29.01 | 24.69 | 35.72 | 30.27 | 36.05 | 31.39 | 32.28 | 27.61 | 29.94 |
| 60 | 51.10 | 46.24 | 51.28 | 46.61 | 57.92 | 53.33 | 58.59 | 53.47 | 54.72 | 49.91 | 52.32 |
| 120 | 63.63 | 59.39 | 65.07 | 60.43 | 68.36 | 66.04 | 67.92 | 65.45 | 66.25 | 62.83 | 64.54 |
| 240 | 73.53 | 71.91 | 74.07 | 71.90 | 76.40 | 77.39 | 74.80 | 76.36 | 74.70 | 74.39 | 74.54 |
| $k_{obs}$ (M^-1 sec^-1) | 273.68 | 223.72 | 284.60 | 229.70 | 380.76 | 315.17 | 381.73 | 318.77 | 330.19 | 271.84 | 301.02 |
| $k_{adj}$ (M^-1 sec^-1) | 489.52 | 376.20 | 499.37 | 390.26 | 742.81 | 508.53 | 815.92 | 584.14 | 636.91 | 464.78 | 550.84 |
| Unreacted Fracrtion (%) | 14.68 | 13.71 | 14.16 | 13.90 | 15.84 | 12.06 | 17.68 | 14.77 | 15.59 | 13.61 | 14.60 |

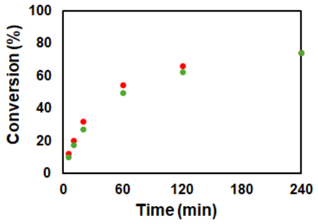

#### 1 µM Clients + 50% PEG400: 1554.18 $M^{-1}$ $s^{-1}$

| | Conversion (%) | | | | | | | | | | |
|---|---|---|---|---|---|---|---|---|---|---|---|
| | Sample 1 - Gel 1 | | Sample 1 - Gel 2 | | Sample 2 - Gel 1 | | Sample 2 - Gel 2 | | Average | | |
| Reaction Time (min) | mCh-ST | GFP-SC | mCh-ST | GFP-SC | mCh-ST | GFP-SC | mCh-ST | GFP-SC | mCh-ST | GFP-SC | Total |
| 5 | 20.74 | 21.02 | 22.41 | 20.56 | 32.22 | 28.67 | 32.41 | 28.20 | 26.94 | 24.61 | 25.78 |
| 10 | 34.45 | 29.03 | 35.40 | 29.64 | 42.74 | 38.87 | 43.16 | 39.36 | 38.94 | 34.22 | 36.58 |
| 20 | 50.23 | 43.14 | 50.45 | 43.24 | 55.56 | 52.10 | 55.86 | 52.13 | 53.03 | 47.65 | 50.34 |
| 60 | 64.83 | 68.39 | 65.27 | 68.61 | 71.53 | 70.79 | 72.30 | 71.08 | 68.48 | 69.72 | 69.10 |
| 120 | 75.60 | 73.83 | 76.44 | 74.86 | 76.53 | 79.46 | 76.80 | 77.90 | 76.34 | 76.51 | 76.43 |
| 240 | 79.74 | 85.61 | 79.47 | 87.40 | 79.84 | 85.01 | 80.05 | 83.44 | 79.77 | 85.37 | 82.57 |
| kobs (M^-1 sec^-1) | 668.80 | 602.15 | 697.07 | 617.83 | 995.12 | 888.54 | 1019.04 | 878.22 | 845.01 | 746.68 | 795.85 |
| kadj (M^-1 sec^-1) | 1369.74 | 902.11 | 1452.17 | 854.97 | 2326.19 | 1545.90 | 2336.00 | 1646.36 | 1871.03 | 1237.33 | 1554.18 |
| Unreacted Fracrtion (%) | 16.14 | 9.74 | 16.39 | 7.94 | 17.99 | 12.54 | 17.64 | 14.06 | 17.04 | 11.07 | 14.06 |

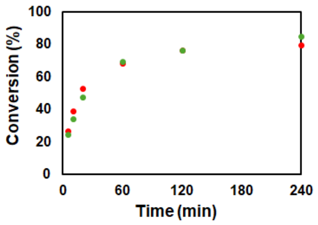

#### 5 µM Clients + 5% glycerol: 90.49 $M^{-1}$ $s^{-1}$

| | Conversion (%) | | | | | | | | | | |
|---|---|---|---|---|---|---|---|---|---|---|---|
| | Sample 1 - Gel 1 | | Sample 1 - Gel 2 | | Sample 2 - Gel 1 | | Sample 2 - Gel 2 | | Average | | |
| Reaction Time (min) | mCh-ST | GFP-SC | mCh-ST | GFP-SC | mCh-ST | GFP-SC | mCh-ST | GFP-SC | mCh-ST | GFP-SC | Total |
| 1 | 3.24 | 3.03 | 2.79 | 3.56 | 2.90 | 2.94 | 3.29 | 2.96 | 3.06 | 3.12 | 3.09 |
| 2 | 4.93 | 4.76 | 5.07 | 5.34 | 5.44 | 5.14 | 5.86 | 5.09 | 5.33 | 5.08 | 5.20 |
| 5 | 12.03 | 10.53 | 12.48 | 10.88 | 12.45 | 10.48 | 12.67 | 10.28 | 12.41 | 10.54 | 11.47 |
| 10 | 21.23 | 18.16 | 20.83 | 18.06 | 22.09 | 18.38 | 22.19 | 17.97 | 21.59 | 18.14 | 19.86 |
| 20 | 33.06 | 28.70 | 33.98 | 29.55 | 34.07 | 28.57 | 34.46 | 28.82 | 33.89 | 28.91 | 31.40 |
| 60 | 59.36 | 54.31 | 60.74 | 56.29 | 61.08 | 53.75 | 60.02 | 52.61 | 60.30 | 54.24 | 57.27 |
| 120 | 72.66 | 68.86 | 73.65 | 70.19 | 73.64 | 67.16 | 74.64 | 68.16 | 73.65 | 68.59 | 71.12 |
| 240 | 83.49 | 81.61 | 83.89 | 81.56 | 82.99 | 77.67 | 83.89 | 78.82 | 83.56 | 79.91 | 81.74 |
| kobs (M^-1 sec^-1) | 80.67 | 66.06 | 83.93 | 69.57 | 84.86 | 62.38 | 85.75 | 62.65 | 83.80 | 65.17 | 74.48 |
| kadj (M^-1 sec^-1) | 94.96 | 75.95 | 96.65 | 80.99 | 103.53 | 89.11 | 100.54 | 82.17 | 98.92 | 82.06 | 90.49 |
| Unreacted Fracrtion (%) | 4.34 | 3.81 | 3.78 | 4.13 | 5.25 | 9.35 | 4.22 | 7.24 | 4.40 | 6.13 | 5.26 |

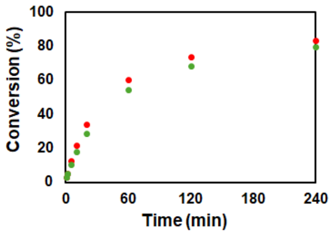

#### 5 µM Clients + 10% glycerol: 118.68 $M^{-1}$ $s^{-1}$

| | Conversion (%) | | | | | | | | | | |
|---|---|---|---|---|---|---|---|---|---|---|---|
| | Sample 1 - Gel 1 | | Sample 1 - Gel 2 | | Sample 2 - Gel 1 | | Sample 2 - Gel 2 | | Average | | |
| Reaction Time (min) | mCh-ST | GFP-SC | mCh-ST | GFP-SC | mCh-ST | GFP-SC | mCh-ST | GFP-SC | mCh-ST | GFP-SC | Total |
| 1 | 3.13 | 3.19 | 3.58 | 3.53 | 3.72 | 3.56 | 3.58 | 4.03 | 3.50 | 3.58 | 3.54 |
| 2 | 6.18 | 5.21 | 6.06 | 5.38 | 6.80 | 5.94 | 7.16 | 6.85 | 6.55 | 5.85 | 6.20 |
| 5 | 15.17 | 12.31 | 15.63 | 13.14 | 16.11 | 13.38 | 16.23 | 14.17 | 15.79 | 13.25 | 14.52 |
| 10 | 25.56 | 21.06 | 26.57 | 21.81 | 26.63 | 22.39 | 27.05 | 22.77 | 26.45 | 22.01 | 24.23 |
| 20 | 40.46 | 33.90 | 40.92 | 34.15 | 41.39 | 35.02 | 41.83 | 35.85 | 41.15 | 34.73 | 37.94 |
| 60 | 65.26 | 57.85 | 65.79 | 58.16 | 66.58 | 59.20 | 66.64 | 60.20 | 66.07 | 58.85 | 62.46 |
| 120 | 77.73 | 72.14 | 78.32 | 72.43 | 78.44 | 73.21 | 78.70 | 73.86 | 78.30 | 72.91 | 75.60 |
| 240 | 86.70 | 82.06 | 86.72 | 82.35 | 87.32 | 83.18 | 87.05 | 83.12 | 86.95 | 82.68 | 84.81 |
| kobs (M^-1 sec^-1) | 107.79 | 78.75 | 111.13 | 80.62 | 113.99 | 84.35 | 115.48 | 87.77 | 112.10 | 82.87 | 97.49 |
| kadj (M^-1 sec^-1) | 124.75 | 101.04 | 129.81 | 104.70 | 131.75 | 107.43 | 135.65 | 114.36 | 130.49 | 106.88 | 118.68 |
| Unreacted Fracrtion (%) | 3.83 | 6.51 | 4.06 | 6.78 | 3.78 | 6.28 | 4.19 | 6.81 | 3.96 | 6.59 | 5.28 |

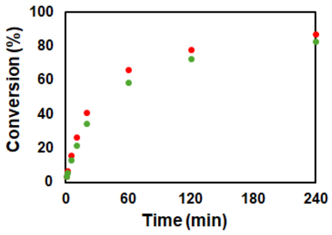

#### 5 µM Clients + 25% glycerol: 275.51 $M^{-1}$ $s^{-1}$

| | Conversion (%) | | | | | | | | | | |
|---|---|---|---|---|---|---|---|---|---|---|---|
| | Sample 1 - Gel 1 | | Sample 1 - Gel 2 | | Sample 2 - Gel 1 | | Sample 2 - Gel 2 | | Average | | |
| Reaction Time (min) | mCh-ST | GFP-SC | mCh-ST | GFP-SC | mCh-ST | GFP-SC | mCh-ST | GFP-SC | mCh-ST | GFP-SC | Total |
| 1 | 8.79 | 7.19 | 8.52 | 7.52 | 9.80 | 8.83 | 9.90 | 8.59 | 9.25 | 8.03 | 8.64 |
| 2 | 14.64 | 11.91 | 14.78 | 12.33 | 14.26 | 12.47 | 14.07 | 12.25 | 14.44 | 12.24 | 13.34 |
| 5 | 30.29 | 24.51 | 29.82 | 24.19 | 28.35 | 25.14 | 27.42 | 24.54 | 28.97 | 24.59 | 26.78 |
| 10 | 45.12 | 37.50 | 43.71 | 36.51 | 42.55 | 38.32 | 41.84 | 37.80 | 43.30 | 37.53 | 40.42 |
| 20 | 60.72 | 51.29 | 59.72 | 50.66 | 56.91 | 53.04 | 56.55 | 52.20 | 58.47 | 51.80 | 55.14 |
| 60 | 81.10 | 72.51 | 79.91 | 71.53 | 77.80 | 75.75 | 77.41 | 74.94 | 79.05 | 73.68 | 76.37 |
| 120 | 87.78 | 78.82 | 87.48 | 78.76 | 84.82 | 83.57 | 84.40 | 83.13 | 86.12 | 81.07 | 83.60 |
| 240 | 92.82 | 85.02 | 93.13 | 85.48 | 90.83 | 89.65 | 90.44 | 89.59 | 91.80 | 87.43 | 89.62 |
| kobs (M^-1 sec^-1) | 265.83 | 171.23 | 254.51 | 166.18 | 230.70 | 193.88 | 223.87 | 187.21 | 243.73 | 179.62 | 211.68 |
| kadj (M^-1 sec^-1) | 305.55 | 273.42 | 292.62 | 263.59 | 295.08 | 246.33 | 288.49 | 238.98 | 295.43 | 255.58 | 275.51 |
| Unreacted Fracrtion (%) | 3.55 | 11.20 | 3.55 | 11.02 | 6.11 | 5.99 | 6.30 | 6.10 | 4.88 | 8.58 | 6.73 |

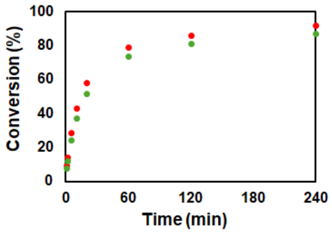

#### 5 µM Clients + 50% glycerol: 415.35 $M^{-1}$ $s^{-1}$

| | Conversion (%) | | | | | | | | | | |
|---|---|---|---|---|---|---|---|---|---|---|---|
| | Sample 1 - Gel 1 | | Sample 1 - Gel 2 | | Sample 2 - Gel 1 | | Sample 2 - Gel 2 | | Average | | |
| Reaction Time (min) | mCh-ST | GFP-SC | mCh-ST | GFP-SC | mCh-ST | GFP-SC | mCh-ST | GFP-SC | mCh-ST | GFP-SC | Total |
| 1 | 11.00 | 9.28 | 10.65 | 8.84 | 10.87 | 8.60 | 10.62 | 8.52 | 10.78 | 8.81 | 9.80 |
| 2 | 18.73 | 15.17 | 17.91 | 14.40 | 19.20 | 15.68 | 18.98 | 15.66 | 18.71 | 15.23 | 16.97 |
| 5 | 36.74 | 29.52 | 36.31 | 29.35 | 37.76 | 31.97 | 37.68 | 32.11 | 37.12 | 30.74 | 33.93 |
| 10 | 53.05 | 44.06 | 53.05 | 44.21 | 52.61 | 45.93 | 53.02 | 46.62 | 52.93 | 45.20 | 49.07 |
| 20 | 68.14 | 58.10 | 67.73 | 57.78 | 65.29 | 59.28 | 66.15 | 60.10 | 66.83 | 58.81 | 62.82 |
| 60 | 84.28 | 74.63 | 84.38 | 74.90 | 80.73 | 76.70 | 82.32 | 77.99 | 82.93 | 76.05 | 79.49 |
| 120 | 88.93 | 79.60 | 88.49 | 79.42 | 86.64 | 83.76 | 87.59 | 83.85 | 87.91 | 81.66 | 84.78 |
| 240 | 93.74 | 85.37 | 93.13 | 84.83 | 90.69 | 88.95 | 90.82 | 88.91 | 92.10 | 87.01 | 89.55 |
| kobs (M^-1 sec^-1) | 364.40 | 226.29 | 357.65 | 223.95 | 344.13 | 254.11 | 352.77 | 262.13 | 354.74 | 241.62 | 298.18 |
| kadj (M^-1 sec^-1) | 429.15 | 389.26 | 425.28 | 386.67 | 476.78 | 371.78 | 466.64 | 377.22 | 449.46 | 381.23 | 415.35 |
| Unreacted Fracrtion (%) | 4.12 | 12.72 | 4.38 | 12.84 | 7.89 | 9.17 | 6.87 | 8.83 | 5.81 | 10.89 | 8.35 |

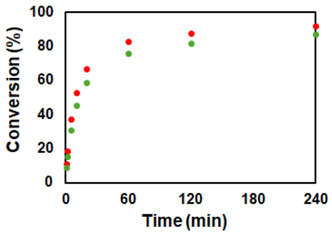

#### - SpyTag003-SpyCatcher003 pair (bulk)

##### 0.25 µM Clients: 13537.60 $M^{-1}$ $s^{-1}$

| | Conversion (%) | | | | | | | | | | |
|---|---|---|---|---|---|---|---|---|---|---|---|
| | Sample 1 - Gel 1 | | Sample 1 - Gel 2 | | Sample 2 - Gel 1 | | Sample 2 - Gel 2 | | Average | | |
| Reaction Time (min) | mCh-ST | GFP-SC | mCh-ST | GFP-SC | mCh-ST | GFP-SC | mCh-ST | GFP-SC | mCh-ST | GFP-SC | Total |
| 1 | 15.45 | 16.42 | 14.96 | 15.17 | 12.05 | 13.31 | 12.37 | 12.93 | 13.71 | 14.46 | 14.08 |
| 2 | 25.58 | 24.20 | 23.87 | 23.55 | 19.94 | 20.61 | 20.47 | 20.47 | 22.47 | 22.21 | 22.34 |
| 5 | 45.49 | 43.85 | 43.94 | 43.29 | 39.22 | 39.52 | 39.56 | 39.23 | 42.05 | 41.47 | 41.76 |
| 10 | 58.31 | 57.49 | 56.35 | 56.59 | 51.15 | 51.81 | 51.14 | 51.72 | 54.24 | 54.40 | 54.32 |
| 20 | 68.16 | 68.52 | 66.56 | 67.74 | 62.49 | 63.46 | 62.37 | 63.60 | 64.89 | 65.83 | 65.36 |
| 60 | 79.79 | 81.26 | 78.15 | 81.37 | 75.14 | 76.85 | 75.47 | 77.43 | 77.14 | 79.23 | 78.18 |
| 120 | 84.45 | 85.32 | 82.65 | 85.62 | 79.91 | 81.63 | 80.24 | 81.87 | 81.81 | 83.61 | 82.71 |
| 240 | 87.59 | 87.67 | 85.97 | 88.52 | 82.63 | 84.93 | 83.43 | 85.00 | 84.91 | 86.53 | 85.72 |
| $k_{obs}$ (M^-1 sec^-1) | 8949.88 | 8729.69 | 8086.60 | 7920.12 | 6179.22 | 6570.25 | 6265.33 | 6564.71 | 7370.26 | 7446.19 | 7408.22 |
| $k_{adj}$ (M^-1 sec^-1) | 15559.18 | 14350.32 | 15136.43 | 12999.03 | 12871.19 | 12422.17 | 12827.06 | 12135.43 | 14098.47 | 12976.74 | 13537.60 |
| Unreacted Fracrtion (%) | 12.51 | 11.40 | 14.04 | 10.48 | 16.31 | 14.35 | 15.91 | 13.94 | 14.69 | 12.54 | 13.62 |

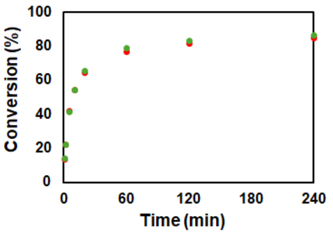

##### 0.5 µM Clients: 13537.48 $M^{-1}$ $s^{-1}$

| | Conversion (%) | | | | | | | | | | |
|---|---|---|---|---|---|---|---|---|---|---|---|
| | Sample 1 - Gel 1 | | Sample 1 - Gel 2 | | Sample 2 - Gel 1 | | Sample 2 - Gel 2 | | Average | | |
| Reaction Time (min) | mCh-ST | GFP-SC | mCh-ST | GFP-SC | mCh-ST | GFP-SC | mCh-ST | GFP-SC | mCh-ST | GFP-SC | Total |
| 1 | 25.97 | 22.44 | 27.65 | 23.35 | 22.46 | 21.74 | 22.26 | 21.38 | 24.59 | 22.23 | 23.41 |
| 2 | 41.86 | 36.70 | 42.53 | 36.34 | 39.23 | 38.24 | 39.51 | 38.61 | 40.78 | 37.47 | 39.13 |
| 5 | 62.41 | 56.32 | 63.03 | 56.63 | 58.44 | 59.95 | 57.86 | 59.65 | 60.44 | 58.14 | 59.29 |
| 10 | 72.53 | 67.76 | 73.28 | 67.83 | 67.61 | 71.10 | 67.43 | 71.06 | 70.21 | 69.44 | 69.83 |
| 20 | 81.61 | 78.16 | 82.23 | 78.30 | 74.71 | 80.95 | 74.92 | 81.20 | 78.37 | 79.65 | 79.01 |
| 60 | 88.82 | 85.45 | 88.88 | 85.40 | 81.26 | 88.47 | 81.10 | 88.43 | 85.02 | 86.94 | 85.98 |
| 120 | 90.57 | 87.51 | 90.91 | 87.54 | 84.48 | 90.67 | 84.44 | 90.95 | 87.60 | 89.17 | 88.38 |
| 240 | 93.67 | 89.09 | 93.96 | 89.44 | 86.40 | 92.37 | 86.24 | 92.45 | 90.07 | 90.83 | 90.45 |
| $k_{obs}$ (M^-1 sec^-1) | 10340.41 | 7857.87 | 10831.04 | 7006.91 | 7924.77 | 9002.17 | 7862.29 | 8995.35 | 9239.63 | 8215.57 | 8727.60 |
| $k_{adj}$ (M^-1 sec^-1) | 14169.30 | 12437.20 | 14796.11 | 12137.71 | 15338.94 | 12113.97 | 15284.95 | 12021.62 | 14897.33 | 12177.62 | 13537.48 |
| Unreacted Fracrtion (%) | 7.14 | 10.31 | 7.03 | 10.02 | 14.20 | 6.91 | 14.28 | 6.76 | 10.66 | 8.50 | 9.58 |

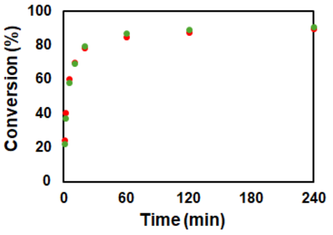

##### 1 µM Clients: 12450.07 $M^{-1}$ $s^{-1}$

| | Conversion (%) | | | | | | | | | | |
|---|---|---|---|---|---|---|---|---|---|---|---|
| | Sample 1 - Gel 1 | | Sample 1 - Gel 2 | | Sample 2 - Gel 1 | | Sample 2 - Gel 2 | | Average | | |
| Reaction Time (min) | mCh-ST | GFP-SC | mCh-ST | GFP-SC | mCh-ST | GFP-SC | mCh-ST | GFP-SC | mCh-ST | GFP-SC | Total |
| 1 | 38.70 | 34.10 | 39.21 | 34.51 | 37.84 | 34.72 | 37.43 | 34.89 | 38.30 | 34.55 | 36.42 |
| 2 | 53.71 | 52.19 | 53.71 | 51.83 | 54.53 | 53.44 | 53.99 | 53.25 | 53.98 | 52.68 | 53.33 |
| 5 | 71.24 | 71.31 | 71.10 | 71.26 | 72.83 | 73.68 | 72.22 | 73.98 | 71.85 | 72.56 | 72.20 |
| 10 | 79.61 | 80.62 | 79.52 | 80.49 | 80.15 | 81.98 | 79.54 | 82.23 | 79.71 | 81.33 | 80.52 |
| 20 | 82.08 | 83.72 | 81.85 | 83.64 | 84.49 | 87.77 | 84.10 | 87.90 | 83.13 | 85.76 | 84.44 |
| 60 | 88.17 | 89.50 | 88.11 | 89.56 | 88.36 | 91.36 | 88.28 | 91.33 | 88.23 | 90.44 | 89.33 |
| 120 | 89.63 | 91.61 | 89.75 | 91.24 | 90.43 | 92.56 | 90.10 | 92.77 | 89.98 | 92.05 | 91.01 |
| 240 | 91.44 | 93.12 | 91.98 | 92.56 | 92.57 | 93.93 | 91.89 | 94.23 | 91.97 | 93.46 | 92.72 |
| $k_{obs}$ (M^-1 sec^-1) | 8521.28 | 8014.74 | 8553.35 | 6776.74 | 8852.57 | 8762.36 | 6802.62 | 8811.83 | 8182.45 | 8091.42 | 8136.94 |
| $k_{adj}$ (M^-1 sec^-1) | 13593.69 | 11203.71 | 13648.34 | 12087.71 | 13224.25 | 11225.44 | 13429.81 | 11187.64 | 13474.02 | 11426.13 | 12450.07 |
| Unreacted Fracrtion (%) | 9.71 | 7.33 | 9.68 | 7.95 | 8.50 | 5.52 | 9.02 | 5.33 | 9.23 | 6.53 | 7.88 |

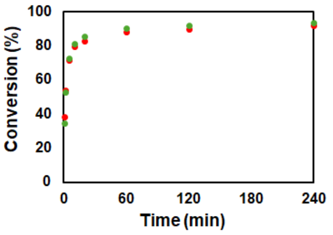

##### 2 µM Clients: 10901.41 $M^{-1}$ $s^{-1}$

| | Conversion (%) | | | | | | | | | | |
|---|---|---|---|---|---|---|---|---|---|---|---|
| | Sample 1 - Gel 1 | | Sample 1 - Gel 2 | | Sample 2 - Gel 1 | | Sample 2 - Gel 2 | | Average | | |
| Reaction Time (min) | mCh-ST | GFP-SC | mCh-ST | GFP-SC | mCh-ST | GFP-SC | mCh-ST | GFP-SC | mCh-ST | GFP-SC | Total |
| 1 | 52.30 | 50.14 | 51.99 | 49.38 | 47.51 | 46.06 | 48.46 | 47.05 | 50.06 | 48.16 | 49.11 |
| 2 | 63.61 | 65.10 | 64.05 | 64.96 | 64.52 | 67.13 | 64.37 | 67.51 | 64.14 | 66.17 | 65.16 |
| 5 | 78.58 | 82.00 | 77.47 | 81.89 | 77.22 | 82.98 | 76.96 | 83.14 | 77.56 | 82.50 | 80.03 |
| 10 | 83.39 | 87.90 | 82.76 | 88.00 | 81.61 | 88.10 | 81.89 | 88.31 | 82.41 | 88.08 | 85.25 |
| 20 | 85.68 | 90.39 | 85.13 | 90.63 | 83.53 | 90.47 | 83.83 | 90.24 | 84.54 | 90.43 | 87.49 |
| 60 | 88.90 | 92.67 | 88.57 | 92.77 | 86.01 | 92.74 | 86.34 | 92.65 | 87.45 | 92.71 | 90.08 |
| 120 | 89.86 | 93.58 | 89.75 | 93.59 | 87.85 | 93.83 | 88.18 | 94.25 | 88.91 | 93.81 | 91.36 |
| 240 | 91.90 | 95.59 | 91.84 | 95.51 | 90.44 | 95.85 | 89.97 | 95.85 | 91.04 | 95.70 | 93.37 |
| $k_{obs}$ (M^-1 sec^-1) | 7099.28 | 7677.95 | 6975.14 | 6397.09 | 6350.77 | 7476.08 | 6443.78 | 7667.51 | 6717.24 | 7304.66 | 7010.95 |
| $k_{adj}$ (M^-1 sec^-1) | 11918.48 | 9878.51 | 11987.10 | 11270.96 | 11583.27 | 9254.86 | 11790.31 | 9527.78 | 11819.79 | 9983.03 | 10901.41 |
| Unreacted Fracrtion (%) | 9.67 | 5.02 | 10.06 | 5.38 | 11.42 | 4.39 | 11.40 | 4.43 | 10.63 | 4.80 | 7.72 |

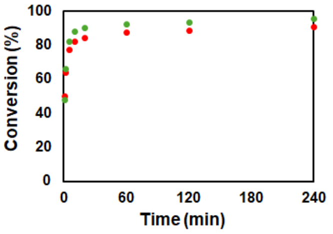

##### 0.1µM Clients (low concentration)

| | Conversion (%) | | | | | | | | | | |
|---|---|---|---|---|---|---|---|---|---|---|---|
| | Sample 1 - Gel 1 | | Sample 1 - Gel 2 | | Sample 2 - Gel 1 | | Sample 2 - Gel 2 | | Average | | |
| Reaction Time (min) | mCh-ST | GFP-SC | mCh-ST | GFP-SC | mCh-ST | GFP-SC | mCh-ST | GFP-SC | mCh-ST | GFP-SC | Total |
| 1 | 7.00 | 8.92 | 8.09 | 9.00 | 6.08 | 7.38 | 5.71 | 6.99 | 6.72 | 8.07 | 7.40 |
| 2 | 12.72 | 13.81 | 13.38 | 14.07 | 10.43 | 11.52 | 10.18 | 11.35 | 11.68 | 12.69 | 12.18 |
| 5 | 21.92 | 24.82 | 22.93 | 25.48 | 22.12 | 22.57 | 21.93 | 22.16 | 22.22 | 23.76 | 22.99 |
| 10 | 32.06 | 35.64 | 33.13 | 36.79 | 31.79 | 31.96 | 31.79 | 31.48 | 32.19 | 33.97 | 33.08 |
| 20 | 44.20 | 48.33 | 45.14 | 48.94 | 42.16 | 42.30 | 42.15 | 42.13 | 43.41 | 45.43 | 44.42 |
| 60 | 57.24 | 64.25 | 58.42 | 64.78 | 60.36 | 61.93 | 59.87 | 61.99 | 58.97 | 63.24 | 61.10 |
| 120 | 64.83 | 69.53 | 64.81 | 70.25 | 65.33 | 65.61 | 64.49 | 65.32 | 64.87 | 67.68 | 66.27 |
| 240 | 64.76 | 60.03 | 63.96 | 62.05 | 70.20 | 68.80 | 70.42 | 70.18 | 67.34 | 65.27 | 66.30 |
| $k_{obs}$ (M^-1 sec^-1) | 4405.75 | 7108.45 | 4658.54 | 7131.64 | 4674.47 | 4826.09 | 4582.76 | 4803.08 | 4580.38 | 5967.31 | 5273.85 |
| $k_{adj}$ (M^-1 sec^-1) | 22865.09 | 21405.46 | 25316.95 | 20694.69 | 17701.21 | 18970.64 | 17696.20 | 17807.53 | 20894.86 | 19719.58 | 20307.22 |
| Unreacted Fracrtion (%) | 31.58 | 24.89 | 32.13 | 23.84 | 27.05 | 27.64 | 27.34 | 26.74 | 29.53 | 25.78 | 27.65 |

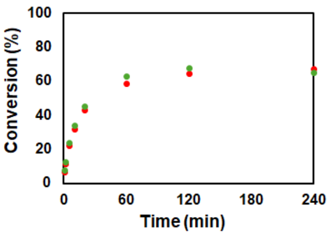

#### - SpyTag003-SpyCatcher003 pair (condensates)

##### PRM-SH3short (100 µM; 5% PUMA), 0.01 µM mCh-ST003-Bcl: 3284.99 $M^{-1}$ $s^{-1}$

| Client Concentration = 0.745 µM | Conversion (%) | | | | | | | | | | |
|---|---|---|---|---|---|---|---|---|---|---|---|
| GFP client ratio = 0.68 [mCh] | Sample 1 - Gel 1 | | Sample 1 - Gel 2 | | Sample 2 - Gel 1 | | Sample 2 - Gel 2 | | Average | | |
| Reaction Time (min) | mCh-ST | GFP-SC | mCh-ST | GFP-SC | mCh-ST | GFP-SC | mCh-ST | GFP-SC | mCh-ST | GFP-SC | Total |
| 5 | 35.35 | 34.53 | 33.92 | 33.85 | 34.02 | 32.31 | 37.74 | 33.94 | 35.26 | 33.66 | 34.46 |
| 10 | 43.59 | 41.21 | 47.92 | 43.14 | 45.75 | 40.23 | 48.34 | 42.49 | 46.40 | 41.77 | 44.08 |
| 20 | 55.18 | 50.51 | 55.31 | 49.80 | 58.51 | 48.58 | 56.95 | 50.79 | 56.49 | 49.92 | 53.20 |
| 60 | 70.70 | 66.99 | 70.38 | 65.59 | 69.97 | 65.86 | 72.16 | 66.72 | 70.80 | 66.29 | 68.55 |
| 120 | 80.77 | 72.99 | 79.93 | 72.28 | 81.20 | 70.85 | 83.32 | 72.15 | 81.30 | 72.07 | 76.69 |
| 240 | 86.66 | 76.97 | 88.14 | 76.82 | 85.83 | 73.42 | 86.37 | 79.18 | 86.75 | 76.60 | 81.67 |
| $k_{obs}$ (M^-1 sec^-1) | 1456.47 | 1122.63 | 1526.38 | 1105.66 | 1546.47 | 990.50 | 1702.41 | 1143.40 | 1557.93 | 1090.55 | 1324.24 |
| $k_{adj}$ (M^-1 sec^-1) | 2754.01 | 3542.14 | 2913.78 | 3736.28 | 2997.96 | 3602.61 | 3246.40 | 3486.74 | 2978.04 | 3591.94 | 3284.99 |
| Unreacted Fracrtion (%) | 13.74 | 22.72 | 13.86 | 23.69 | 14.27 | 25.14 | 13.69 | 22.11 | 13.89 | 23.41 | 18.65 |

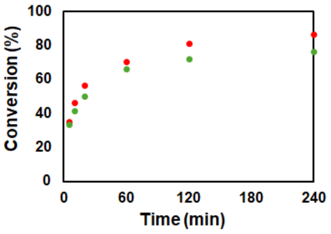

##### PRM-SH3long (100 µM; 5% PUMA), 0.01 µM mCh-ST003-Bcl: 8998.79 $M^{-1}$ $s^{-1}$

| Client Concentration = 0.372 µM | Conversion (%) | | | | | | | | | | |
|---|---|---|---|---|---|---|---|---|---|---|---|
| GFP client ratio = 0.75 [mCh] | Sample 1 - Gel 1 | | Sample 1 - Gel 2 | | Sample 2 - Gel 1 | | Sample 2 - Gel 2 | | Average | | |
| Reaction Time (min) | mCh-ST | GFP-SC | mCh-ST | GFP-SC | mCh-ST | GFP-SC | mCh-ST | GFP-SC | mCh-ST | GFP-SC | Total |
| 5 | 45.53 | 40.70 | 44.78 | 41.22 | 42.51 | 40.58 | 42.77 | 38.90 | 43.90 | 40.35 | 42.12 |
| 10 | 62.07 | 58.04 | 61.11 | 57.98 | 61.62 | 57.74 | 59.53 | 57.42 | 61.08 | 57.79 | 59.44 |
| 20 | 71.63 | 68.34 | 72.76 | 68.20 | 72.00 | 69.38 | 70.27 | 68.68 | 71.67 | 68.65 | 70.16 |
| 60 | 83.80 | 80.99 | 84.26 | 81.21 | 83.44 | 81.84 | 81.97 | 81.53 | 83.37 | 81.39 | 82.38 |
| 120 | 87.17 | 86.10 | 86.62 | 85.78 | 87.16 | 86.03 | 86.36 | 86.07 | 86.83 | 85.99 | 86.41 |
| 240 | 89.87 | 86.34 | 90.71 | 86.49 | 89.66 | 86.28 | 90.19 | 87.07 | 90.10 | 86.54 | 88.32 |
| $k_{obs}$ (M^-1 sec^-1) | 6457.11 | 5255.42 | 6389.95 | 5284.14 | 6131.50 | 5321.65 | 5768.98 | 5123.34 | 6186.89 | 5246.14 | 5716.51 |
| $k_{adj}$ (M^-1 sec^-1) | 9963.20 | 8762.88 | 9526.37 | 8870.89 | 9169.47 | 8703.59 | 8880.68 | 8113.29 | 9384.93 | 8612.66 | 8998.79 |
| Unreacted Fracrtion (%) | 9.04 | 10.93 | 8.44 | 11.02 | 8.62 | 10.58 | 9.20 | 10.04 | 8.83 | 10.64 | 9.73 |

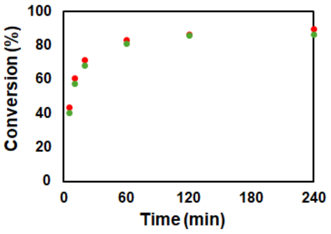

##### LAF (50 µM; 5% PUMA), 0.01 µM mCh-ST003-Bcl: Too fast to measure

| Client Concentration = 0.231 µM | Conversion (%) | | | | | | | | | | |
|---|---|---|---|---|---|---|---|---|---|---|---|
| GFP client ratio = 0.92 [mCh] | Sample 1 - Gel 1 | | Sample 1 - Gel 2 | | Sample 2 - Gel 1 | | Sample 2 - Gel 2 | | Average | | |
| Reaction Time (min) | mCh-ST | GFP-SC | mCh-ST | GFP-SC | mCh-ST | GFP-SC | mCh-ST | GFP-SC | mCh-ST | GFP-SC | Total |
| 5 | 78.27 | 88.54 | 71.31 | 93.58 | 72.76 | 86.28 | 74.14 | 91.51 | 74.12 | 89.98 | 82.05 |
| 10 | 80.80 | 91.72 | 79.46 | 94.96 | 77.29 | 89.86 | 79.69 | 92.14 | 79.31 | 92.17 | 85.74 |
| 20 | 83.06 | 88.26 | 86.40 | 94.42 | 83.49 | 90.81 | 85.41 | 93.28 | 84.59 | 91.69 | 88.14 |
| 60 | 87.21 | 90.45 | 88.21 | 93.89 | 83.54 | 92.58 | 84.43 | 94.95 | 85.85 | 92.97 | 89.41 |
| 120 | 91.01 | 92.90 | 88.86 | 96.28 | 86.82 | 91.96 | 87.84 | 93.40 | 88.63 | 93.64 | 91.13 |
| 240 | 91.37 | 93.29 | 93.22 | 96.64 | 84.85 | 91.20 | 90.53 | 95.32 | 89.99 | 94.11 | 92.05 |
| $k_{obs}$ (M^-1 sec^-1) | 35597 | 79944 | 29432 | 155941 | 27246 | 70453 | 30894 | 111837 | 30792 | 104544 | 67668 |

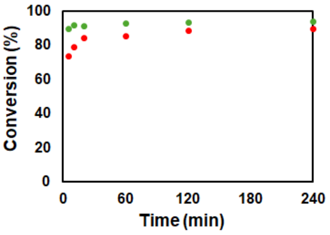
